# Leveraging Chlorination-Based Mechanism for Resolving Subcellular Hypochlorous Acid

**DOI:** 10.1101/2024.08.22.609247

**Authors:** Fung Kit Tang, Lawrence Tucker, Maheshwara Reddy Nadiveedhi, Colby Hladun, Jared Morse, Mahnoor Ali, Noah Payne, Matthias Schmidt, Kaho Leung

## Abstract

Hypochlorous acid (HOCl) is crucial for pathogen defense, but an imbalance in HOCl levels can lead to tissue damage and inflammation. Existing HOCl indicators employ an oxidation approach, which may not truly reveal the chlorinative stress environment. We designed a suite of indicators with a new chlorination-based mechanism, termed HOClSense dyes, to resolve HOCl in sub-cellular compartments. HOClSense dyes allow the visualization of HOCl with both switch-on and switch-off detection modes with diverse emission colors, as well as a unique redshift in emission. HOClSense features a minimalistic design with impressive sensing performance in terms of HOCl selectivity, and our design also facilitates functionalization through click chemistry for resolving subcellular HOCl. As a proof of concept, we targeted plasma membrane and lysosomes with HOClSense for subcellular HOCl mapping. With utilizing HOClSense, we discovered the STING pathway-induced HOCl production and the abnormal HOCl production in Niemann-Pick diseases. To the best of our knowledge, this is the first chlorination-based HOCl indicator series for resolving subcellular HOCl.

## Introduction

Hypochlorite (HOCl) is a potent reactive oxygen species (ROS) that plays a critical role in the immune response to defend against pathogens. When bacteria invade cells, neutrophils detect the foreign material, elicit a downstream immune response, then capture and break down these materials via NETosis (extracellular) and phagocytosis (intracellular).^1^ HOCl is the key component in killing pathogens as it is membrane permeable.^2^ It originates from myeloperoxidase (MPO) in lysosomes, which rapidly catalyzes H_2_O_2_ and Cl^−^ to produce HOCl.^3^ During respiratory burst, HOCl is generated to eliminate pathogens through the oxidative and chlorinative mechanisms. The imbalance of HOCl causes oxidative and chlorinative damage to host tissues, responsible for tissue injury and inflammatory diseases, such as rheumatoid arthritis,^4^ cancer,^5^ and atherosclerosis.^6^

To detect and visualize HOCl, existing HOCl indicators have predominantly relied on modulating fluorescence intensity via the oxidation of specific fluorophore motifs, such as pyrroles,^7,8^ thiols,^9-11^ oximes/imines,^12-14^ catechols,^15,16^ chalcogens^17-19^ and oxidative bond cleavages.^20-25^ In addition to its oxidative properties, HOCl plays a crucial role in chlorinating biomolecules such as amino acids, nucleic acids, lipids, and carbohydrates.^26-31^ These chlorination reactions contribute to a form of cellular stress known as chlorinative stress, which can significantly disrupt cellular signaling pathways. This disruption influences critical processes such as cell survival and apoptosis, and modulates immune responses, thereby linking chlorinative stress to inflammation^32-34^ and age-related diseases.^35^ This highlights the importance of studying chlorinative stress to better understand its impact on cellular function and disease progression. Existing HOCl indicators employ an oxidation approach that may not truly reveal the chlorinative stress environment.^31,35,36^ A chlorination-based HOCl indicator capable of “capturing” chlorine from HOCl would be ideal for detecting HOCl and chlorinative stress. It also minimizes potential interference from other ROS or reactive nitrogen species (RNS). However, chlorination-based HOCl detection is challenging because halides are generally known to quench fluorophores,^37,38^ and non-selective chlorination often lead to fluorescence quenching.^39^

HOCl is a weak acid with a pKa of 7.6, meaning that its equilibrium with hypochlorite ions (OCl^-^) depends on the pH. At pH values below 7.6, the stronger oxidant HOCl predominates, whereas above 7.6, OCl^-^ is predominant, indicating that pH determines which form of HOCl is more prevalent and consequently influences its impact on biomolecules and cells.^40^ On the other hand, cell compartmentalization allows different organelles to maintain distinct internal pH environments, enabling them to carry out specialized functions. Therefore, a HOCl indicator that can selectively label specific cellular compartment for subcellular HOCl imaging is crucial to study the chlorinative stress. However, the cellular distribution of existing HOCl indicators mainly relies on their intrinsic property of the molecules, which might require a lengthy synthetic modification to change the subcellular localization of the HOCl indicators. Furthermore, multi-spectral imaging is often employed in chemical biology where sensors with different emission options are preferred but the existing HOCl indicators lack diverse emission options and fine-tuning capabilities, which limits their practical use in imaging studies. Hence, there is a pressing need for HOCl indicators that can detect HOCl via chlorination, offer emission at different wavelengths, and be functionalized for subcellular imaging to advance studies of HOCl.

This work presents the design and synthesis of a novel series of HOCl indicators, termed HOClSense, which detect HOCl through a unique mechanism involving the chlorination of fluorophores. HOClSense indicators exhibit high selectivity and responsiveness to HOCl, enabling both switch-on and switch-off modes of detection, accompanied by a fluorescence emission redshift. This innovative mechanism confers HOClSense with remarkable versatility, allows tuning of emission properties for multispectral imaging and facilitates functionalization for organelle-selective HOCl imaging. The effectiveness of HOClSense is further demonstrated by its facile synthesis, efficient subcellular targeting, and robust performance across a range of biological applications. The structure-activity relationship study provides critical insights into the detection mechanism, while the ability of HOClSense dyes to monitor HOCl levels has been validated through in-cell HOCl calibration and the assessment of LPS-stimulated HOCl production. HOClSense was also used to investigate the hyper-production of HOCl following PAMPs stimulation in pharmacologically induced NP-A/B and NP-C cell models. Additionally, we observed elevated basal HOCl production in primary cells derived from NP-C patients. These discoveries demonstrate the immediate practical relevance of this tool to the research community for studying chlorinative stress.

## Results

### Novel HOCl-detecting mechanism of HOClSense detects HOCl through chlorination

BODIPY emerges as a promising dye for cellular imaging due to its notable high brightness and cellular permeability.^41^ The photophysical properties of BODIPY with various modifications have been thoroughly investigated.^42-44^ It was reported that BODIPY dyes free of substituents at the 2,6-positions can undergo electrophilic substitution reactions in the presence of chlorosulfonic acid, while other electrophiles can be introduced in an analogous manner for further synthetic modification.^45^ For the synthesis of the chlorinated BODIPY, trichloroisocyanuric acid was used for BODIPY chlorination.^46,47^ Trichloroisocyanuric acid serves as a disinfectant, algicide, and bactericide, offering versatility and efficiency in chlorination and oxidation reactions. Similarly, HOCl acts as a primary disinfectant agent and a powerful ROS, inducing both oxidation and chlorination reactions. Furthermore, the quantum yield and the maximum emission wavelength of BODIPY has been previously observed to change upon 2,6-dichlorination.^46^ We hypothesized that HOCl has the potential to chlorinate BODIPY dye, leading to a red-shift in its fluorescence emission spectrum and change in its quantum yield. In other words, the BODIPY core that free of substituents at the 2,6-positions could function as chlorination-based HOCl-sensing fluorophores.

Firstly, **1a** was synthesized^46,48^ and its ability to detect HOCl was investigated. Upon addition of sodium hypochlorite (NaOCl), which served as the source of HOCl, **1a** exhibited a red-shifted emission in a physiological buffer with its emission wavelength changing from 512 nm to 546 nm (Fig. 1b, Supplementary Figure 40). The fluorescence intensity of **1a** decreased significantly in the presence of HOCl, making **1a** a “Turn-Off” probe for HOCl detection. The UV-Vis absorption spectra also showed the shift in absorption peak maximum from 510 nm to 540 nm, suggesting **1a** reacted with HOCl and formed a new compound with different absorption, causing the red-shift change in excitation and emission (Supplementary Figure 40). The results of high-resolution mass spectroscopy (HRMS) further validate the hypothesis that HOCl can chlorinate **1a** (Supplementary Figure 41). Notably, **1a** can detect HOCl in a pH-independent manner, as evidenced by the fluorescence intensity of **1a** across a pH range of 4 to 8. (Supplementary Figure 42).

**Figure 1.**
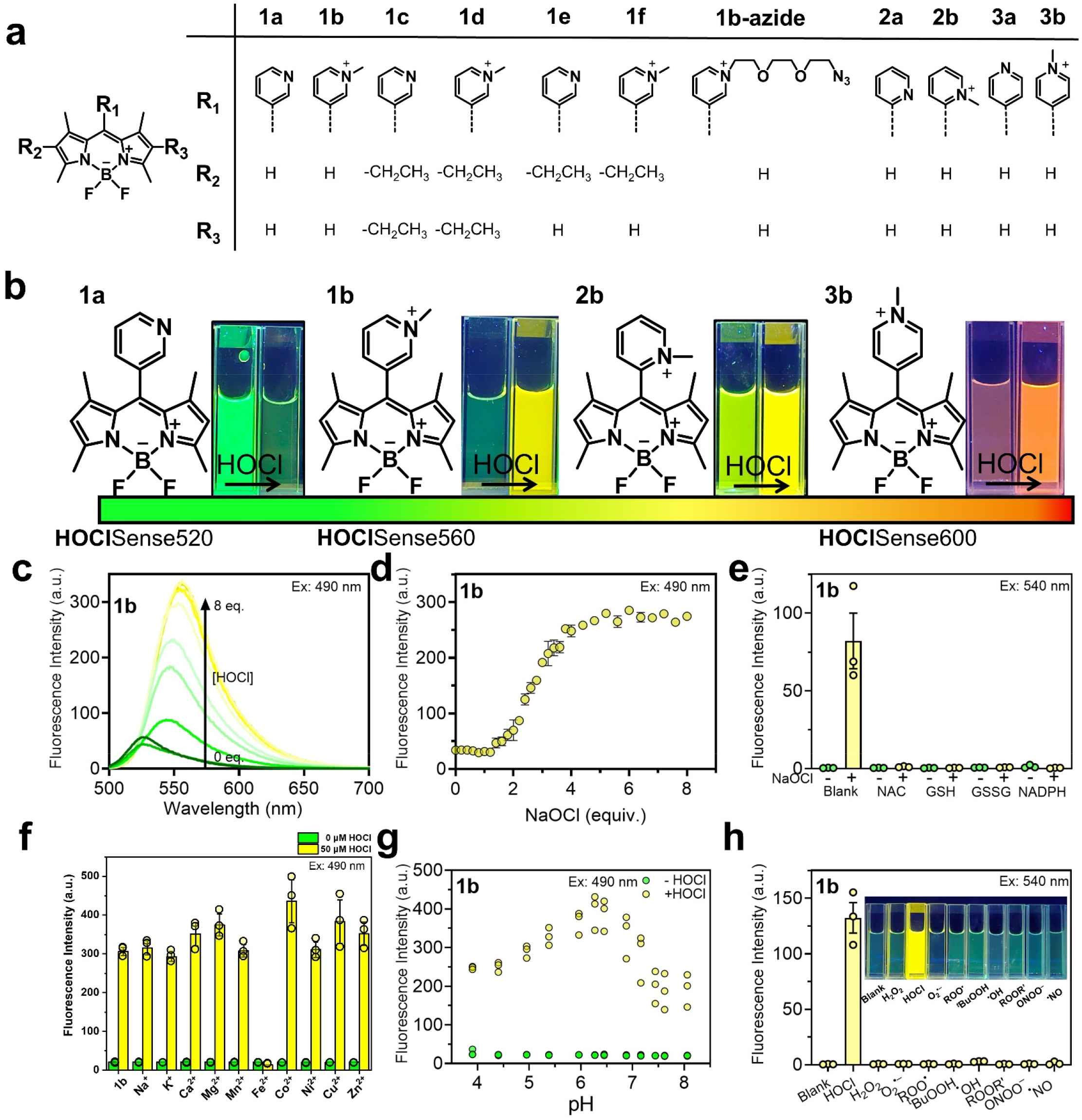
HOClSense dyes, a series of selective HOCl indicators with diverse emission options and detection mode. **a**, Molecular structure of HOClSense dyes. **b**, Multispectral fluorescent response of HOClSense dyes (**1a, 1b, 2b**, and **3b**) to HOCl. **c**, Emission spectra of **1b** (5 µM) in the presence of HOCl (0-8 equiv.). Conditions: water containing 0.5% DMF, λ_ex_ = 490 nm. **d**, Fluorescence intensity of **1b** (5 µM) in the presence of HOCl (0-8 equiv.). Conditions: water containing 0.5% DMF, λ_ex_ = 490 nm, λ_em_ = 560 nm. **e**, Fluorescence intensity at 560 nm of **1b** (5 µM) in the presence of various ROS scavengers (300 µM *N*-Acetyl Cysteine, 1 mM GSH, 1mM GSSG and 100 µM NADPH) with and without HOCl (10 equiv.). Conditions: water containing 0.5% DMF, λ_ex_ = 540 nm, λ_em_ = 560 nm. **f**, Fluorescence intensity of **1b** (5 µM) in the presence of different metal ion (20 µM for Mn^2+^, Fe^2+^, Co^2+^, Ni^2+^, Cu^2+^, Zn^2+^, and 100 mM for Na^+^, K^+^, Ca^2+^, Mg^2+^) with and without HOCl (10 equiv.). Conditions: water containing 0.5% DMF. λ_ex_ = 490 nm, λ_em_ = 560 nm. **g**, Fluorescence intensity of **1b** (5 µM) at 560 nm in the presence and absence of HOCl (10 equiv.) at different pH. Conditions: citrate-phosphate buffer (10 mM) containing 0.5 % DMF, λ_ex_ = 490 nm, λ_em_ = 560 nm, n = 3. **h**, Fluorescence intensity of **1b** (5 µM) in the presence of different ROS/ RNS (10 equiv.) with and without HOCl (10 equiv.). Photo taken under UV light. Conditions: phosphate buffer (60 mM, pH 7.2) containing 0.5% DMF, λ_ex_ = 540 nm, λ_em_ = 560 nm. Error bars indicate the mean ± standard error of mean (s.e.m.) of three independent measurements.

To design a HOCl-sensing fluorophore with a fluorescence “Turn-On” signal, this was proceeded by methylating **1a** to form **1b**. It has been reported that the robust fluorescence characteristic of pyridyl-BODIPY, such as **1a**, is suppressed upon protonation or coordination of the pyridine nitrogen atom.^49-51^ Furthermore, the fluorescence intensity of *N*-methylpyridinium BODIPY is significantly quenched due to the photoinduced electron transfer.^52^ Consequently, we hypothesized that **1b** would initially exhibit weak fluorescence and become bright upon chlorination by HOCl. As anticipated, **1b** exhibited weak fluorescence, whereas **1b** demonstrated robust fluorescence with an emission red-shift from 525 nm to 555 nm upon addition of HOCl (Fig. 1b-d, Supplementary Figure 40). Remarkably, the “Turn-On” fluorescence response of **1b**, characterized by a red-shift emission, manifested a higher fold-change (170-fold) when excited at 540 nm as opposed to 490 nm (Supplementary Figure 44).

To develop HOCl-sensing fluorophores with distinct emission wavelengths, *N*-methylated(pyridyl)-BODIPY compounds were synthesized with different pyridyl groups at the 8-position of the BODIPY chromophore (**2a**-**3b**, Fig. 1a). Both **2a** and **3a** exhibited an emission of around 515 nm and a “Turn-Off” response towards HOCl, which is similar to that of **1a** (Supplementary Figure 40). Upon addition of HOCl, the emissions of **2a** and **3a** decreased significantly and red-shifted. **2b** displayed an emission shift from 537 nm to 555 nm with a reasonable increase in fluorescence. Notably, **3b** displayed an emission at 594 nm, which shifted to 600 nm with enhanced emission in the presence of HOCl (Fig. 1b, Supplementary Figure 40). In other words, by replacing the pyridyl group at the 8-position of the BODIPY chromophore from *meso*-(*N*-methyl-3-pyridyl)-BODIPY (**2b**) to *meso*-(*N*-methyl-4-pyridyl)-BODIPY (**3b**), the emission of the HOCl-sensing fluorophore was tuned to the orange emission range. These results demonstrate that the novel HOCl detection mechanism not only enables both switch-off and switch-on modes of detection but also offers diverse emission options suitable for multispectral imaging, with the emission shift response further enhancing the signal-to-noise ratio for improved detection sensitivity.

### *in vitro* characterization of HOCl-sensing fluorophore

Among all the dyes evaluated, **1b** showed the highest emission change with the bathochromic shift of around 30 nm upon reaction with HOCl in the testing conditions, compared to **2b** with around 18 nm emission red-shift and **3b** with *ca*. 6 nm emission red-shift. **1b** also showed fast kinetic response to HOCl, responding within minutes (Supplementary Figure 43, video S1). Hence, **1b** was chosen for further characterization due to the favorable fluorescent change. As seen in the fluorescence intensity of the titration curve (Fig. 1c-d, Supplementary Figure 44), **1b** showed gradual fluorescence enhancement with an excitation of 490 nm. To confirm that the emission enhancement of **1b** is triggered by HOCl, ROS scavengers were added. The emission enhancement of **1b** in the presence of HOCl was significantly reduced when *N*-acetyl cysteine, GSH, GSSG, and NADPH were introduced, as the majority of the highly oxidizing HOCl reacted with these biologically relevant reductants (Fig. 1e, Supplementary Figure 45). The impact of physiological ions on **1b** was also investigated. The HOCl detecting ability of **1b** was not affected by the presence of physiological ions (Na^+^, K^+^, Ca^2+^, Mg^2+^) and trace ions (Zn^2+^, Mn^2+^, and Cu^2+^), but was greatly reduced in the presence of ferrous ions (Fe^2+^), which was possibly due to the Fenton type redox reactions (Fig. 1f, Supplementary Figure 46).^53,54^ **1b** recognized HOCl and exhibited a high fluorescence in physiological pH range 4.0-7.0, which is useful for HOCl detection in different cellular organelles (Fig. 1g, Supplementary Figure 47).

There are numerous types of ROS produced in cells. ROS/RNS are known for their high reactivity with short lifetimes^55^ which could cause quenching of fluorophores or no response. Therefore, a selective HOCl indicator is important to study HOCl-involved biological processes. The peak-to-peak emission was enhanced by about 150-fold in the presence of HOCl upon 540 nm excitation while the other ROS/RNS (including H_2_O_2_, O_2_**•-**, ^*t*^BuOOH, ^•^OH, ^•^NO, ROOR’, ROO^•^ and ONOO^−^) showed no fluorescence enhancement and no bathochromic shift. These results indicate that our design in **1b** achieved a high selectivity towards HOCl and displayed a unique emission red-shift of 30 nm from original 525 nm to 555 nm (Fig. 1h, Supplementary Figure 48).

### Validation of new HOCl detection mechanism

NMR titration was performed to further validate the HOCl detecting mechanism of **1b**. The NMR titration of **1b** showed a gradual downfield shift (labeled as proton *e* in Fig. 2a) of *β*-hydrogen at the 2,6-positions of the pyrrole moiety from ∼6.7 ppm to ∼7.0 ppm after the addition of HOCl. The peak integration decreased from two to one (Fig. 2a), as the structure transitioned from the non-chlorinated reactant (labeled as proton *e*) to mono-chlorinated product (labeled as proton *e’*), suggesting the substitution of the *β*-hydrogen on pyrrole by a electron-withdrawing Cl atom. As **1b** and its reaction product with HOCl contain a +1 charge, HRMS served as an effective way to verify the detection mechanism. The reaction mixture showed the peaks [M-H+Cl]^+^ with *m/z* = 374.1403 and [M-2H+2Cl]^+^ with *m/z* = 408.1015, corresponding to mono-chlorinated product and di-chlorinated product respectively (Fig. 2b-c). Similar chlorination products were also observed in the other two methylated BODIPYs **2b** and **3b** (Supplementary Figure 49) as well as the non-methylated BODIPYs **1a, 2a** and **3a** (Supplementary Figure 41). In addition, the *m/z* peaks corresponding to the **1b** remained unchanged in the presence of both HOCl and the scavengers (Supplementary Figure 50). The results of the HRMS and NMR data are consistent with the proposed HOCl detecting mechanism that HOCl chlorinates the BODIPY core of **1a**-**b, 2a**-**b** and **3a**-**b**, leading to a red-shift in its fluorescence emission spectrum and a change in its quantum yield.

**Figure 2.**
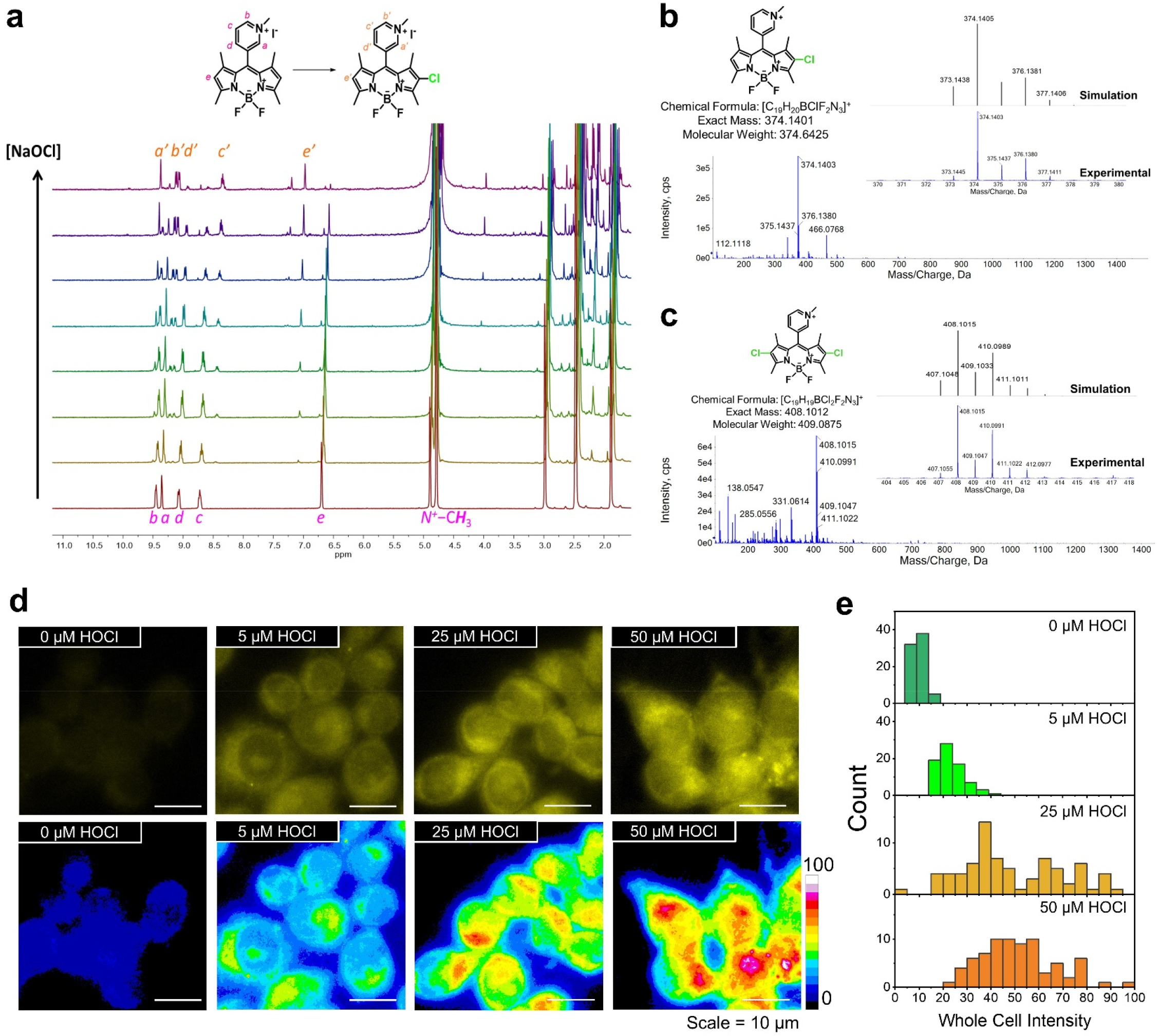
HOClSense detects HOCl based on fluorophore chlorination. **a, 1b** undergoes chlorination with HOCl. Stacked ^1^H NMR spectrum for **1b** with the titration of increasing amounts of NaOCl. **b**−**c**, Mass spectral characterization of HOCl chlorinated products from **1b**. Diagram of HRMS (+ve ESI) spectrum of **1b** after the addition of HOCl, showing the **b**, chlorination and **c**, dichlorination reaction product. **d**, In-cell HOCl calibration. Representative fluorescence images of **1b**-labelled RAW 264.7 cells clamped at the indicated [HOCl]. **e**, Histogram for the whole cell intensity of **1b**-labeled cells at the indicated [HOCl]. Imaging experiments were performed in triplicate.

To further understand the design principles of this new class of HOCl-sensing fluorophores, several structures with variations at the 2,6-positions were synthesized to mask the *β*-hydrogens on the pyrroles. The HOCl detecting ability of these variations (**1c**−**f**) were investigated to determine the structure-activity relationship. The *β*-hydrogen were replaced by ethyl substituents, resulting in the formation of di-substituted **1d** and mono-substituted **1f**. When the *β*-hydrogen on pyrroles were absent in **1d**, addition of HOCl resulted in disappearance of the characteristic BODIPY absorption peak. (Supplementary Figure 40). HRMS analysis of the reaction mixture (**1d** with HOCl) suggested structural decomposition including demethylation and non-selective chlorination (Supplementary Figure 51). In such way, the d-PeT is not able to switch off and results in fluorescence quenching. For mono-substituted **1f**, the reaction mixture showed a peak corresponding to the chlorinated product [M-H+Cl]^+^ with *m/z* = 402.1719 (Supplementary Figure 52). These results demonstrate that the *β*-hydrogens at the 2,6-positions of the pyrrole core can direct a selective chlorination to produce the characteristic “Turn-On, red-shift” fluorescence response. In other words, the absence of substituents at the 2- or 6-positions of the BODIPY core in **1b** is essential for HOCl-induced chlorination, which subsequently alters the emission wavelength and fluorescence intensity of **1b**.

### *in cellulo* HOCl calibration of HOClSense dyes

HOClSense dyes were assessed for their effect on cell viability. **1a, 1b**, and **3b** exhibited almost no cytotoxicity up to 100 µM (Supplementary Figure 53). These dyes were selected for cell imaging applications based on their favorable fluorescence response, high specificity towards HOCl, and minimal cytotoxicity. RAW 264.7 cells were incubated with the dyes for 30 min (referred to as the ‘pulse’), washed the cells, and then allowed them to grow in complete medium for 1 h (referred to as the ‘chase’). The fluorescence intensity of **1b**-labeled cells was significantly higher than that of the unlabeled cells. The uptake experiment results demonstrated that **1b** effectively labeled RAW 264.7 cells (Supplementary Figure 54). The intracellular HOCl-sensing abilities of **1a, 1b**, and **3b** in HOCl-clamped RAW 264.7 cells were then investigated (Supplementary Figure 55). Upon cell fixation, the dyes distributed throughout the cytosol. The intracellular HOCl calibration result is consistent with the *in vitro* experiments, showing a dose-dependent response to [HOCl] for **1a, 1b**, and **3b** (Fig. 2d-e, Supplementary Figure 56-58). The fluorescence intensity of **1b**- and **3b**-labeled cells increased with increasing concentration of clamped HOCl. Conversely, **1a** detected HOCl with a “Turn-Off” response, as the intracellular fluorescence intensity decreased with increasing [HOCl]. These results indicate that **1a, 1b**, and **3b**, which detect HOCl through a novel chemical mechanism, are capable of monitoring intracellular HOCl production at physiological concentrations (20 to 400 μM).^56,57^

### Functionalization of HOClSense dyes for extracellular HOCl detection

The pyridinium design of HOClSense dyes also supported chemical functionalization, as the pyridine of HOClSense dyes was PEGylated with azide-PEG_3_-iodide through an S_*N*_2 reaction to form **1b-azide** (Fig. 3d). The resulting **1b-azide** was conjugated with different organelle-targeting groups or carriers through strain-promoted azide-alkyne cycloaddition. This modification did not extensively alter the pyridinium BODIPY structure, and the conjugation by click reaction did not significantly affect the fluorescence characteristics of HOClSense dyes. For proof of concept, we targeted the plasma membrane with HOClSense to evaluate extracellular HOCl exposure as HOCl serves as a biomarker for chronic inflammatory conditions,^58^ and elevated extracellular levels of HOCl have been associated with necrotic cell death.^57^ A reported plasma-membrane-labeling ligand **mb**^59,60^ was utilized to prepare **DBCO-mb**, a ligand for efficient labeling via click reaction. **mb** was confirmed to target the plasma membrane, guiding the BODIPY dye to specifically label the membrane instead of the entire cell (Fig. 3b-c, Supplementary Figure 59a-d). By clamping the cells with HOCl, it was shown that **1b-mb** effectively detects HOCl around the membrane, consistent with in-cell HOCl calibration of **1b** (Fig. 3f, Supplementary Figure 60-61). PAMPs, including lipopolysaccharides (LPS) and phorbol 12-myristate 13-acetate (PMA), are known to induce HOCl production. By incubating LPS and PMA with RAW 264.7 macrophages, we successfully employed **1b-mb** to monitor the resultant HOCl levels, demonstrating the probe’s efficacy in detecting extracellular HOCl. A significantly higher pseudo-color intensity was observed across the cell membrane when the cells were stimulated with PAMPs (Fig. 3e-g, Supplementary Figure 62). These results demonstrated that **1b-mb** could monitor the extracellular HOCl exposure of the plasma membrane.

**Figure 3.**
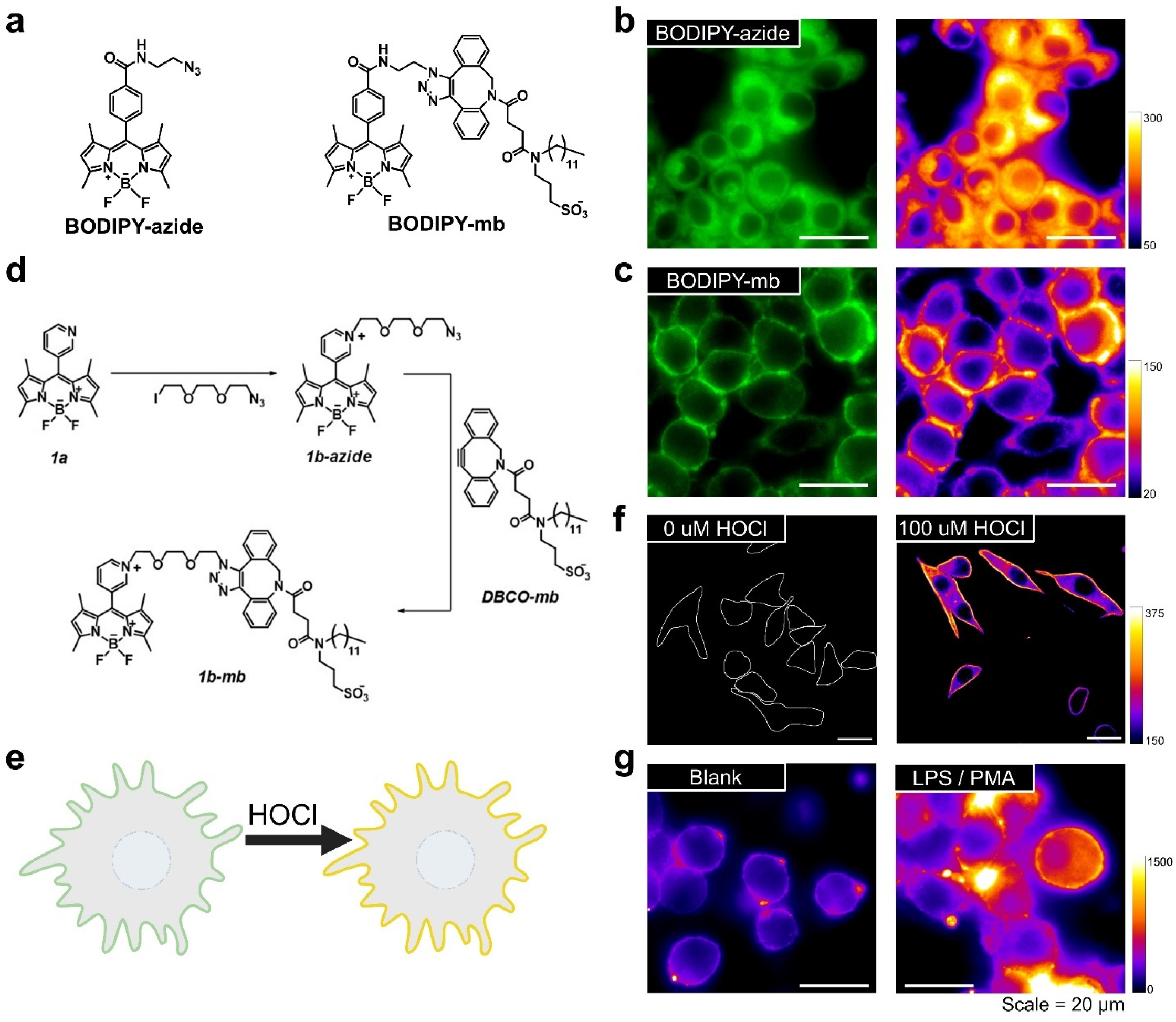
Functionalization of HOClSense dye for plasma membrane labeling and extracellular HOCl mapping. **a**, Structures for **BODIPY-azide** and **BODIPY-mb.** **b**-**c**, Representative fluorescence images of RAW 264.7 cells labeled with **BODIPY-azide** or **BODIPY-mb** respectively. **d-f**, Functionalization of **1b** for plasma membrane labeling. **d**, Synthesis scheme of **1b-mb.** **f**, In-cell HOCl calibration. Representative fluorescence images of **1b-mb**-labelled cells clamped at the indicated [HOCl]. **e**, Schematic diagram illustrating **1b-mb** response to HOCl. **g, 1b-mb** detects PAMP-stimulated HOCl production. Representative fluorescence image of **1b-mb**-labelled RAW 264.7 cells upon stimulation of LPS and PMA. Experiments were performed in triplicate.

### Functionalization of HOClSense dyes for lysosomal HOCl detection

MPO is localized in lysosomes, which are central to various metabolic pathways. Elevated HOCl levels in lysosomes are associated with numerous diseases and may reflect cellular integrity and functionality. Therefore, measuring lysosomal HOCl levels, without triggering an immune response, can serve as a valuable indicator of lysosomal integrity and overall cellular health. To detect lysosomal HOCl production without stimulating the immune response, **1b** was conjugated to 10k molecular weight dextran (Fig. 4a), which can be taken up by cells through fluid-phase endocytosis and reach the endolysosomes (Fig. 4b).^61-63^ **Dextran-1b** was prepared via a two-step functionalization and click chemistry (Fig. 4a). The DBCO linker was conjugated to amino-dextran to form **DBCO-dextran**, and then conjugated it with **1b-azide** to form **dextran-1b**. Using **dextran-1b**, we measured lysosomal HOCl production in RAW 264.7 cells following PAMP stimulation and observed an increase in lysosomal fluorescence intensity. To confirm that this increase was due to MPO-produced HOCl, cells were treated with 50 µM MPO inhibitor aminobenzoic acid hydrazide (4-ABAH). The decreased lysosomal fluorescence intensity upon inhibitor treatment confirmed that the fluorescence increase was due to MPO-dependent HOCl production (Fig. 4c-d). After validating the lysosomal HOCl-monitoring ability of **dextran-1b**, we aimed to determine if the STING pathway induces HOCl production, given that STING pathway is a key component of the innate immune system that recognizes bacterial secondary messengers and has recently been implicated in various neurodegenerative diseases.^64^ Upon STING activation with DMXAA, a significant increase in lysosomal fluorescence intensity was observed (Fig. 4e-f). Cells pre-treated with H-151, a potent STING antagonist that inhibits STING palmitoylation, showed a reduction in **dextran-1b** signal. This reduction indicates that the increased signal is directly attributable to STING activation. The observed HOCl generation upon STING activation highlights a compelling correlation between the STING signaling axis and the resulting HOCl production.

**Figure 4.**
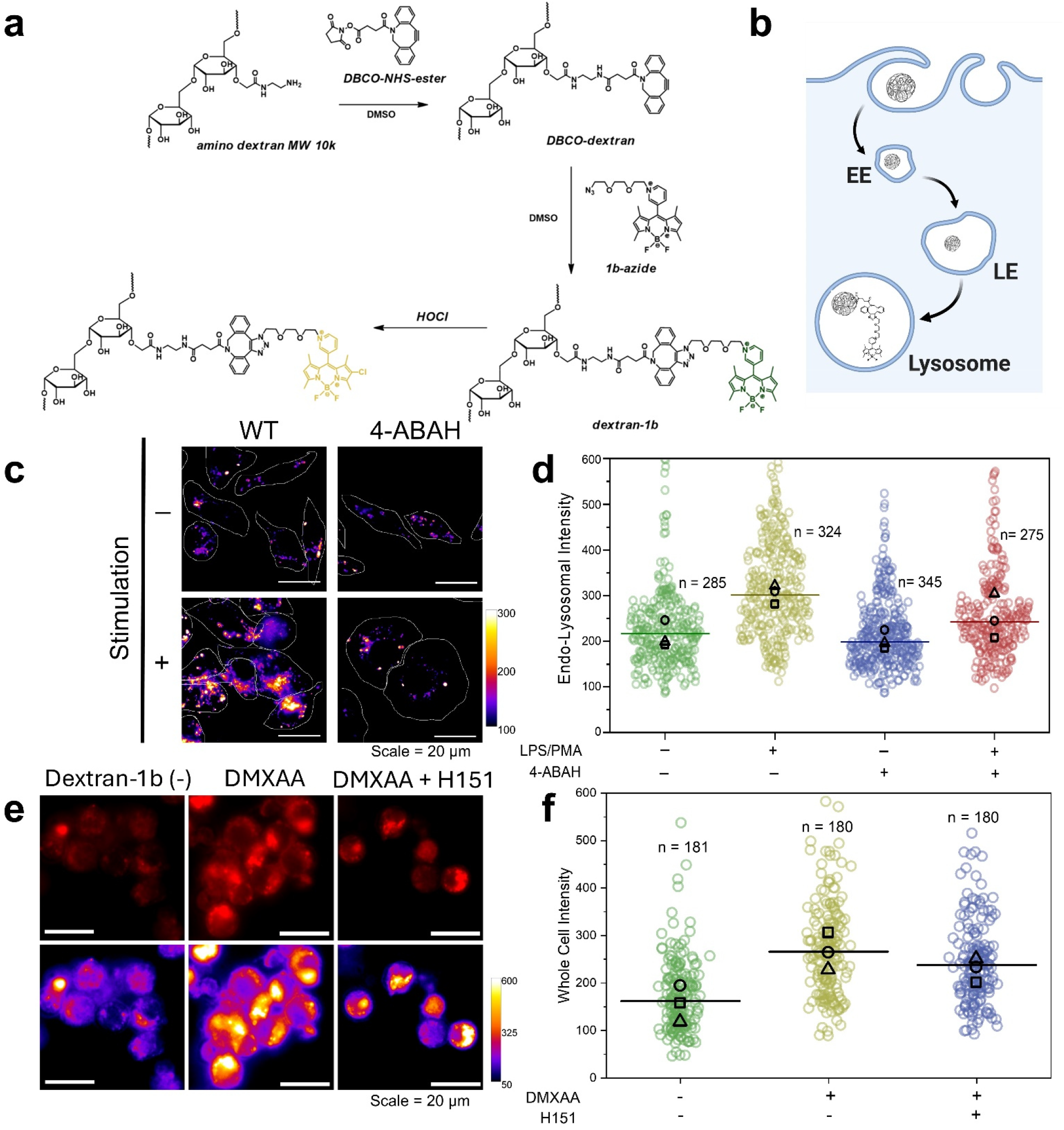
Functionalization of HOClSense dyes for lysosomal HOCl imaging. **a**, Synthesis scheme of **dextran-1b**. By reacting the **1b-azide** with DBCO-functionalized amino dextran, **dextran-1b** is synthesized, combining the fluorescent properties of **1b** with the lysosome-labeling capabilities of dextran. **b**, Trafficking of **dextran-1b** through endocytosis pathway (EE, early endosome; LE, late endosome; LY, lysosome). **c-d, dextran-1b** detects PAMP-stimulated lysosomal HOCl production. **c**, Representative fluorescence images of **dextran-1b**-labelled RAW 264.7 cells showing the enhanced fluorescence intensity upon PMAPs stimulation while the enhanced signal was reduced upon treatment of MPO inhibitor 4-ABAH. **d**, Quantification of endolysosomal fluorescence intensity of **dextran-1b**-labelled RAW 264.7 cells at the indicated conditions. Experiments were performed in triplicate. The median value of each trial is given by a square, circle, and triangle symbol (n = number of lysosomes). **e-f**, STING activation induces HOCl production. **e**, Representative fluorescence images of **dextran-1b**-labelled RAW 264.7 with or without murine STING stimulator dimethylxanthenone acetic acid (DMXAA, 30 μM) and STING inhibitor H151 (15 μM). **f**, Quantification of whole cell fluorescence intensity of **dextran-1b**-labelled RAW 264.7 cells at the indicated conditions. Experiments were performed in triplicate. The median value of each trial is given by a square, circle and triangle symbol, and median of whole experiment represented by line (n = number of cells).

### Basal HOCl production in Niemann–Pick diseases

Niemann–Pick (NP) disease is a lysosomal storage disorder sub-classified into three types: A, B, and C. Types A and B are caused by mutations of acid sphingomyelinase (ASM), while type C results from mutations of NPC1 or NPC2 protein.^62,65^ These mutations lead to abnormal accumulation of lysosomal cargos. Elevated ROS levels is also observed in NP phenotype cells due to altered mitochondrial and peroxisomal function.^66^ Therefore, **dextran-1b** was utilized to assess the lysosomal HOCl level in Niemann–Pick diseases. To prepare the pharmacologically induced NP-A/B and NP-C cell models, cells were pre-incubated with amitriptyline hydrochloride (AH) and U18666A, which are inhibitors of ASM and NPC1, respectively, to mimic the deficiency of these lysosomal proteins. Without any PAMPs stimulation, the lysosomal HOCl levels in both NP-A/B and NP-C models were higher than those in WT cells (Fig. 5a-b). The NP-A/B model exhibited a two-fold increase in lysosomal fluorescence intensity, while the NP-C model displayed a four-fold increase in lysosomal fluorescence intensity. Upon PAMPs stimulation, enhanced lysosomal fluorescence intensity was observed in both NP-A/B and NP-C models, with a marked hyperproduction of HOCl in the NP-C model (Fig. 5a-b). Based on the results from the NP cell models, whether patient samples exhibit elevated basal levels of HOCl production was investigated. **Dextran-1b** was utilized to evaluate the HOCl level of primary cells derived from normal healthy individuals and NP-C patients. Our results showed that NP-C patient fibroblasts had a higher average HOCl level than normal healthy individuals (Fig. 5c-d) and these results were consistent with the observed elevated basal HOCl production in the pharmacologically induced NP-A/B and NP-C cell models. Given that inflammation is a significant driver of NPC pathology, this previously unreported basal HOCl production provides novel insights into the mechanisms underlying the disease.

**Figure 5.**
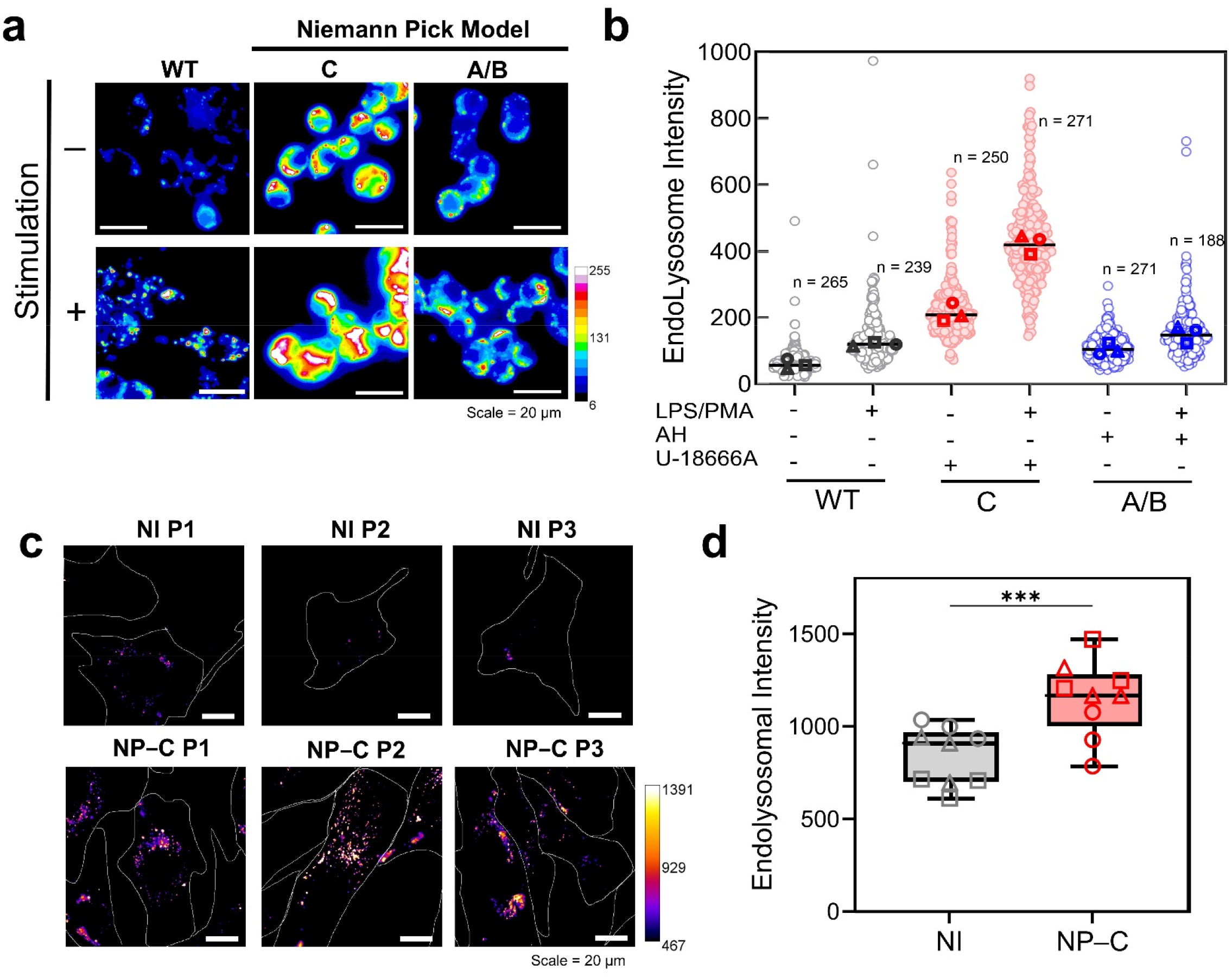
HOClSense dye detects the basal lysosomal HOCl production in Niemann–Pick diseases. **a**-**b**, HOClSense dyes detects basal lysosomal HOCl production in pharmacologically induced NP A/B and C cell models. **a**, Representative fluorescence images of **dextran-1b**-labelled RAW 264.7 cells treated with 65 µM amitriptyline (AH) or 20 µM U18666A with and without stimulation of LPS and PMA. **b**, Quantification of lysosomal fluorescence intensity in indicated condition. **c**-**d** HOClSense dye detects the basal lysosomal HOCl production in primary skin fibroblast divided from NP-C patients. **c**, Representative fluorescence images of **dextran-1b**-labeled primary skin fibroblast from apparently healthy individuals and NP-C patients. **d**, Quantification of lysosomal fluorescence intensity of fibroblast samples from apparently healthy individuals and NP-C patients. Experiments were performed in triplicate for each cell line sample. The median value of each trial is given by square, circle, and triangle symbols. Error bars indicate the mean ± standard error of the mean (s.e.m.) of nine independent measurements. \*\*\**P* < 0.001. One-way analysis of variance (ANOVA) followed by Dunnett’s test for multiple comparison.

## Discussion

Through a strategic methylation/alkylation design, we effectively overcame four critical challenges simultaneously, providing robust and precise chemical tools for resolving subcellular HOCl. The water solubility of pyridyl BODIPYs was greatly enhanced through the methylation/alkylation because of bearing +1 charge. After the methylation/alkylation, the fluorescence of pyridinium BODIPYs (**1b, 2b** and **3b**) switched to a “turn-on” response towards HOCl based on the d-PeT mechanism while most fluorophores including pyridyl BODIPY (**1a, 2a** and **3a**) feature a “turn-off” response. In addition, the extended glycol linker in **1b-azide** facilitated easy functionalization, enabling the installation of organelle-targeting properties. As a result, membrane-targeting **1b-mb** and lysosome-targeting **dextran-1b** were successfully prepared and demonstrated for subcellular HOCl imaging. Finally, changing the *N*-methylated-3-pyridyl group to *N*-methylated-4-pyridyl group at the 8-position of the BODIPY shifted the emission of the HOCl-sensing fluorophores from yellow to orange. Furthermore, it is noteworthy that fluorescence ‘turn-on’ response with a peak shift is particularly useful for imaging, as it enables the monitoring of fluorescence intensity at the shifted emission peak, resulting in minimal background signal in the absence of the target analyte and a significantly higher fold change upon its presence (Supplementary Figure 45).

The new chlorination-based HOCl detection mechanism allows us to have a universal method to functionalize the developed HOClSense dyes for subcellular imaging. For proof of concept, we targeted plasma membrane and lysosomes with HOClSense for monitoring extracellular HOCl exposure and lysosomal HOCl production. The capability of HOClSense for subcellular HOCl mapping was also demonstrated by monitoring PAMPs-induced lysosomal HOCl. The lysosomal HOCl production upon STING pathway activation was then visualized, marking the first report of STING signaling-induced HOCl production. It demonstrated the capability of HOClSense dyes to monitor HOCl distribution, enabling the study of its impact on various cellular compartments.

HOCl overproduction was also observed in Niemann–Pick disease samples using our lysosome-labeling HOCl indicator. Our findings indicate that hyper-HOCl-production is present in pharmacologically induced NP-A/B and NP-C cell models upon PAMPs stimulation. Additionally, an increased basal HOCl production was detected even in the absence of PAMPs stimulation, a phenomenon also observed in primary cells from NPC patients. To the best of our knowledge, this is the first report of abnormal higher HOCl basal level in NP diseases. These findings suggest that elevated basal HOCl production may contribute to the pathology of Niemann–Pick disease. Given that inflammation is a significant driver of NPC pathology,^67,68^ this unreported basal HOCl production provides novel insights into the mechanisms underlying the disease.

In conclusion, a series of HOClSense dyes were developed and featured a novel chlorination-based detection mechanism. It is promising that the minimalistic design of HOClSense dyes also allows for tuning the emission, adjusting detection mode and functionalizing at the same time, which greatly shortens the time for synthesis and further modification. All these properties make these dyes ideal for multispectral imaging. Structural activity relationship of HOClSense dyes was studied and we discovered that *β*-hydrogen on the pyrrole of BODIPY is essential for the directed chlorination by HOCl. The strategic alkylation design is beneficial to modulate the cellular distribution of HOClSense by click conjugation. As a proof of concept, we targeted plasma membrane and lysosomes with HOClSense for monitoring extracellular HOCl exposure and lysosomal HOCl production. Our tools successfully monitor the PAMPs-induced extracellular and lysosomal HOCl, find out the HOCl production induced by STING activation, and discovered the elevated basal HOCl production in drug-induced NP disease cell models and primary fibroblast samples from NP patients.

## Supporting information

Supporting information_synthesis

## Declaration of interests

The authors declare no competing financial interests.

## Resources availability

We are delighted to provide our developed tools upon request.

## Acknowledgements

This work was supported by NIH grants R35GM147112 (K.L.), R35GM147112-02S2 (K.L.), R35GM147112-02S1 (K.L. & M.S.), and Clarkson University start-up fund. We thank Patrick Lutz and Samuel Tartakoff from St. Lawrence University for providing NMR support. We extend our sincere gratitude to St. Lawrence University for generously providing access to their JEOL 400 MHz nuclear magnetic resonance (NMR) spectrometer.

## Author contributions

F.T., L.T., and K.L. wrote the manuscript. F.T., L.T., wrote the supporting information with the input of all authors. All authors discussed the results and commented on the manuscript. F.T., L.T., and K.L. designed the project. Compounds **1a**-**1f, 2a**-**b, 3a**-**b, 1b-N**_**3**_ were synthesized by F.T., L.T., N.P., and M.S. **DBCO-NHS, DBCO-mb** were synthesized by M.R.N. and M.A. The *in vitro* characterization, NMR titration, mass spectrometry, cellular distribution, *in cellulo* HOCl clamping, HOCl imaging were performed by F.T. and L.T. HOCl production during STING activation was evaluated by C.H. J.M performed the cell viability assay and provide imaging supporting.

## Notes

### Competing Interest Statement

The authors have declared no competing interest.

