## Supplementary material for "Leveraging Chlorination-Based Mechanism for Resolving Subcellular Hypochlorous Acid"

### METHODS
### Chemicals
Chemicals were purchased from TCI, Sigma Aldrich, Ambeed, Thermo Fisher Scientific, Alfa Aesar, Supelco., Lumiprobe, Fina Biosolutions, and Cayman Chemicals (see **Supplementary Table 1**).

**Supplementary Table 1:** Information on Purchasing Source.

| Supplier | TCI | Sigma Aldrich | Ambeed | Thermo Fisher Scientific | |
|---|---|---|---|---|---|
| Chemicals | β-Nicotinamide adenine dinucleotide, disodium salt, hydrate (NADPH) | Sodium chloride | Hexafluorophosphate Azabenzotriazole Tetramethyl Uronium (HATU) | Aluminum oxide (neutral, Brokmann I, 4-300 µM, 60 Å) | Potassium superoxide ($KO_2$) |
| | *tert*-Butyl hydroperoxide (70% w/v) | Potassium chloride | 2,4-dimethylpyrrole | *N,N*-Diisopropylethylamine (DIPEA), | Di-*tert*-butyl peroxide |
| | Glutathione (GSH) | Calcium chloride dihydrate | 4-pyridinecarboxaldehyde | Glutathione Disulfide (GSSG) | Iodomethane (MeI) |
| | Boron trifluoride diethyl etherate ($BF_3$•$O(C_2H_5)_2$) | Magnesium chloride hexahydrate | 3-pyridinecarboxaldehyde | Ammonium Hexafluorophosphate ($NH_4PF_6$) | Manganese nitrate tetrahydrate |
| | 2-pyridinecarboxaldehyde | Phorbol 12-myristate 13-acetate (PMA) | 3-Ethyl-2,4-dimethylpyrrole | Sodium Hypochlorite (NaOCl, 13% w/v), | Iron(II) sulfate heptahydrate |
| | 3-Ethyl-2,4-dimethylpyrrole | Lipopolysaccharides (LPS) from Escherichia coli (O111:B4) | | Hydrogen peroxide ($H_2O_2$, 30% w/v) | Iron(III) chloride hexahydrate |
| | | | | 2,2'-Azobis(2-methylpropionamidine) dihydrochloride (AAPH) | Cobalt(II) nitrate hexahydrate |
| | | | | Sodium hypochlorite, 11-15% | Nickel(II) chloride hexahydrate |
| | | | | Hydrogen peroxide 30% | Copper(II) chloride dihydrate |
| | | | | | Zinc nitrate hexahydrate |
| Supplier | Alfa Aesar | Supelco. | Lumiprobe | Fina Biosolutions | Cayman Chemicals |
| Chemicals | *N*-Acetylcysteine (NAC) | TLC silica gel (60 Å, $F_{254}$) | Azide-PEG3-iodide (CAS 1309457-01-9) | Amino Dextra 10 kDa (AD10x10) | U-18666A |
| | | Silica gel (60 Å, 70-230 mesh) | | | Spermine NONOate |
| | | | | | Amitriptyline hydrochloride (AH) |

### UV-Vis absorption, fluorescence spectroscopy, and selectivity assay
Solution samples were contained in quartz cuvettes with a volume of 1.5 mL, 1 cm of path length and 0.4 cm of slit length. All aqueous solutions were prepared with Milli-Q water (18.2 MΩ•cm). Briefly, stock DMF solution of HOClSense dyes (1 mM) and stock analyte solutions (1 mM / 2 mM) were freshly prepared and added to aqueous solution according to the detection condition to form a final mixture with a volume of 1 mL. The final concentration of HOClSense dyes is 5 µM, and the final concentration of ROS/ RNS analyte is 50 µM (10 mol. equiv.). All ROS and RNS were freshly prepared or diluted as described below:
HOCl: Hypochlorite was diluted to 1 mM from a 13% *w/w* stock NaOCl solution.
$H_2O_2$: Hydrogen peroxide was diluted to 1 mM from a 30% *w/w* $H_2O_2$ stock solution.

ROO$^{\bullet}$: Peroxy radical was generated by a 1 mM AAPH solution (2,2'-Azobis(2-methylpropionamidine) dihydrochloride, CAS 2997-92-4).

$O_2^{\bullet-}$: Superoxide was prepared from saturated $KO_2$ solution in DMSO (~1 mM).[1]

$^{\bullet}$NO: Nitric oxide was generated from a 1 mM spermine NONOate solution.

$^{\bullet}$OH: Hydroxyl radical was generated by Fenton reaction ($Fe^{2+} + H_2O_2 \rightarrow Fe^{3+} + {}^{\bullet}OH + OH^-$). The stock $H_2O_2$ solution was added to the sensor solution containing ferrous ion (1 mM, $FeSO_4 \bullet 7H_2O$) The final concentration of $^{\bullet}$OH was estimated to be the same as the final $H_2O_2$ concentration (50 µM).

ROOR': Alkyl peroxide was diluted to 1 mM solution from di-*tert* butyl peroxide.

$ONOO^-$: The interaction between $H_2O_2$ and $NaNO_2$ was utilized to produce peroxynitrite ($ONOO^-$) according to the literature. The concentration of $ONOO^-$ was determined from absorbance at 302 nm (ε = 1670 L mol$^{-1}$ cm$^{-1}$) and diluted to 1 mM solution for use.[2]

$^{t}$BuOOH: *Tert*-butyl hydroperoxide was diluted to 1 mM from a 70% *w/w* stock $^{t}$BuOOH solution.

The final reaction mixture was vortexed and incubated at ambient condition for 15 min before emission measurement. Fluorescence spectra and UV–Vis absorption spectra were collected on a PTI QM–4/2005 spectrometer and an Agilent Cary 8454 UV–Vis Diode Array System, respectively.

**Effect of pH**

Aqueous solutions and buffer solutions were prepared with ranges of pH. The pH of aqueous solutions was adjusted using NaOH (1 M) or HCl (1 M). Citric phosphate buffer (10 mM) was prepared and adjusted with varying concentrations of citric acid and $Na_2HPO_4$. Stock DMF solution of HOClSense dye (1 mM) and stock analyte solution (2 mM) were freshly prepared. HOClSense dye and analyte were added to aqueous solutions or buffer solutions with a final HOClSense dye concentration of 5 µM, a final analyte concentration of 50 µM, and a final volume of 1 mL. Fluorescence spectra were collected for the resulting solutions.

**Ion interference**

Stock solutions of ions were prepared using sodium chloride (NaCl), potassium chloride (KCl), calcium chloride dihydrate ($CaCl_2 \bullet 2H_2O$), magnesium chloride tetrahydrate ($MgCl_2 \bullet 6H_2O$), manganese nitrate tetrahydrate ($Mn(NO_3)_2 \bullet 4H_2O$), iron(II) sulfate heptahydrate ($FeSO_4 \bullet 7H_2O$), Iron(III) chloride hexahydrate ($FeCl_3 \bullet 6H_2O$), Cobalt(II) nitrate hexahydrate ($Co(NO_3)_2 \bullet 6H_2O$), nickel(II) chloride hexahydrate ($NiCl_2 \bullet 6H_2O$), copper(II) chloride dihydrate ($CuCl_2 \bullet 2H_2O$), and zinc nitrate hexahydrate ($Zn(NO_3)_2 \bullet 6H_2O$). HOClSense dye, analyte solution, metal ion solution, and Milli-Q water were added together to make a final HOClSense dye concentration of 5 µM, a final analyte concentration of 50 µM, a final metal ion concentration of either 20 µM or 100 mM, and a final volume of 1 mL. Solutions were prepared in the presence and absence of HOCl analyte. Fluorescence spectra were collected for the resulting solutions.

**Effect of ROS scavenger**

Stock solutions of ROS scavengers were prepared using *N*-Acetyl-L-Cysteine (NAC), reduced Glutathione (GSH), oxidized Glutathione (GSSG), and reduced β-Nicotinamide adenine dinucleotide, disodium salt, hydrate (NADPH). Stock DMF solution of HOClSense dye (1 mM) and stock analyte solution (2 mM) were freshly prepared. HOClSense dye, ROS scavengers, and analyte were added to aqueous solution with a final HOClSense dye concentration of 5 µM, a final analyte concentration of 50 µM, a final concentration of 300 µM *N*-Acetyl-L-Cysteine, 1 mM GSH, 1 mM GSSG and 100 µM NADPH, and a final volume of 1 mL. Fluorescence spectra were collected for the resulting solutions.

**Mammalian cell culture**

RAW 264.7 cells (ATCC Number: TIB-71™) and Primary Human Dermal Fibroblasts (HDF, ATCC Number: PCS-201-012™) were purchased from American Type Culture Collection (ATCC). DMEM/F12 was purchased from Thermo Fisher Scientific. DMEM (10% FBS) was purchased from Corning. RAW 264.7 cells were cultured in Dulbecco's Modified Eagle's Medium (DMEM) and 10% fetal bovine serum (FBS) with Pen-Strep (100 U mL$^{-1}$ - 100 µg mL$^{-1}$). HDF cells were cultured in Dulbecco's Modified Eagle's Medium/ Gibco Ham's Nutrient Mix F-12 (DMEM/F12) and 10% fetal bovine serum (FBS), Pen-Strep (100 U mL$^{-1}$ - 100 µg mL$^{-1}$). Cells were cultured in 37 °C with 5% $CO_2$ atmosphere.

**Wide field Fluorescent Imaging**

An Olympus IX83 Inverted Microscope with a Photometrics Prime BSI CMOS Camera was used for all fluorescent imaging experiments. Olympus's cellSens Dimension 4.1 program was used to control the CMOS camera, shutter, and filter cubes. Images obtained were processed using ImageJ (NIH). Background intensity was subtracted for all images by taking mean intensity over an adjacent cell-free area. For details on which of the fluorescence imaging channels (and their respective excitation filter, emission filter, and dichroic filter) were used on which compounds, see **Supplementary Table 2**.

**Supplementary Table 2:** Information on imaging channels, respective filters, and which compounds were used.

| Channel | Excitation filter | Emission filter | Dichroic filter | Compounds |
|---|---|---|---|---|
| GFP | AT480/30x band pass | AT535/40M band pass | AT505DC | 1a, 1b-mb |
| Cy3 | ET545/25x band pass | ET605/70M band pass | T565lpxr | 1a, 1b, 3b, dextran-1b, 1b-mb |

**HOCl clamping**

RAW 264.7 cells were plated on 35 mm Glass Bottomed Dishes (catalog #: D35-10-1.5-N, CellVis, Mountain view, CA, USA) and incubated overnight with complete medium. This was followed by pulsing with 20 μM of **1a**, **1b**, or **3b** in OPTI-MEM for 1 h at 37 ℃ (100 nM of **1b-mb** in OPTI-MEM for 10 min at 37 ℃). Next, the cells were washed with 1×PBS thoroughly and chased with complete medium for 1 h at 37 ℃ (0 min for **1b-mb**). The media was removed, and the cells were fixed with 100 % MeOH at 4℃ for 30 min. The fixing solution was removed, and the cells were incubated in the clamping buffer solutions of indicated HOCl concentration for 24 h at 4 ℃. Cells were imaged in the HOCl clamping buffer using a 100×, 1.45 NA, apochromat oil immersion objective (UPlanXApo, Olympus Corporation of the Americas, Center Valley, PA, USA). **1a** was imaged under the GFP channel, **1b-mb** was imaged under GFP and Cy3 channel, and all others were imaged under the Cy3 channel. HOCl clamping buffer solutions were freshly prepared in UB4 (pH 6.0, 20 mM HEPES, 20 mM MES, 20 mM NaOAc, 140 mM NaCl, 10 mM KCl and 2.5 mM $CaCl_2$) at varying [HOCl] using a stock 1.9 M NaOCl solution. Whole cell intensity was measured for each cell by taking mean intensity of defined cell specific ROIs.

**Cell viability assay**

In a 96-well culture plate, $0.4\times10^5$ cells were seeded overnight. Cells were treated with HOClSense dye diluted to specified working concentrations in OPTI-MEM for 1 h followed by washing with PBS and then incubated in cell media for 0.5 h. The CellTiter-Blue® Cell Viability Assay was then performed according to the manufacturers' protocol (Promega, Madison, Wi, USA). Plates were read using SpectraMaxTM i3/i3x multi-mode plate reader.

**Extracellular ROS detection**

RAW 264.7 cells were plated on 35 mm Glass Bottomed Dishes (catalog #: D35-10-1.5-N, CellVis, Mountain view, CA, USA) and incubated overnight with complete media. Cells were treated with 80 ng $mL^{-1}$ lipopolysaccharides (LPS) and incubated for 3.5 h at 37 ℃. Without taking off the media, they were treated with 10 ng $mL^{-1}$ Phorbol 12-myristate 13-acetate (PMA) and incubated for 30 min at 37 ℃. Next, the cells were washed with 1×PBS. This was followed by pulsing with **1b-mb** diluted to 20 μM in OPTI-MEM (2% DMSO) for 10 min. Next, the cells were washed with 1×PBS three times. The cells were buffered using UB4 (pH 6.0, 20 mM HEPES, 20 mM MES, 20 mM NaOAc, 140 mM NaCl, 10 mM KCl and 2.5 mM $CaCl_2$) and were observed using a 100×, 1.45 NA, apochromat oil immersion objective (UPlanXApo, Olympus Corporation of the Americas, Center Valley, PA, USA). Images were obtained in the GFP channel.

**Pharmacologically induced cell culture model for Niemann–Pick Disease type A/B and type C**

Niemann–Pick type A/B and type C models were prepared according to the reported literature.[3,4] RAW 264.7 cells were incubated with 65 µM ASM inhibitor Amitriptyline Hydrochloride (AH) or 20 µM NPC1 protein inhibitor U-18666A overnight (for 24 h) to mimic NP type A/B or NP type C cell conditions respectively. Then cells were activated for ROS production using LPS (500 ng $mL^{-1}$ for 4 h) and PMA (500 ng $mL^{-1}$ for 0.5 h) in wildtype, type A/B and type C conditions. Then the cells were pulsed with **dextran-1b** (0.1 mg $mL^{-1}$) for 30 min, washed with 1×PBS twice and then chased with either AH (65 µM) or U-18666A (20 µM) for 1 h. The cells were washed with 1×PBS and imaged in OPTI-MEM containing either AH (65 µM) or U-18666A (20 µM). Endo-lysosomal intensity was measured for each data point by taking mean intensity of defined cell specific ROIs. Images were obtained in Cy3 channel and the intensities for the endo-lysosomal intensities were analysed.

**Lysosomal HOCl measurements using human fibroblast from healthy normal individuals and NP disease type C patients.**

Human dermal fibroblasts were maintained in DMEM/F12 medium with 10% heat inactivated FBS while NP disease type C patients were maintained in MEM with 15% FBS (see **Supplementary Table 3** for detailed information of cell lined used). The cells were pulsed with **dextran-1b** (0.5 mg mL-1) for 2 h, washed with 1×PBS twice and chased in DMEM media for 15 h. The media was changed to OPTI-MEM and imaged. Endo-lysosomal intensity was measured for each data point by taking mean intensity of defined cell specific ROIs. Images were obtained in Cy3 channel and the intensities for the endo-lysosomal intensities were analysed.

**Supplementary Table 3:** Information of human primary skin fibroblast used in this study.

| Sample | Cell Line Name | Gene | Gene Mutation | Age (at sampling) |
|---|---|---|---|---|
| **NPC-P1** | GM00110 | NPC1 | 2215(or 2217)delTCCTTT(or CTTTTC) | M (9 YR) |
| **NPC-P2** | GM17913 | NPC1 | VAL1165MET | M (23 YR) |
| **NPC-P3** | GM18388 | NPC1 | ARG404GLN | F (No Data) |
| **NI-1** | HDFa (ATCC) | N/A | N/A | N/A |
| **NI-2** | GM21808 | N/A | N/A | Male (1 DA) |
| **NI-3** | GM22136 | N/A | N/A | Male (1 DA) |

where DA = day, YR = year.

**Supplementary Table 4. Summary of fluorescence properties of HOClSense dyes**

| Name | Before HOCl | | After HOCl | | Bathochromic shift in emission | Final Wavelength Fluorescence Increase/ Decrease | Fluorescence Parameters | Peak-to-Peak Fluorescence Increase/ Decrease |
|---|---|---|---|---|---|---|---|---|
| | λex (nm) | λem (nm) | λex (nm) | λem (nm) | | | | |
| **1a** | 505 | 512 | 535 | 546 | 34 nm | ↓ 33% Decrease, 1.5-Fold | (Ex505, Em545/ Ex535, Em545) | ↓ 85% Decrease, 6.5-Fold |
| **1b** | 507 | 525 | 530 | 555 | 30 nm | ↑ 1550% Increase, 17-Fold | (Ex507, Em555/ Ex530, Em555) | ↑ 641% Increase, 7.4-Fold |
| **1c** | 529 | 541 | 529* | 559* | 18 nm | ↓ 99% Decrease, 90-Fold | (Ex529, Em541/ Ex529, Em559) | ↓ 99% Decrease, 256-Fold |
| **1d** | 535 | 557 | 300* | 467* | 90 nm | ↓ 66% Decrease, 2.9-Fold | (Ex535, Em557/ Ex300, Em557) | ↓ 94% Decrease, 16-Fold |
| **1e** | 512 | 528 | 525 | 541 | 13 nm | ↓ 98% Decrease, 40-Fold | (Ex512, Em541/ Ex525, Em541) | ↓ 98% Decrease, 52-Fold |
| **1f** | 520 | 541 | 535 | 560 | 19 nm | ↑ 435% Increase, 5.4-Fold | (Ex520, Em545/ Ex535, Em560) | ↑ 296% Increase, 4-Fold |
| **2a** | 503 | 515 | 520 | 543 | 28 nm | ↓ 16% Decrease, 1.1-Fold | (Ex503, Em543/ Ex520, Em543) | ↓ 76% Decrease, 4-Fold |
| **2b** | 520 | 537 | 532 | 555 | 18 nm | ↑ 81% Increase, 1.8-Fold | (Ex520, Em555/ Ex532, Em555) | ↑ 50% Increase, 1.5-Fold |
| **3a** | 501 | 514 | 529 | 547 | 33 nm | ↓ 52% Decrease, 1.5-Fold | (Ex501, Em547/ Ex529, Em547) | ↓ 85% Decrease, 6.5-Fold |
| **3b** | 508 | 594 | 531 | 598 | 4 nm | ↑ 250% Increase, 3.5-Fold | (Ex508, Em598/ Ex531, Em598) | ↑ 268% Increase, 3.7-Fold |

* Peak maxima are estimated due to low fluorescence intensities from quenching.

1a

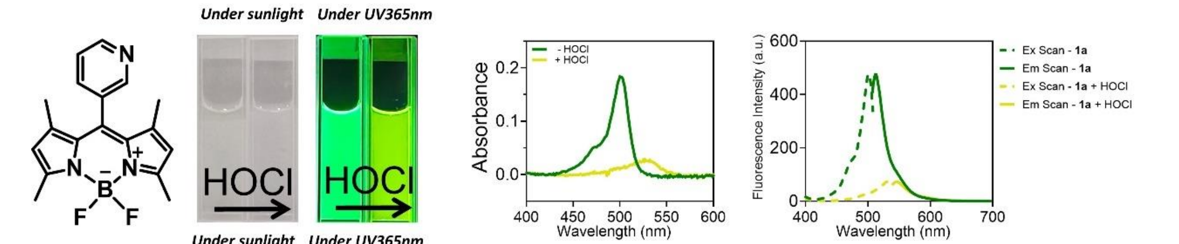


1b

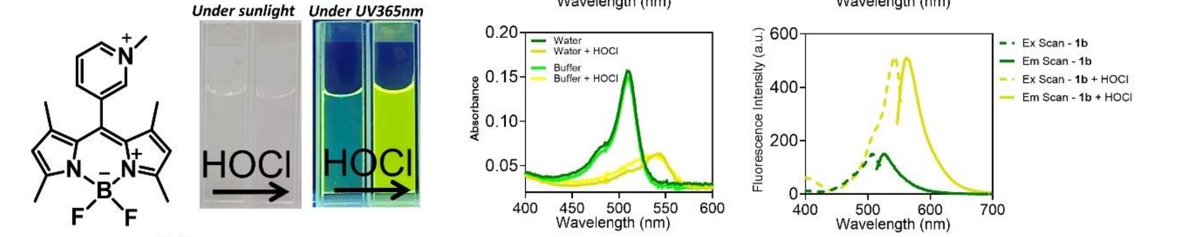


1b-azide

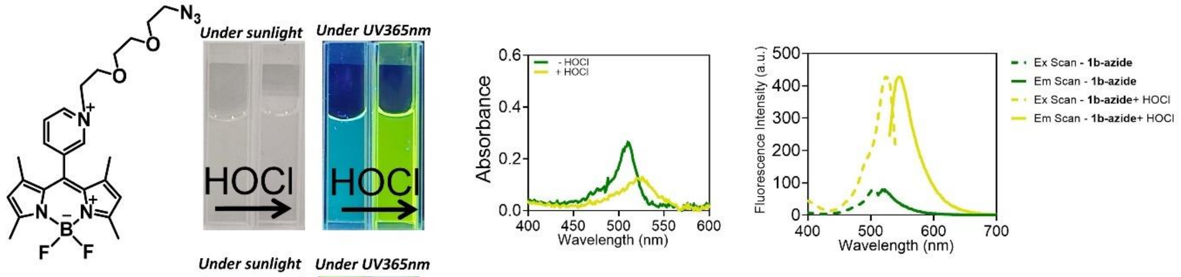


1c

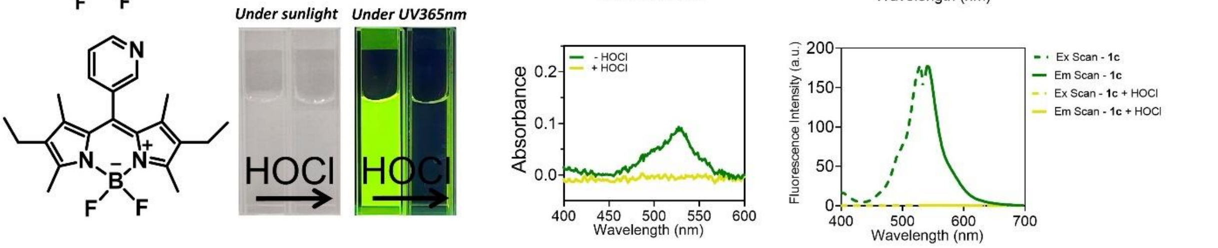


1d

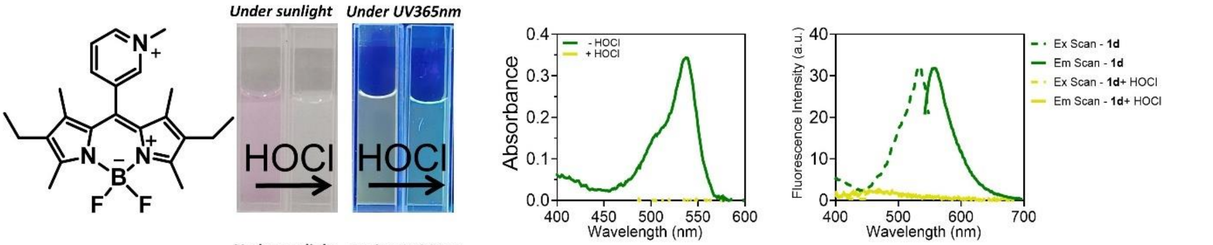


1e

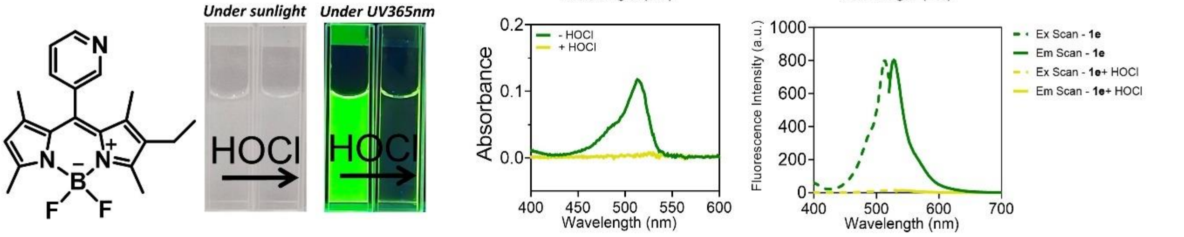


1f

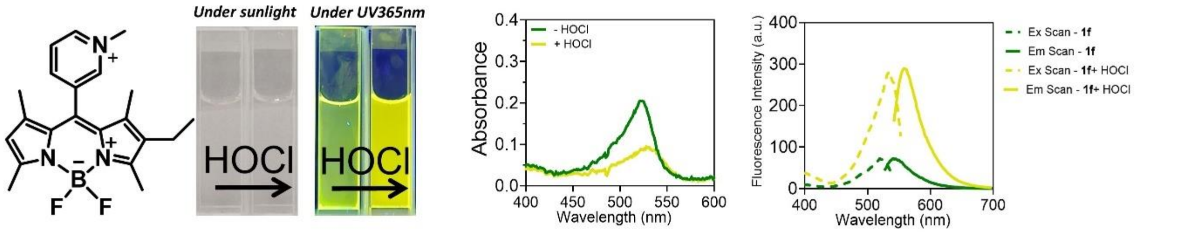

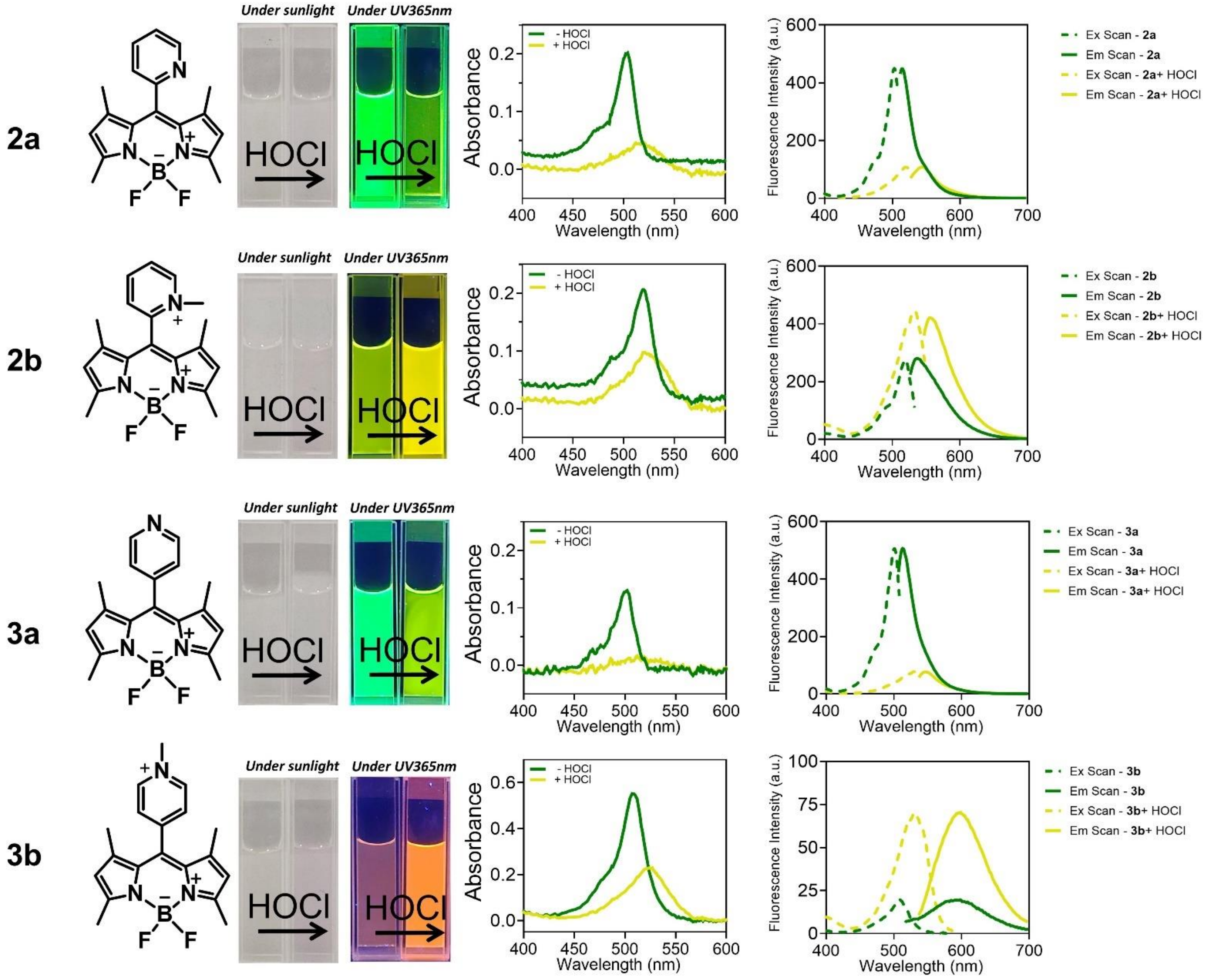


**Figure S40. Spectroscopic measurement of HOClSense dyes. a**, Chemical structure of HOClSense dyes. **b**, HOClSense dyes in the absence and the presence of 10 equivalent of HOCl under irradiation of UV light at 365 nm. **c**, UV-Vis spectra and **d**, fluorescence excitation scans and emission scans of HOClSense dyes in the absence and the presence of 10 equivalent of HOCl. Conditions: 5 µM of HOClSense dyes in water containing 0.5% DMF.

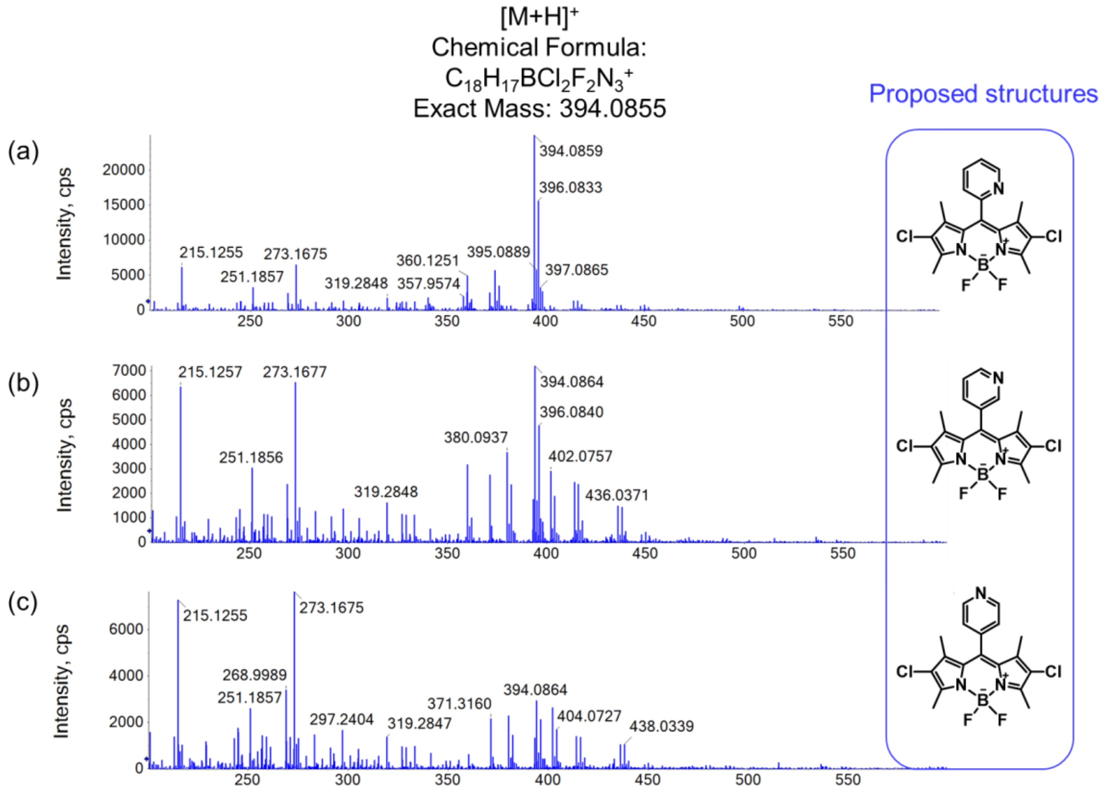


**Figure S41.** HRMS (+ve ESI) of the reaction mixtures of HOClSense dyes (**1a**, **2a** and **3a**) with HOCl (10 equiv.). The peak with *m/z* = 394.09 corresponds to the di-chlorinated product.

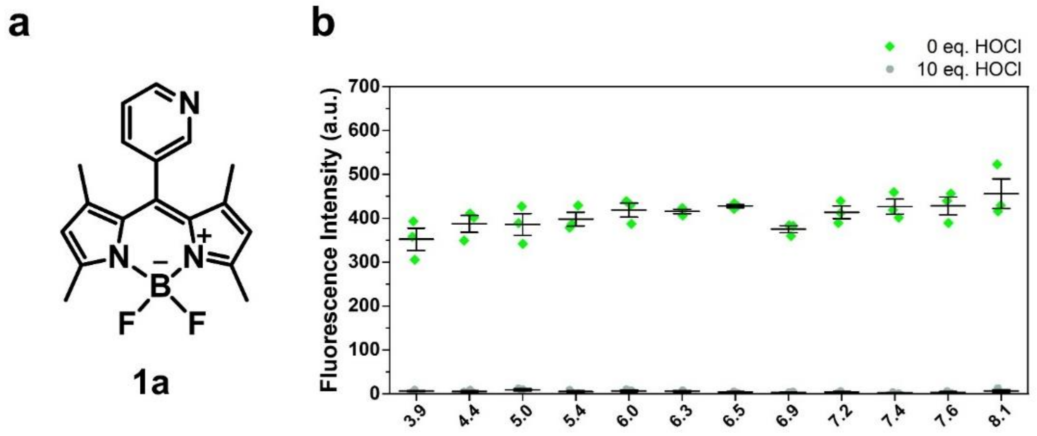


**Figure S42. pH response profile of 1a. a**, Chemical structure of **1a**. **b**, Fluorescence intensities of **1a** (5 µM) at 513 nm in the absence or presence of HOCl (10 equiv.) at different pH values. Conditions: citrate-phosphate buffer (10 mM) containing 0.5 % DMF. Error bars indicate the mean ± standard error of the mean (s.e.m.) of three independent measurements.

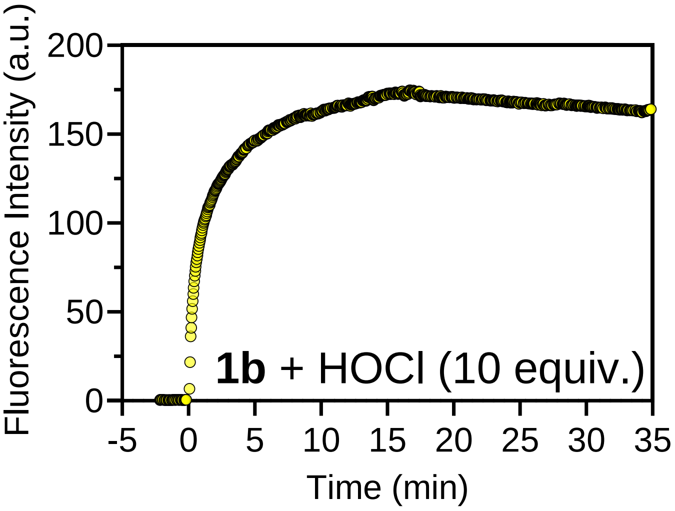


**Figure S43. Time-dependent fluorescence response of 1b (5 μM) in the presence of HOCl (10 equiv.).** Conditions: 60 mM sodium phosphate buffer, pH 7.15, containing 0.5% DMF.

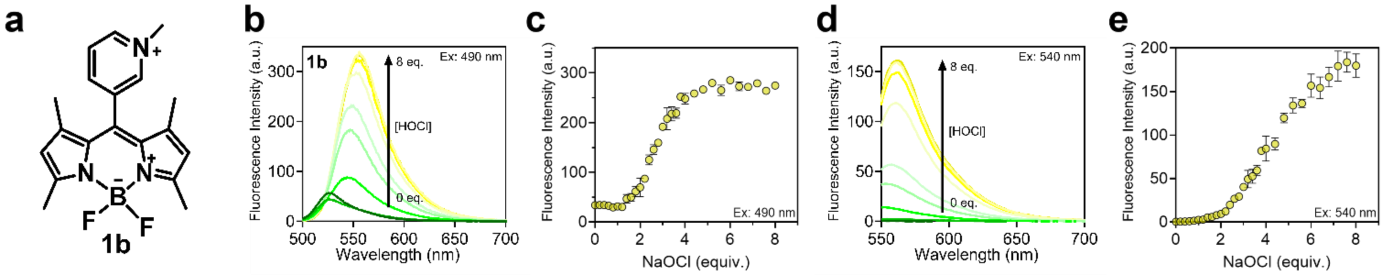


**Figure S44. Dose-dependent fluorescence response of 1b to HOCl. a**, Chemical structure of **1b**. **b**, **d**, **F**luorescence spectra and **c**, **e**, fluorescence intensities of **1b** (5 μM) in the presence of HOCl (0–8 equiv.) upon excitation at **b–c**, $\lambda_{ex}$ = 490 nm, **d–e** $\lambda_{ex}$ = 540 nm. Conditions: water containing 0.5% DMF. Error bars indicate the mean ± standard error of mean (s.e.m.) of three independent measurements.

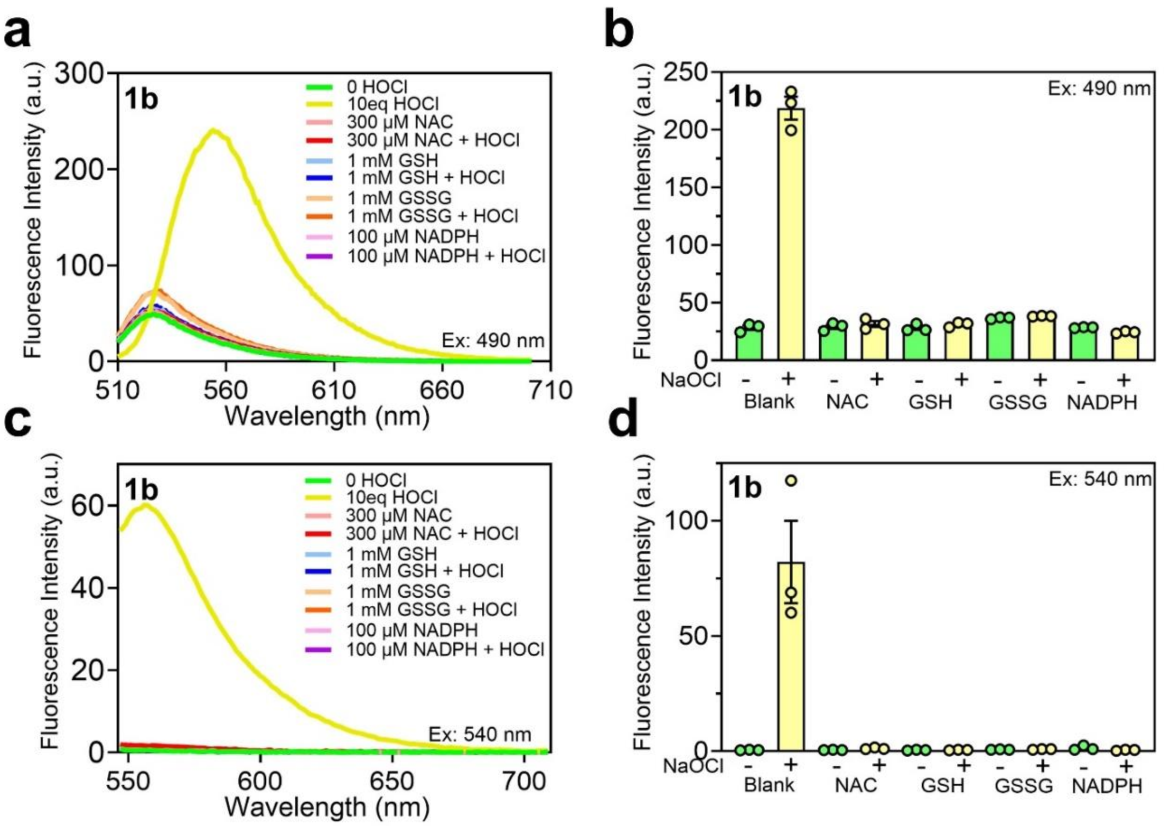


**Figure S45. Interference of ROS scavengers in HOCl detection**. **a**, **c**, Fluorescence spectra and **b**, **d**, fluorescence intensities at 560 nm of **1b** (5 µM) in the presence of various ROS scavengers (300 µM *N*-Acetyl Cysteine, 1 mM GSH, 1mM GSSG and 100 µM NADPH) with and without HOCl under excitation at **a**, **b**, $\lambda_{ex}$ = 490 nm and **c**, **d**, $\lambda_{ex}$ = 540 nm. Conditions: water containing 0.5% DMF. Error bars indicate the mean ± standard error of mean (s.e.m.) of three independent measurements.

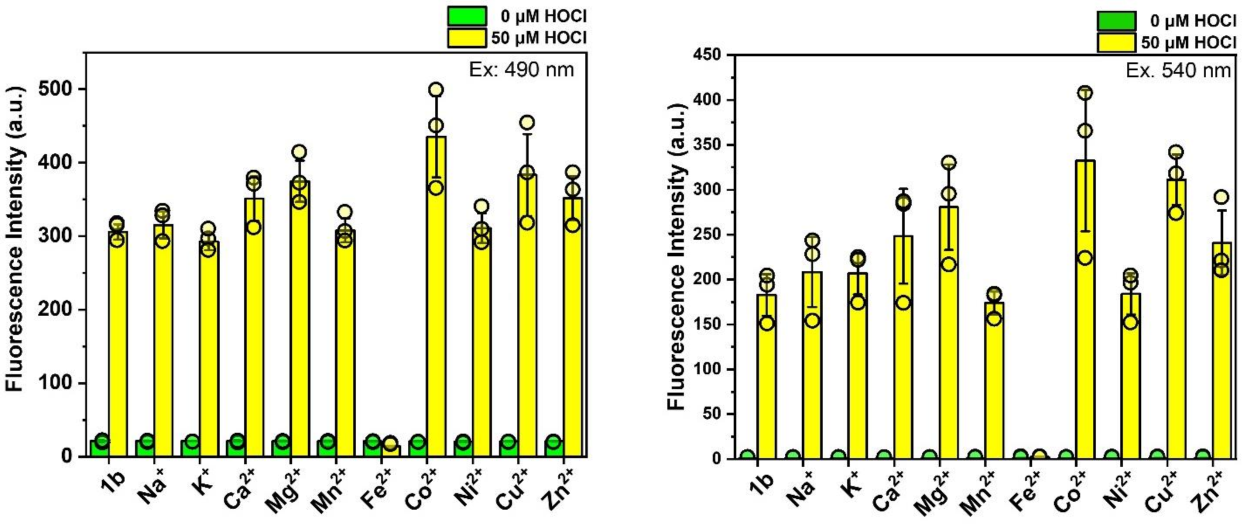


**Figure S46**. **Metal ion sensitivity of 1b.** Fluorescence intensity of **1b** (5 µM) in the presence of different metal ion species (20 µM for $Mn^{2+}$, $Fe^{2+}$, $Co^{2+}$, $Ni^{2+}$, $Cu^{2+}$, $Zn^{2+}$, and 100 mM for $Na^{+}$, $K^{+}$, $Ca^{2+}$, $Mg^{2+}$) with and without 50 µM HOCl. Conditions: water containing 0.5% DMF. **a**, $\lambda_{ex}$ = 490 nm. **b**, $\lambda_{ex}$ = 540 nm. Error bars indicate the mean ± standard error of mean (s.e.m.) of three independent measurements.

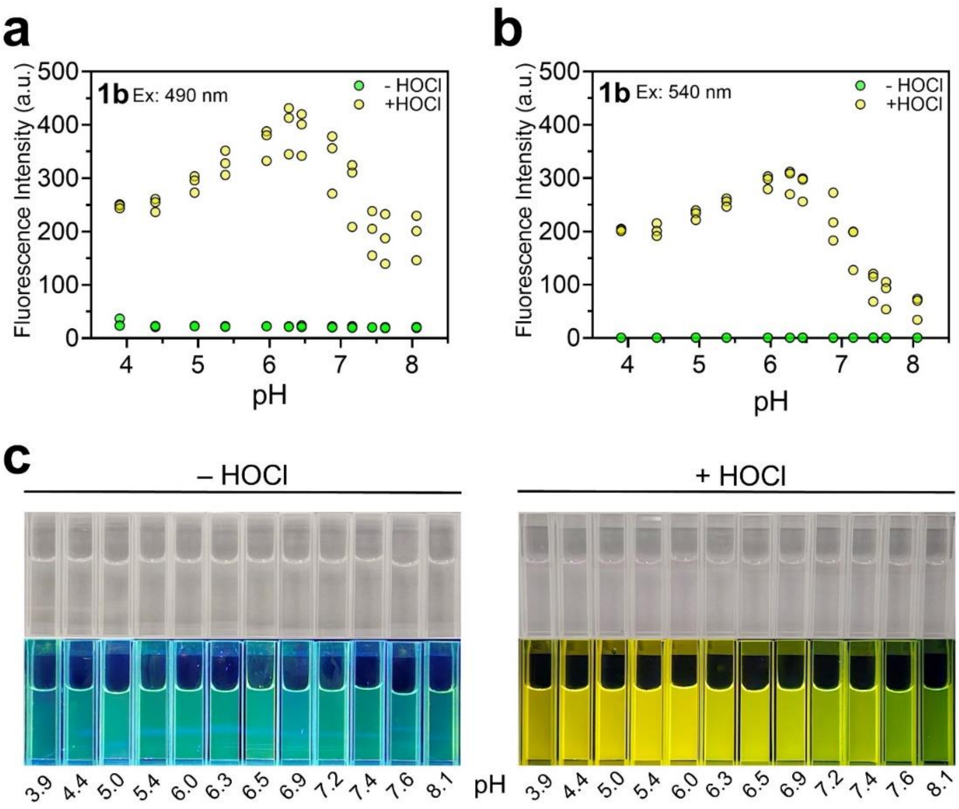


**Figure S47**. **pH response profiles of 1b.** Fluorescence intensities of **1b** (5 μM) at 560 nm in the presence and absence of HOCl (10 equiv.) at different pH upon excitation at **a**, $\lambda_{ex}$ = 490 nm and **b**, $\lambda_{ex}$ = 540 nm. Conditions: citrate-phosphate buffer (10 mM) containing 0.5 % DMF, n = 3. **c**, Quartz cuvette photos of **1b** (5 μM) in the presence and absence of HOCl (10 equiv.) at different pH values under room lighting and under UV (365 nm) illumination.

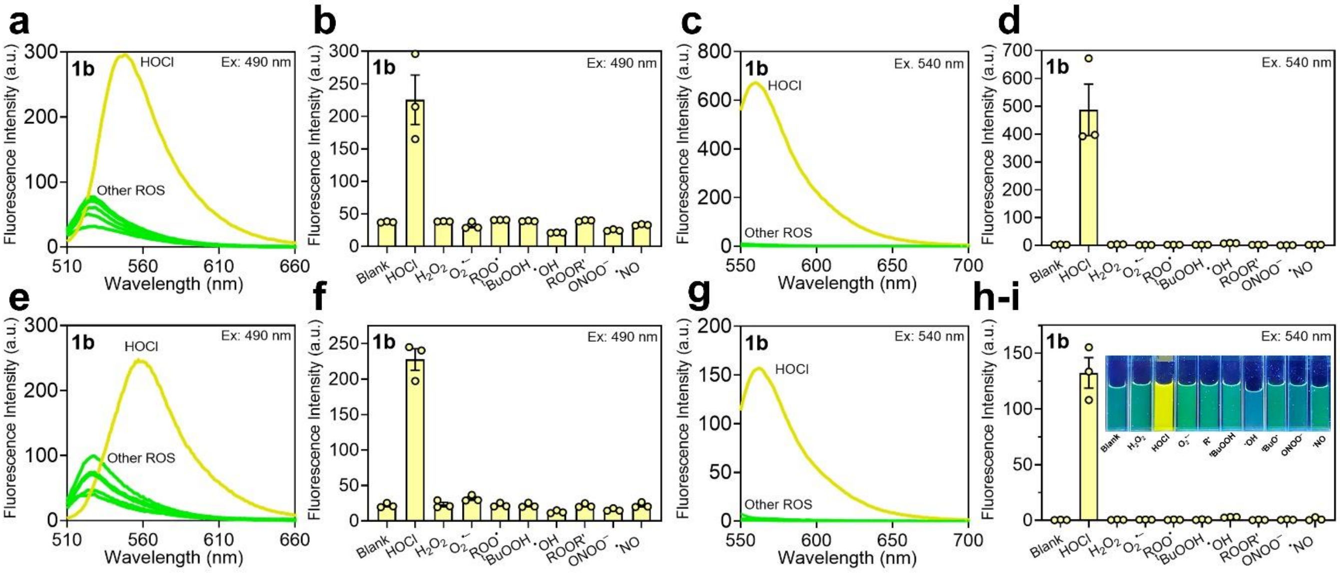


**Figure S48**. **Selectivity profiles of 1b. a**, **c**, **e**, **g**, Fluorescence spectra and **b**, **d**, **f**, **h**, fluorescence intensities of **1b** (5 μM) in the presence of different ROS/ RNS (10 equiv.) under excitation at **a–b** & **e–f**, $\lambda_{ex}$ = 490 nm, and **c–d** & **g–h**, $\lambda_{ex}$ = 540 nm in **a–d** water and **e–i** phosphate buffer (60 mM, pH 7.2). Photographs of **1b** (5 μM) in the presence of different ROS/ RNS (10 equiv.) in quartz cuvettes under UV (365 nm) illumination. Error bars indicate the mean ± standard error of mean (s.e.m.) of three independent measurements.

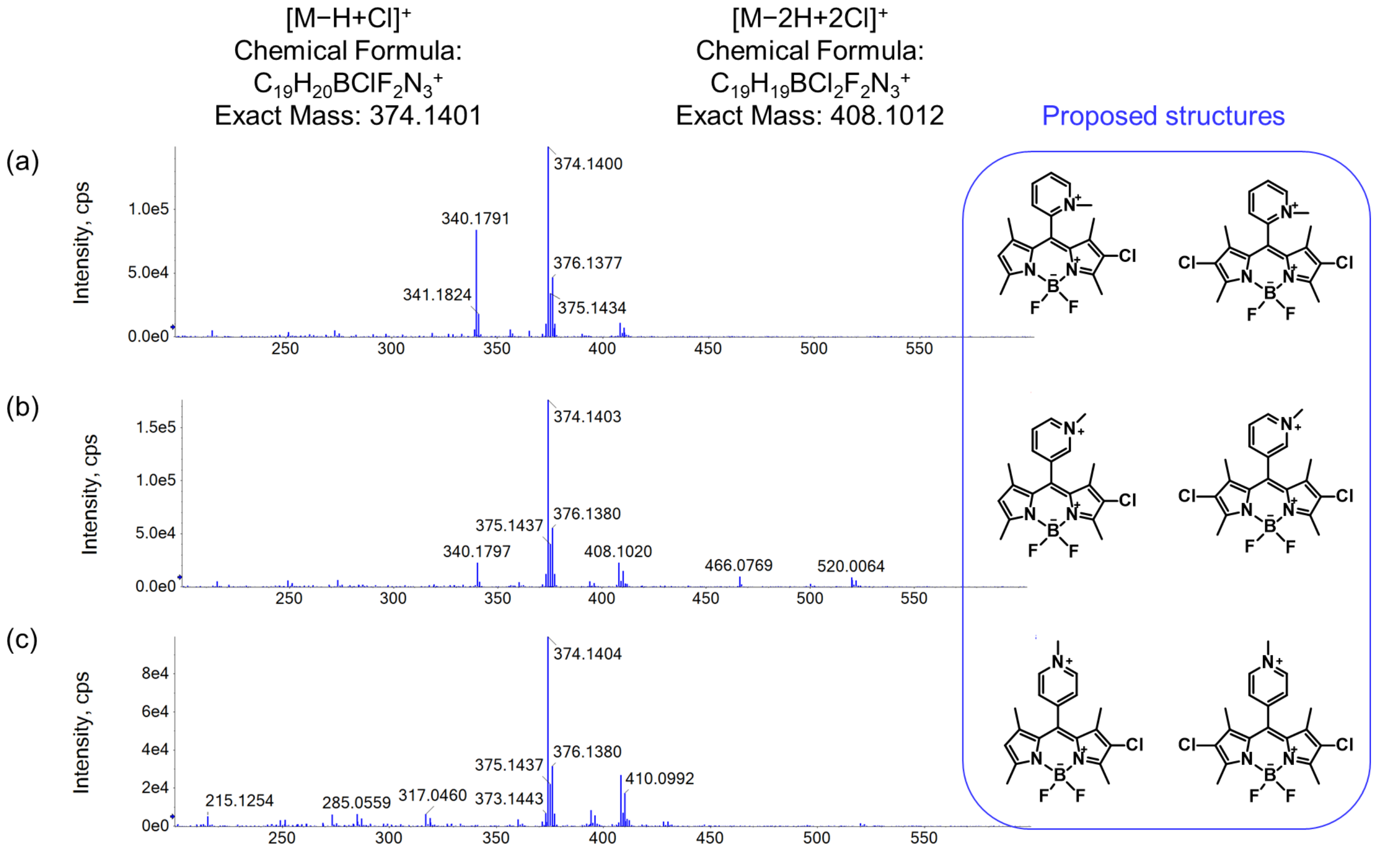


**Figure S49.** HRMS (+ve ESI) of the reaction mixtures of **1b**, **2b**, and **3b** with HOCl (10 equiv.). The peak with *m/z* = 374.14 corresponds to the mono-chlorinated product and the peak with *m/z* = 408.10 corresponds to the di-chlorinated product.

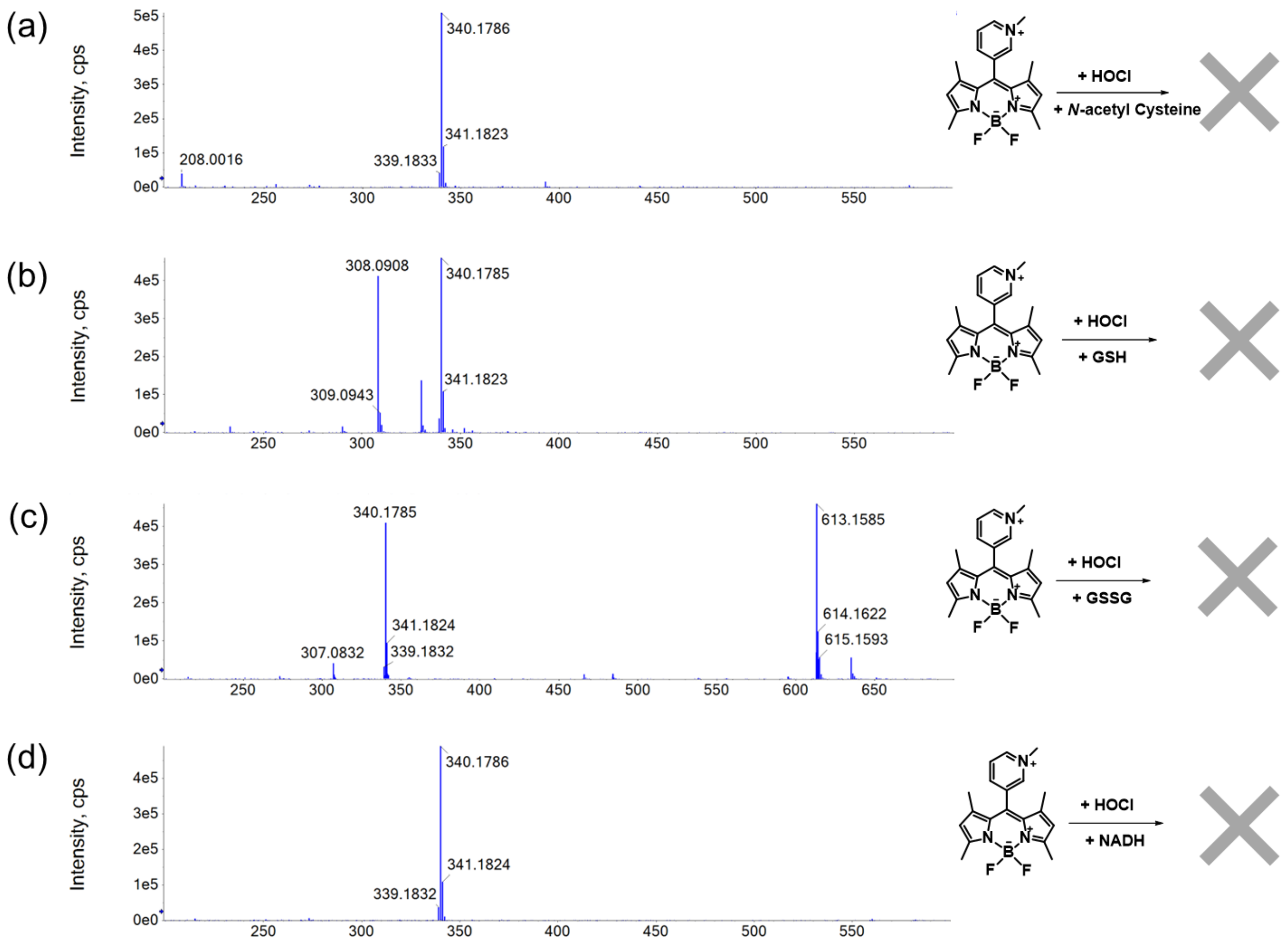


**Figure S50**. HRMS (+ve ESI) of the reaction mixtures of 1b with HOCl (10 equiv.) in the presence of different ROS scavengers. **a**, *N*-Acetyl Cysteine (300 μM). **b**, GSH (1 mM). **c**, GSSG (1 mM). **d**, NADPH (100 μM). The peak with *m/z* = 340.18 corresponds to the unreacted **1b** molecules.

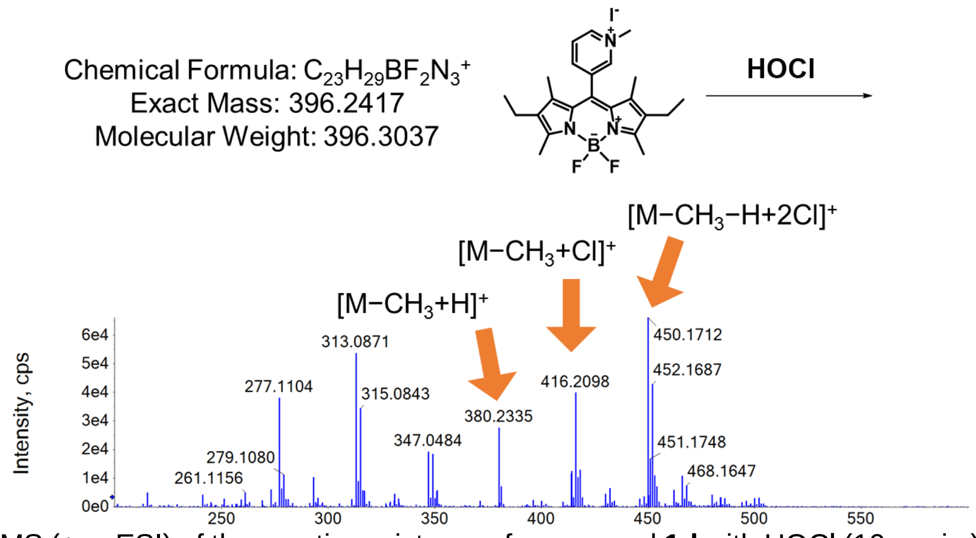


**Figure S51**. HRMS (+ve ESI) of the reaction mixtures of compound **1d** with HOCl (10 equiv.). Decomposition of **1d** occurred including demethylation and chlorination on other positions.

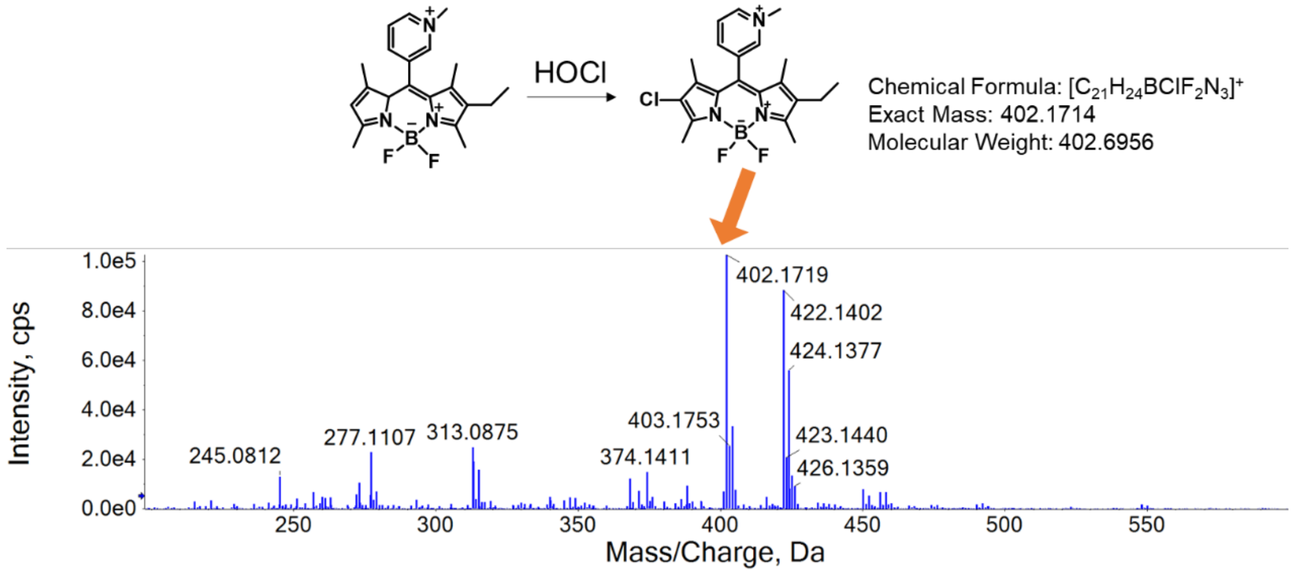


**Figure S52**. HRMS (+ve ESI) of the reaction mixtures of compound **1f** with HOCl (10 equiv.). The peak with *m/z* = 402.1719 corresponds to the mono-chlorinated product.

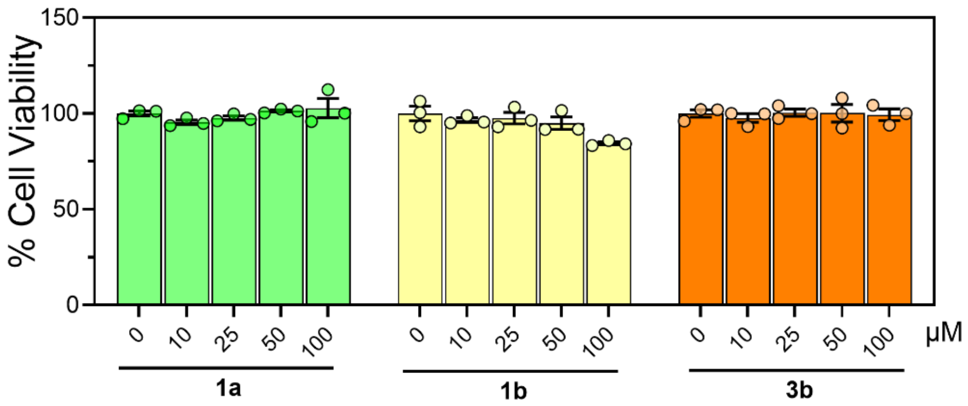


**Figure S53.** Cell viability of RAW 264.7 cells upon labelling of **1a**, **1b** and **3b** at different concentrations (0 µM, 10 µM, 25 µM, 50 µM, and 100 µM). Error bars indicate the mean ± standard error of the mean (s.e.m.) of three independent measurements.

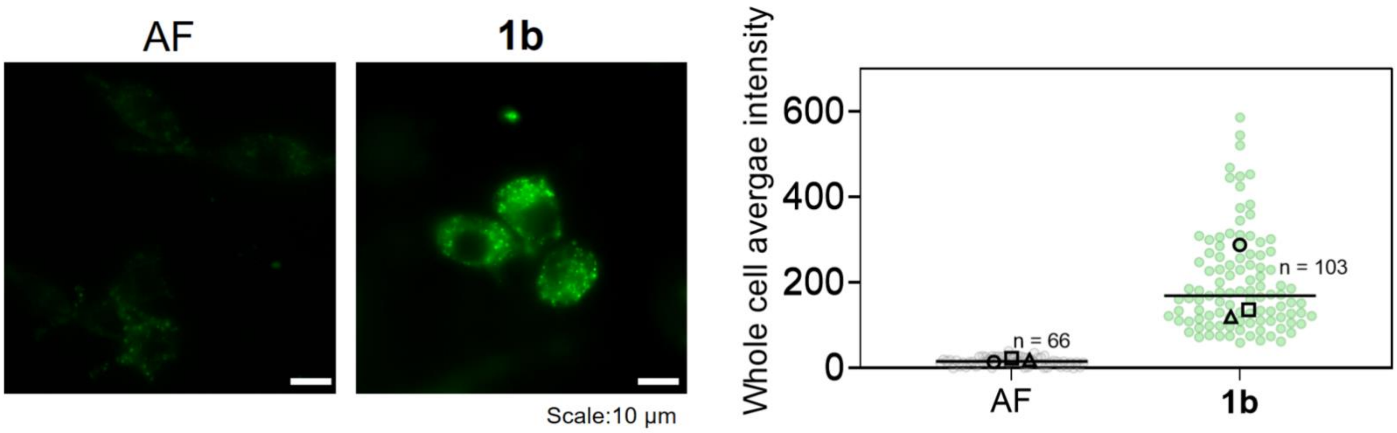


**Figure S54. Cellular uptake of 1b.** Representative fluorescence images of RAW 264.7 cells labelled with **1b** or autofluorescence. Quantification of whole cell average intensity. Experiments were performed in triplicate. The median value of each trial is given by a square, circle, and triangle symbol (n = number of cells).

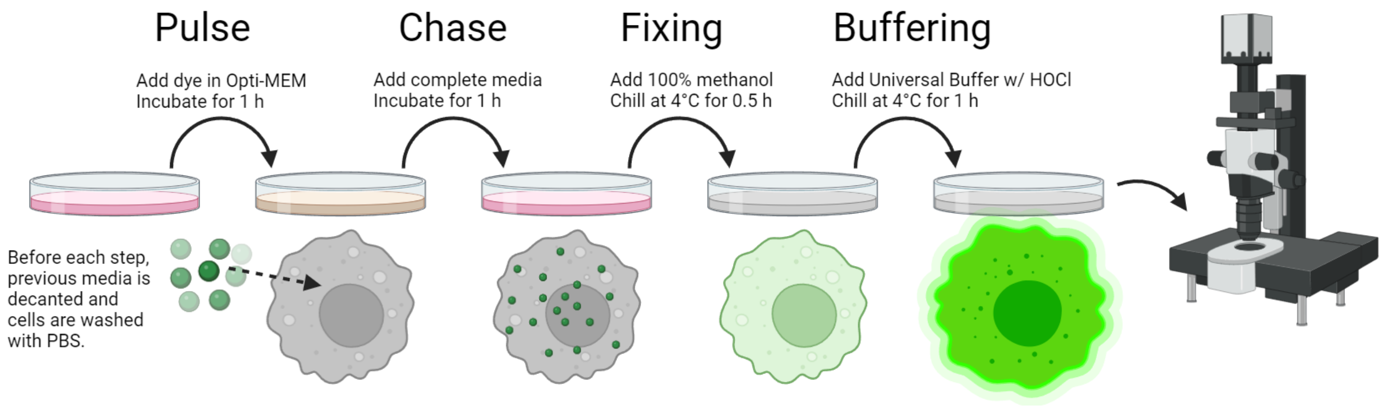


**Figure S55. Schematic diagram showing the protocol of HOCl clamping experiments.** The cells were pulsed with the dyes for 1 h, chased for 1 h, fixed with 4°C methanol for 30 min, and then clamped at the indicated [HOCl] overnight.

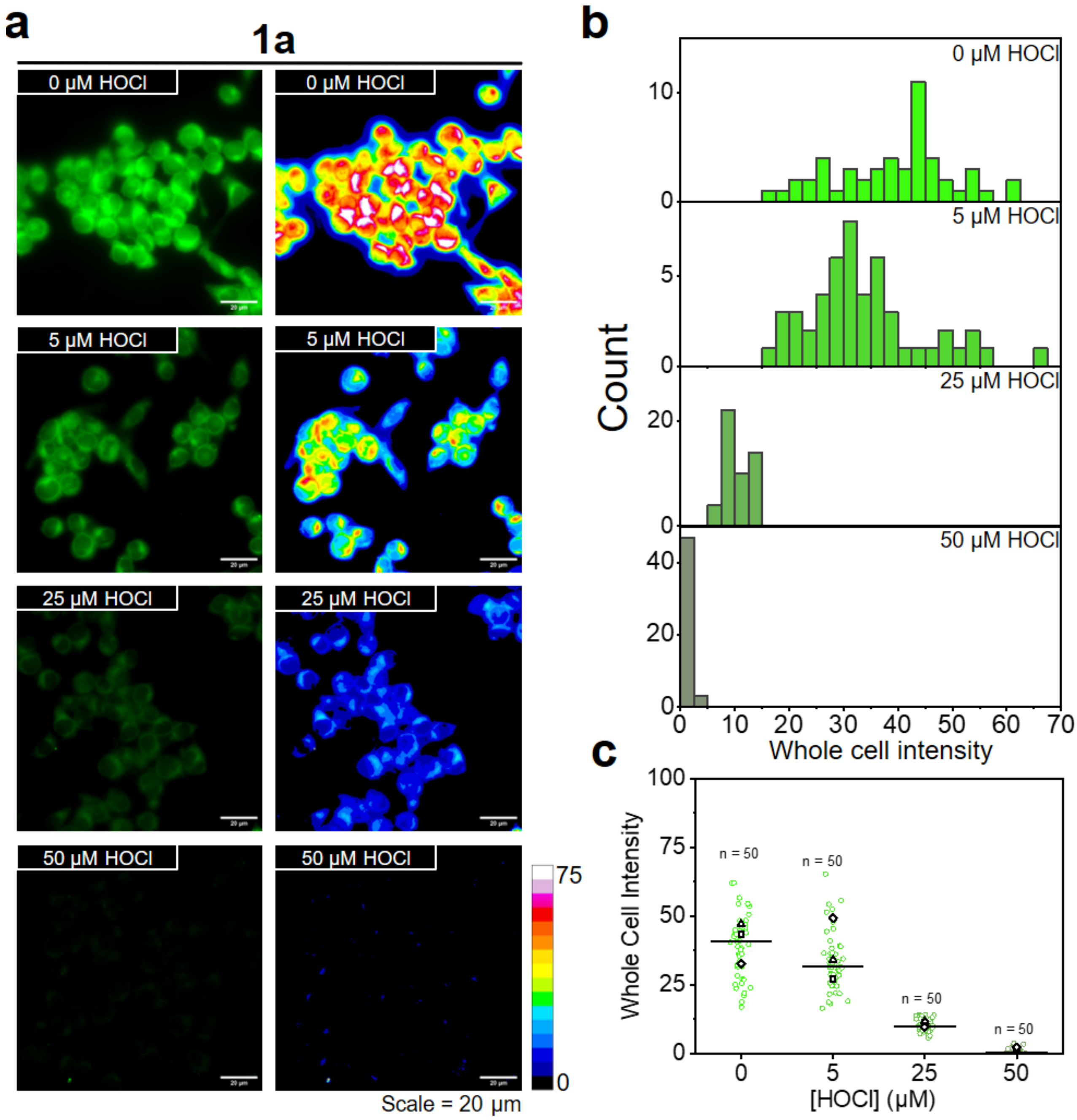


**Figure S56. Intracellular HOCl calibration of 1a**. **a**, Representative fluorescence images of **1a**-labelled RAW 264.7 cells clamped at the indicated [HOCl]. **b**, Histograms and **c**, Scatter plot of the whole cell intensities at the indicated [HOCl]. Experiments were performed in triplicate. The median of all trials is given by a line. The mean value of individual trial is given by a square, triangle, and diamond symbol (n = number of cells).

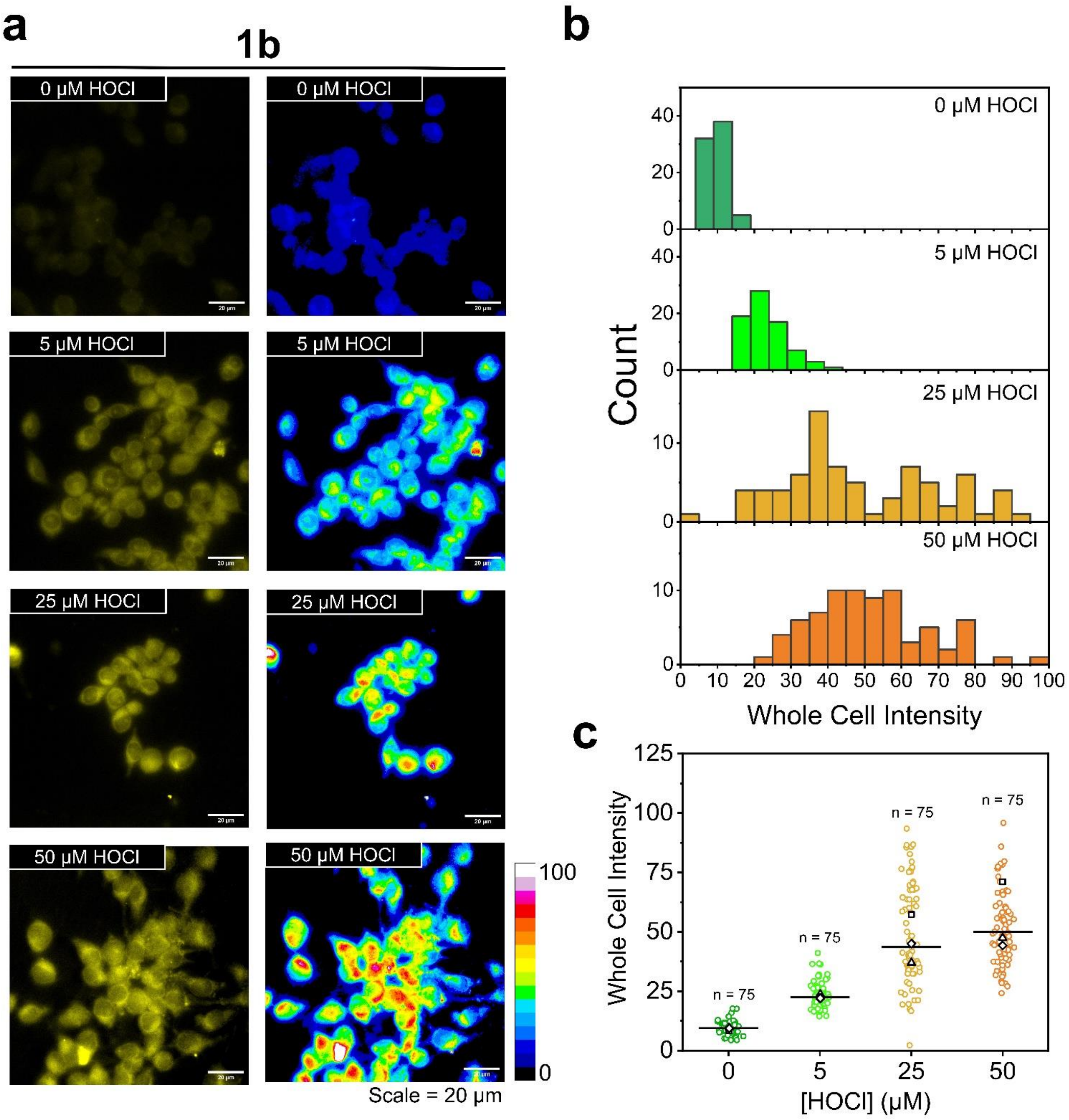


**Figure S57. Intracellular HOCl calibration of 1b**. **a**, Representative fluorescence images of **1b**-labelled RAW 264.7 cells clamped at the indicated [HOCl]. **b**, Histograms and **c**, Scatter plot of the whole cell intensities at the indicated [HOCl]. Experiments were performed in triplicate. The median of all trials is given by a line. The mean value of individual trial is given by a square, triangle, and diamond symbol (n = number of cells).

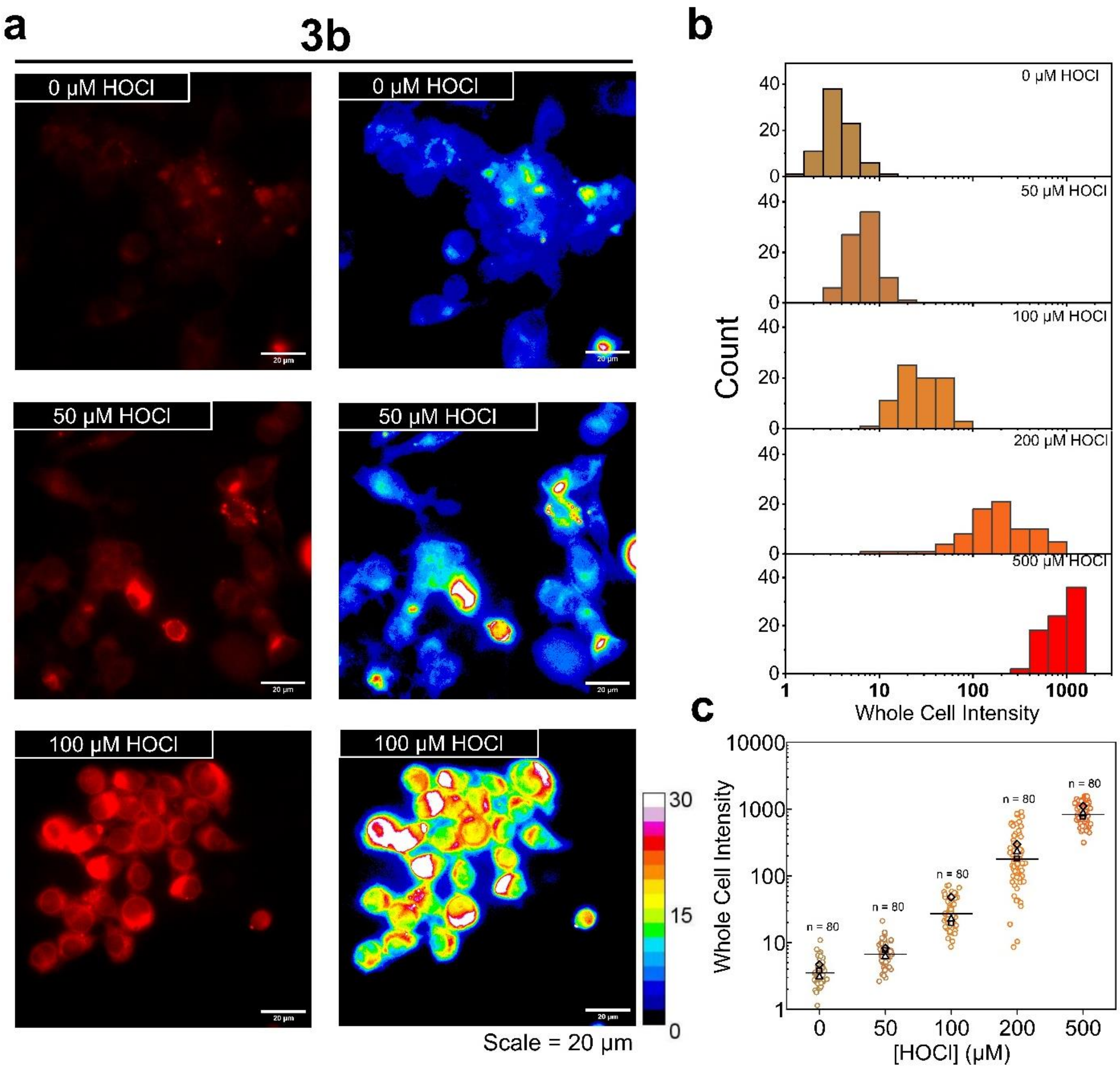


**Figure S58. Intracellular HOCl calibration of 3b**. **a**, Representative fluorescence images of **3b**-labelled RAW 264.7 cells clamped at the indicated [HOCl]. **b**, Histograms and **c**, Scatter plot of the whole cell intensities at the indicated [HOCl]. Experiments were performed in triplicate. The median of all trials is given by a line. The mean value of individual trial is given by a square, triangle, and diamond symbol (n = number of cells).

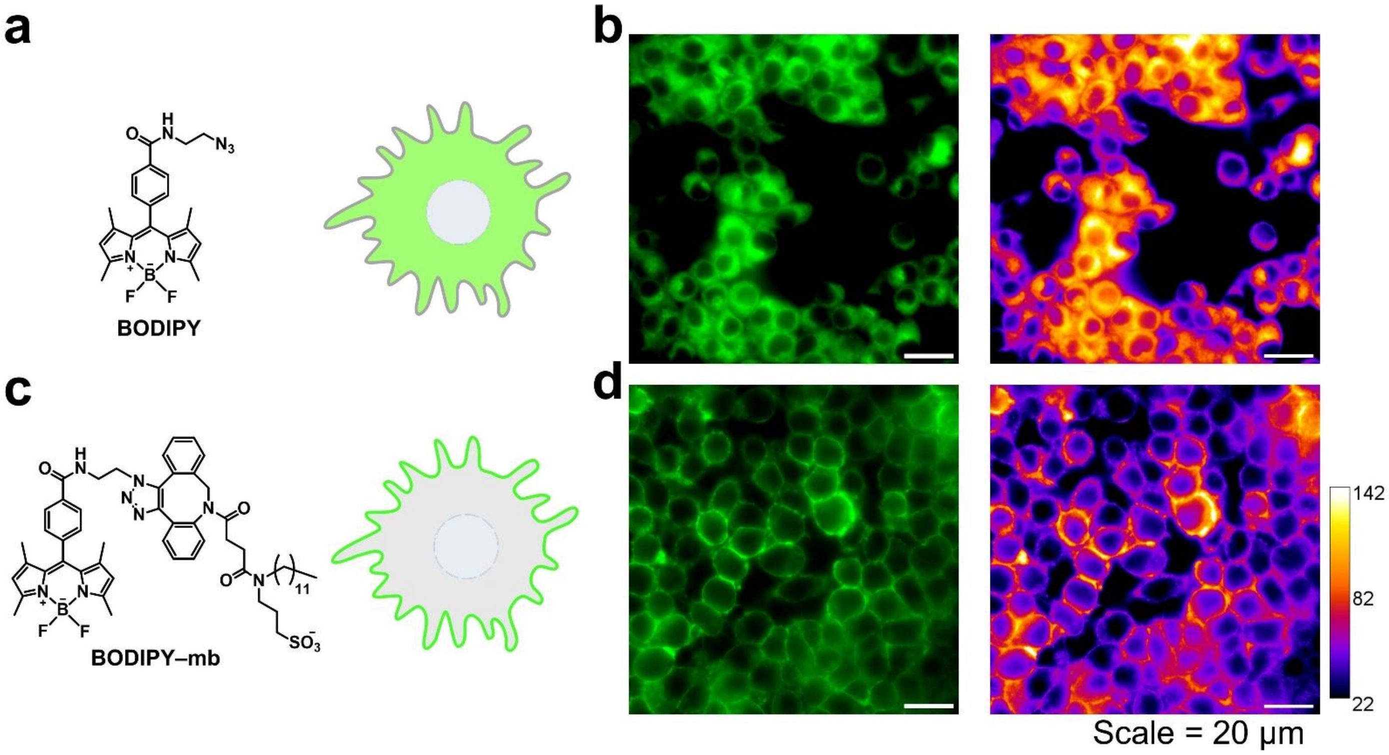


**Figure S59. Selective plasma membrane staining by membrane targeting ligand. a**, Chemical structure of **BODIPY-azide**. **b**, Representative fluorescence images of **BODIPY-azide** labelling whole cells. By conjugating **BODIPY-azide** with a membrane-targeting ligand, **BODIPY-mb** selectively labels the plasma membrane of the cells. **c**, Chemical structure of **BODIPY-mb**. **d**, Representative fluorescence images showing **BODIPY-mb** labelling the plasma membrane of the cells.

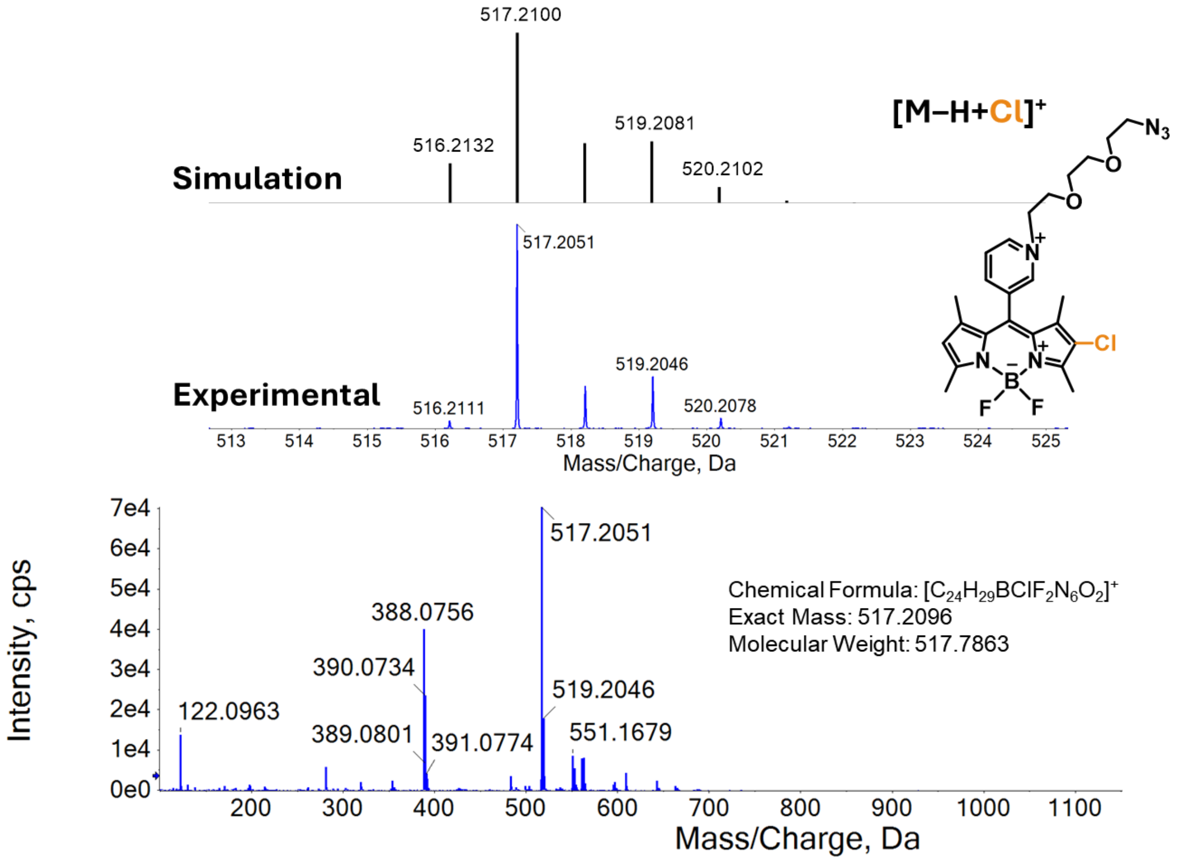


**Figure S60**. HRMS (+ve ESI) of the reaction mixtures of **1b-azide** with HOCl (10 equiv.). The peak with *m/z* = 517.2051 corresponds to the protonated mono-chlorinated product $[M–H+Cl]^+$ ion.

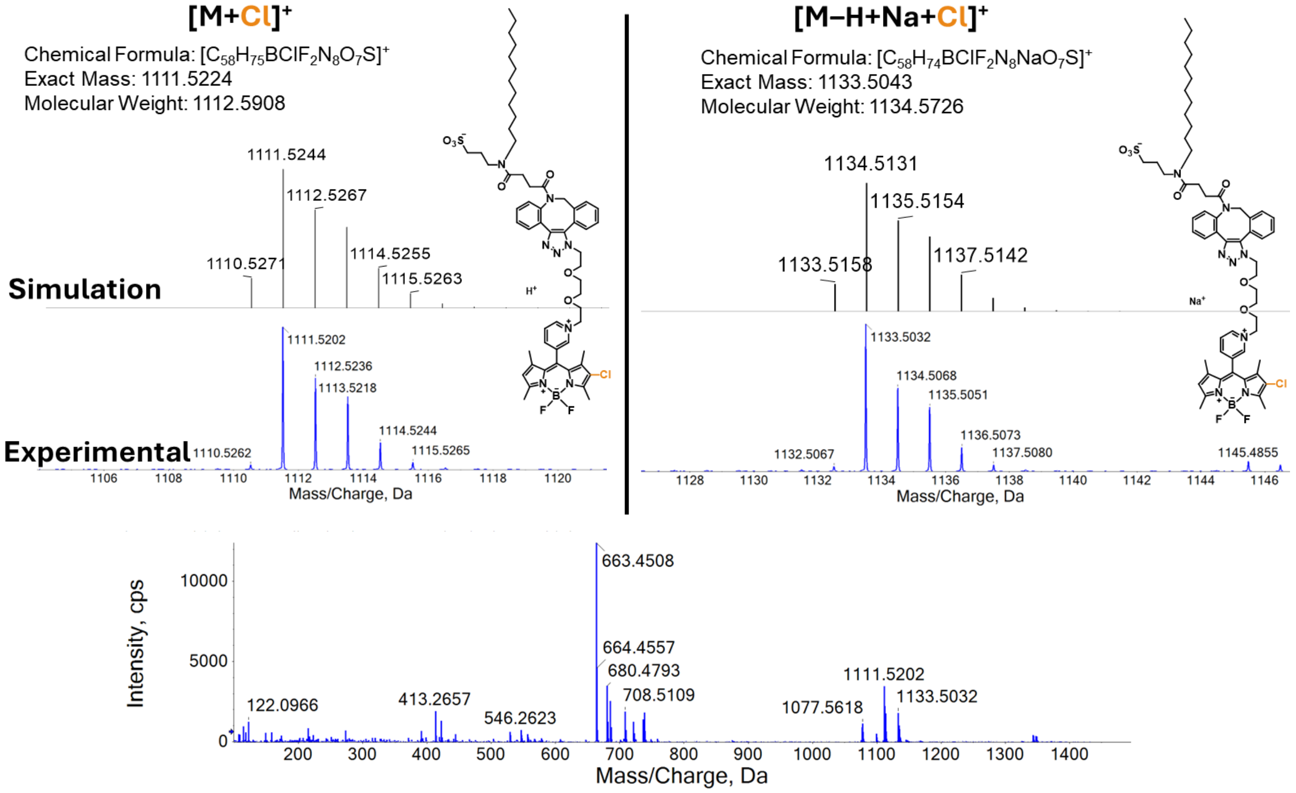


**Figure S61**. HRMS (+ve ESI) of the reaction mixtures of **1b-mb** with HOCl (10 equiv.). The peak with *m/z* = 1111.5202 corresponds to the protonated mono-chlorinated product $[M+Cl]^+$ ion and the peak with *m/z* = 1133.5032 corresponds to the mono-chlorinated product with sodium $[M–H+Na+Cl]^+$ ion.

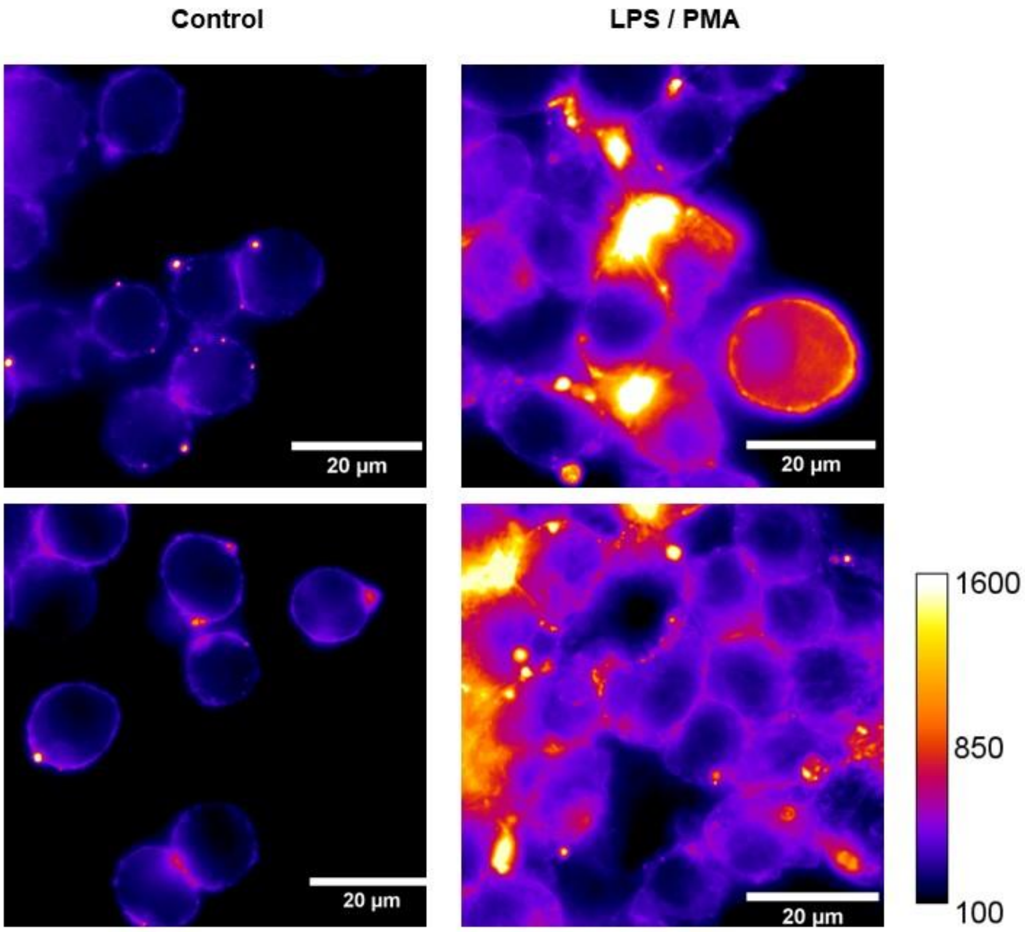


**Figure S62. 1b-mb detects the PAMPs-induced extracellular HOCl.** Representative fluorescence images of **1b-mb**-labelled RAW 264.7 cells showing the relative fluorescence intensity increase upon stimulation of 80 ng $mL^{-1}$ lipopolysaccharides (LPS) and 10 ng $mL^{-1}$ phorbol 12-myristate 13-acetate (PMA).
