## Supporting information_synthesis for "Leveraging Chlorination-Based Mechanism for Resolving Subcellular Hypochlorous Acid"

### General synthetic materials and methods:

Chemicals or reagents were purchased from Fisher Chemicals, Sigma Aldrich, TCI, Thermo Fisher, Ambeed or Alfa Aesar and used as received. All solvents were used directly without further treatment or distillation. Silica gel 60 (70–230 mesh, Supelco®) was used for column chromatography. Thin Layer Chromatography (TLC) was performed using $F_{254}$ silica (aluminium sheet back plates, Supelco®).

**NMR spectroscopy.** NMR spectra were recorded from a Bruker Advance–III 400 NMR spectrometer and a JEOL 400 MHz nuclear magnetic resonance (NMR) spectrometer (Department of Chemistry, St. Lawrence University NY), which are operating at 400 MHz for $^{1}H$ and 101 MHz for $^{13}C\{^{1}H\}$, respectively. Chemical shifts are quoted in ppm. $^{1}H$ and $^{13}C$ chemical shifts were referenced internally with solvent residue chemical shift values ($CDCl_3$: $^{1}H$, 7.26 ppm; $^{13}C$, 77.16 ppm; $CD_3CN$: $^{1}H$, 1.94 ppm; $^{13}C$, 1.32 ppm). NMR data were processed using MestReNova Software (Mestrelab). Coupling constants are reported in Hertz.

**Mass spectrometry.** High-resolution mass spectra (HRMS) were recorded using a SCIEX X500B QTOF mass spectrometer (at Center for Air and Aquatic Resources Engineering and Sciences, CAARES, Clarkson university, NY) which operated in positive ion mode (+ve ESI).

**Anion exchange to form $M^+PF_6^-$ salts.**
The cationic BODIPY-based HOClSense dyes bearing iodide anion was dissolved and sonicated in a small amount of MeCN (10 mL to 20 mL) and diluted with water to a final volume of 300−400 mL. Then excess $NH_4PF_6$ was added into the solution and mixture was allowed to decant overnight. The precipitate was collected by filtration, washed with water, and dried in air. The mass spectra of $M^+PF_6^-$ salts was consistent with the $[M]^+$ peaks.

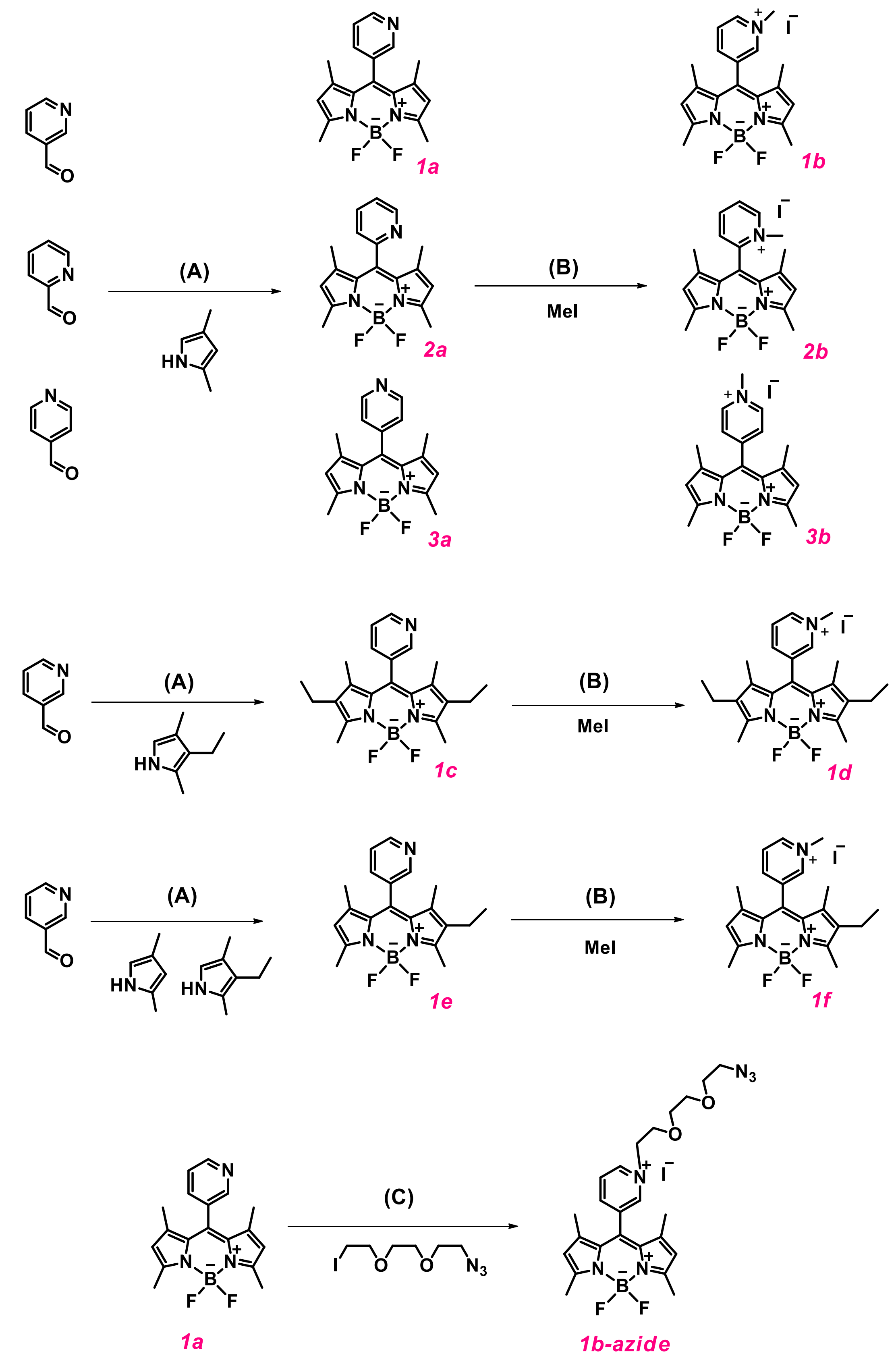


**Scheme S1.** Synthetic scheme of HOClSense dyes. (**A**) (i) Pyrrole, trifluoroacetic acid, room temperature, 24 h (ii) DDQ, 4 h (iii) $NEt_3$ and $BF_3{\cdot}OEt_2$, 24 h; (**B**) Iodomethane, reflux, 24 h; (**C**) Azide-PEG3-iodide, reflux, 24 h.

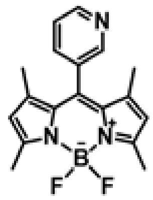

Preparation of **1a**:

The synthesis of BODIPY fluorophores was modified from the reported literature.[1,2] In brief, 3-pyridine carboxaldehyde (0.42 g, 3.90 mmol) and 2,4-dimethylpyrrole (0.75 g, 7.84 mmol) were dissolved in 500 mL of $CH_2Cl_2$, and the solution was degassed with argon for 15 min. Two drops of trifluoroacetic acid were added and the mixture was stirred at room temperature for 24 h in the dark. 2,3-Dichloro-5,6-dicyano-1,4-benzoquinone (1.16 g, 5.11 mmol) was added to the mixture and allowed to react for 4 h at room temperature. Then 6.5 mL of triethylamine was added to the mixture, followed by 6.5 mL of $BF_3$•$OEt_2$. The mixture was allowed to stir overnight. The organic layer was washed with $H_2O$ (300 mL) three times. The organic layer was concentrated by rotary evaporator and the product was purified by silica gel column chromatography using ethyl acetate/hexanes (*v/v* = 1:2) to give **1a** as an orange solid. Yield: 390 mg, 31 %. $^1H$ NMR (400 MHz, $CDCl_3$, 298 K) δ 8.76 (d, J = 4.1 Hz, 1H), 8.57 (s, 1H), 7.68 (dt, J = 7.7, 1.7 Hz, 1H), 7.50 (dd, J = 7.6, 5.0 Hz, 1H), 6.00 (s, 2H), 2.54 (s, 6H), 1.35 (s, 6H). $^{13}C$ NMR (101 MHz, $CDCl_3$, 298 K) δ 156.47, 150.10, 148.39, 142.84, 137.21, 136.29, 131.55, 123.93, 121.89, 15.10, 14.73. HRMS (+ve ESI): calculated for $C_{18}H_{19}BF_2N_3$ $[M+H]^+$ *m/z* 326.1635., found 326.1623.

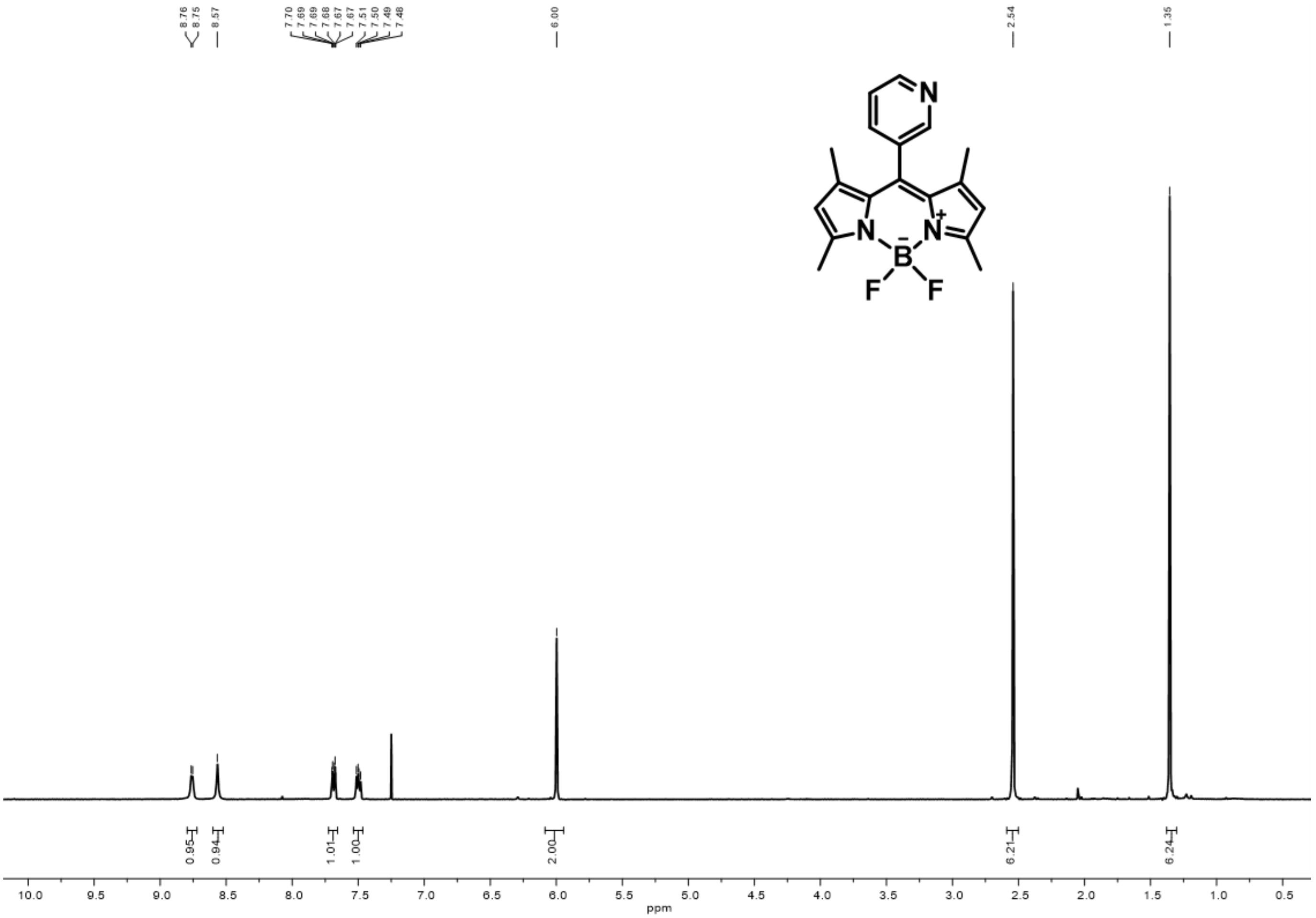


**Figure S1.** $^1H$ NMR spectrum (400 MHz, $CDCl_3$, 298 K) of **1a**.

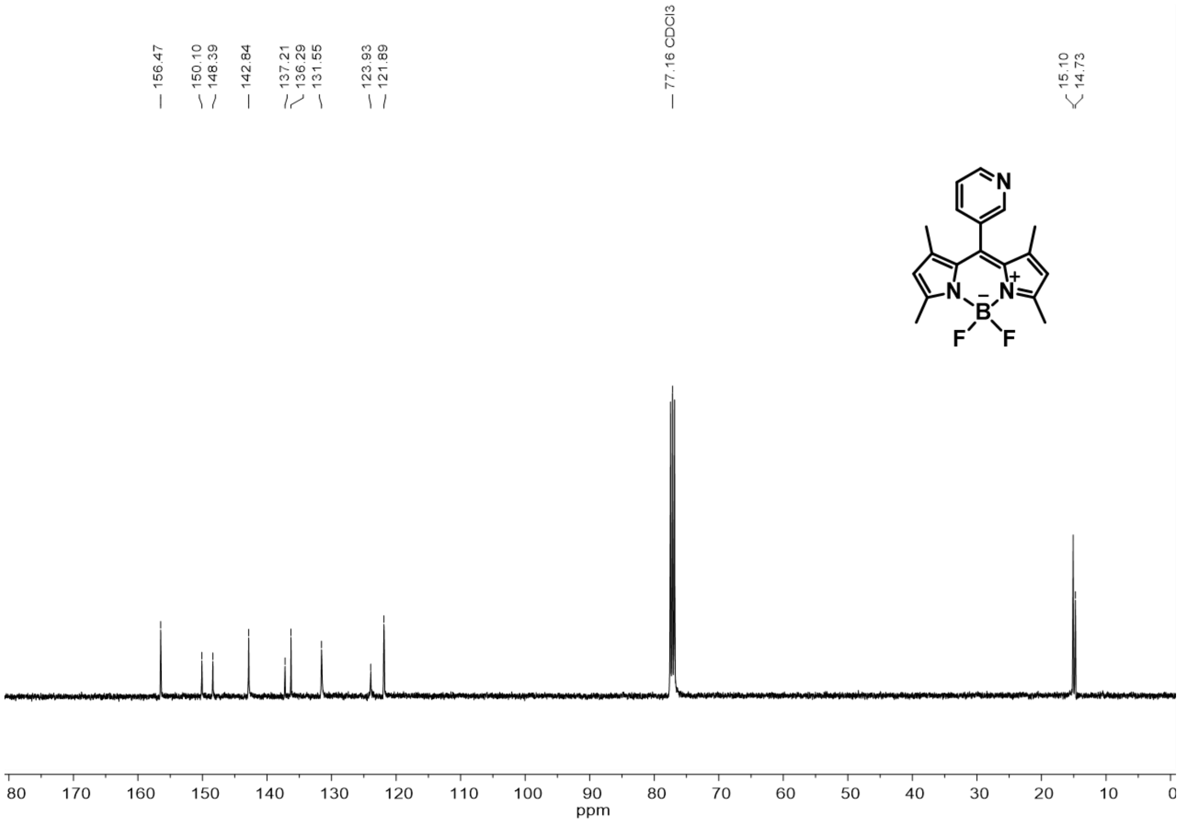


**Figure S2**. $^{13}C\{^{1}H\}$ NMR (101 MHz, $CDCl_3$, 298 K) of **1a**.

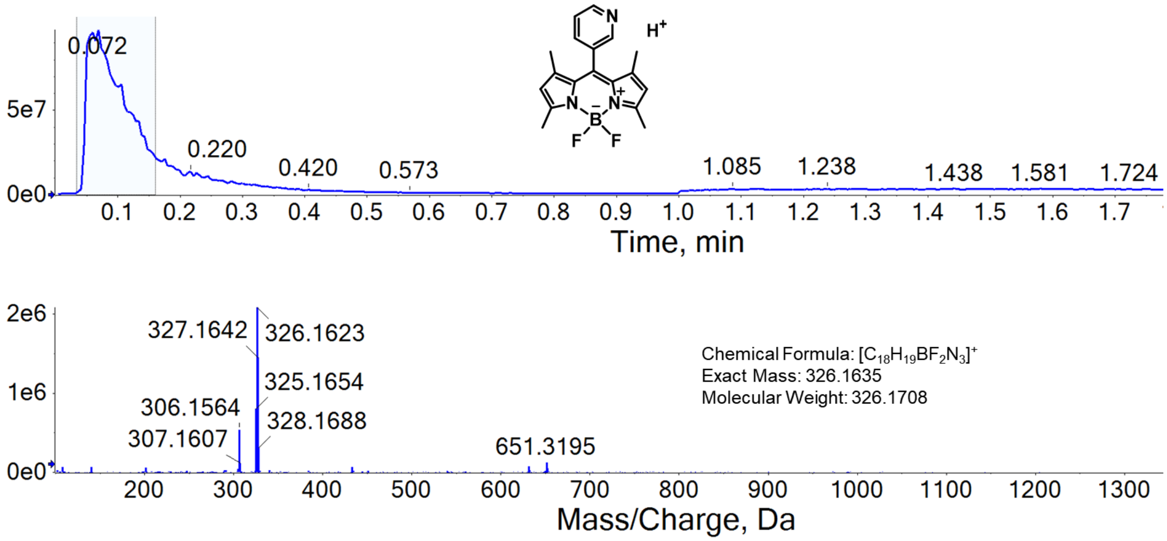


**Figure S3.** HRMS (+ve ESI) spectrum of **1a**

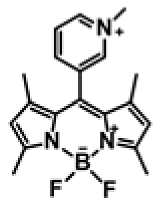

Preparation of **1b**:

**1a** (91.3 mg, 0.28 mmol) was dissolved in 50 mL of MeCN, then methyl iodide (175 μL, 2.81 mmol) was added to the mixture. The mixture was refluxed overnight. The mixture was concentrated by rotary evaporator and the product was purified by neutral aluminium oxide chromatography using $CH_2Cl_2$/MeOH (*v/v* = 10:1) to yield a purple-red solid. Yield: 120 mg, 92 %. $^1$H NMR (400 MHz, $CD_3CN$, 298 K) δ 8.86 (d, J = 5.7 Hz, 2H), 8.62 (d, J = 8.1 Hz, 1H), 8.20 (d, J = 7.5 Hz, 1H), 6.19 (s, 2H), 4.39 (s, 3H), 2.51 (s, 6H), 1.43 (s, 6H). $^{13}$C NMR (101 MHz, $CD_3CN$, 298 K) δ 159.51, 148.32, 147.88, 145.91, 144.88, 136.47, 133.06, 132.55, 130.46, 124.20, 50.54, 16.54, 15.54. HRMS (+ve ESI): calculated for $C_{19}H_{21}BF_2N_3$ $[M]^+$ *m/z* 340.1791, found 340.1842.

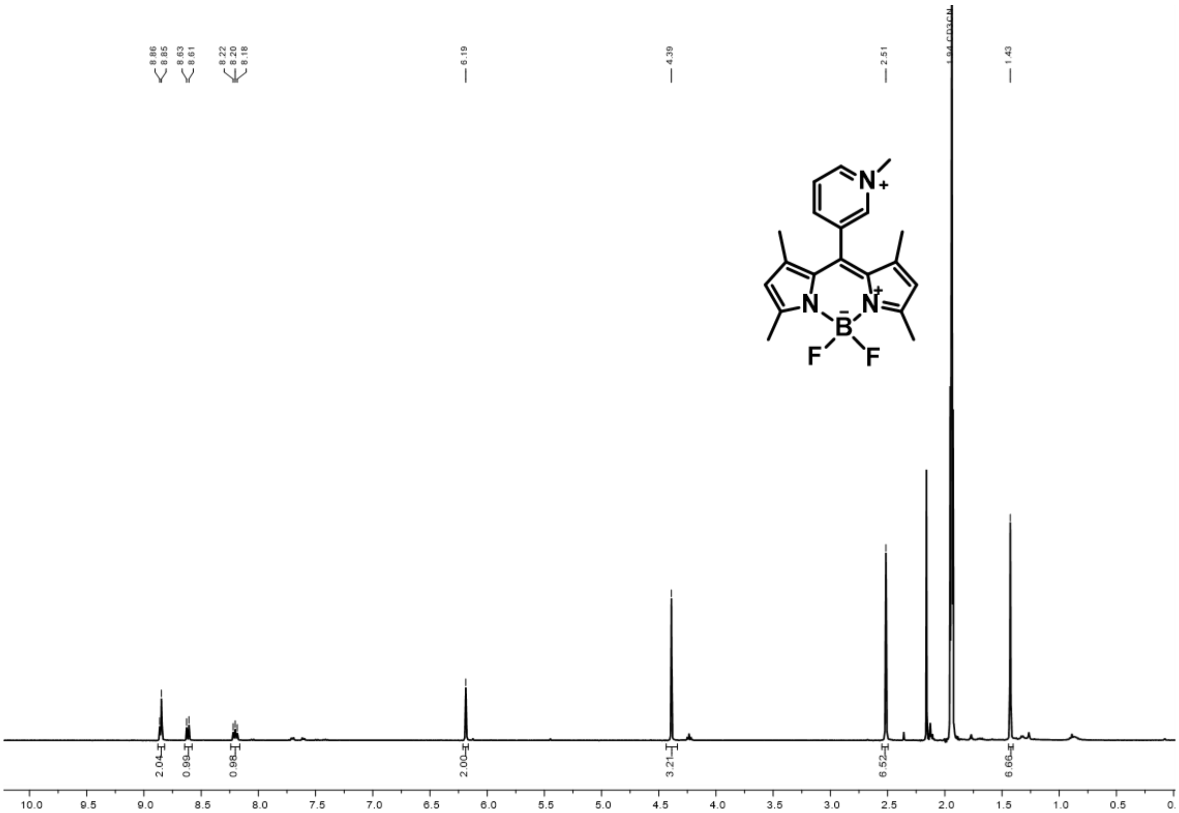


**Figure S4.** $^1$H NMR spectrum (400 MHz, $CD_3CN$, 298 K) of **1b**.

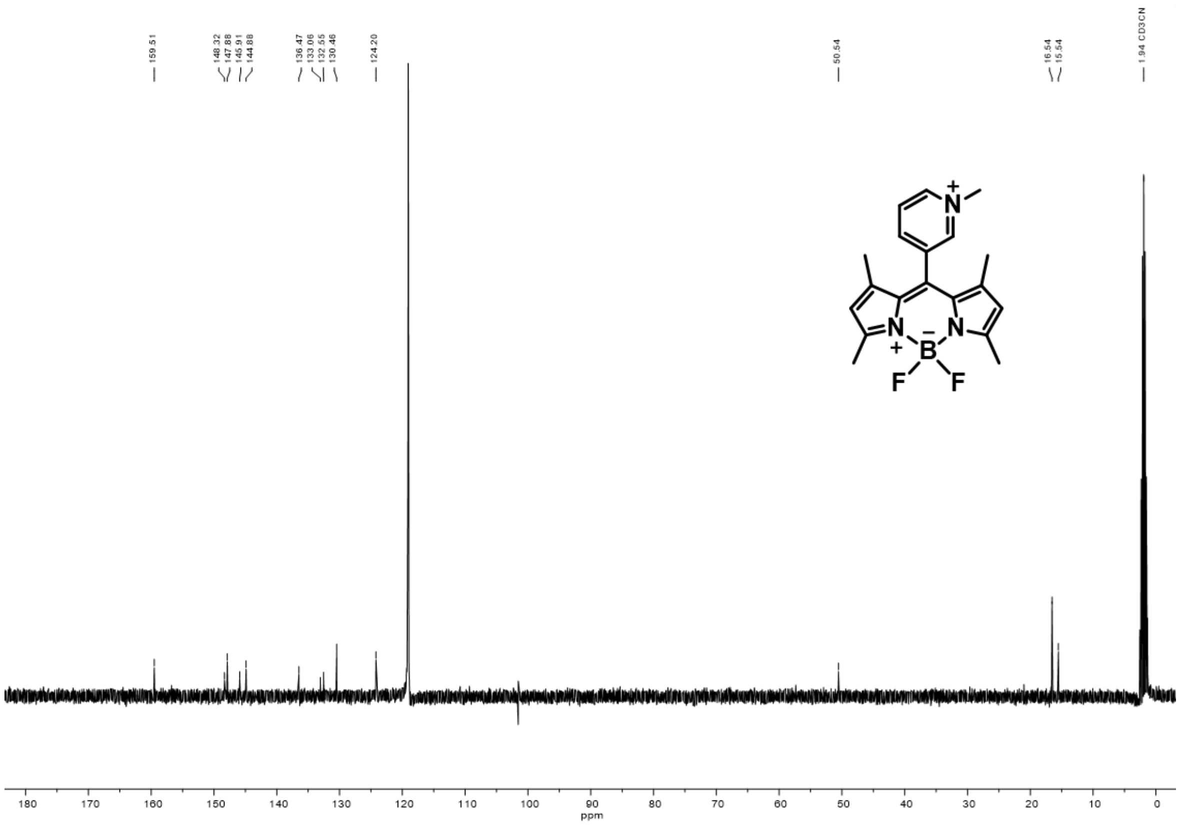


**Figure S5**. $^{13}C\{^{1}H\}$ NMR (101 MHz, $CD_3CN$, 298 K) of **1b**.

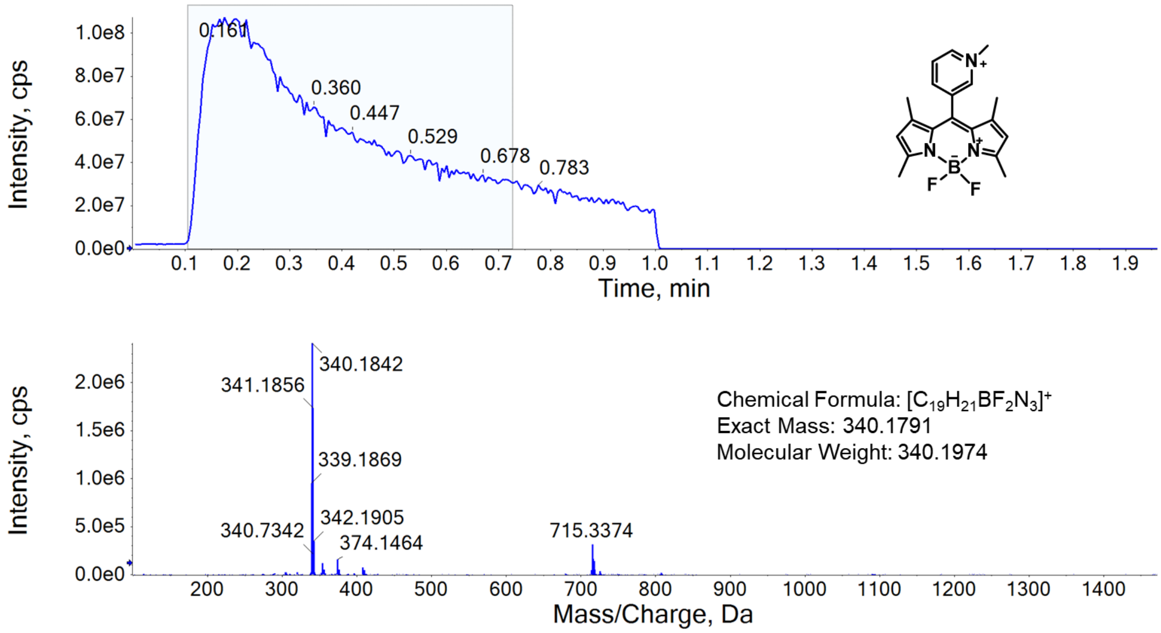


**Figure S6.** HRMS (+ve ESI) spectrum of **1b**.

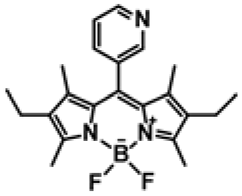

Preparation of **1c**:

The compound was prepared using the similar procedure as described in the preparing of compound **1a** using 3-pyridine carboxaldehyde (0.42 g, 3.9 mmol) and 3-ethyl-2,4-dimethylpyrrole (0.96 g, 7.84 mmol) instead of 2,4-dimethylpyrrole. Yield: 140 mg, 10 %, red solid. $^{1}$H NMR (400 MHz, $CDCl_3$, 298 K) δ 9.01 – 8.96 (m, 1H), 8.70 (s, 1H), 8.31 (d, J = 6.7 Hz, 1H), 8.09 – 7.99 (m, 1H), 2.55 (s, 6H), 2.31 (q, J = 7.5 Hz, 4H), 1.24 (s, 6H), 0.98 (t, J = 7.5 Hz, 6H). $^{13}$C NMR (101 MHz, $CDCl_3$, 298 K) δ 154.87, 149.70, 148.51, 138.02, 136.79, 135.50, 133.51, 132.39, 130.95, 123.97, 17.18, 14.72, 12.71, 12.43. HRMS (+ve ESI): calculated for $C_{22}H_{27}BF_2N_3$ $[M+H]^+$ *m/z* 382.2261, found 382.2210.

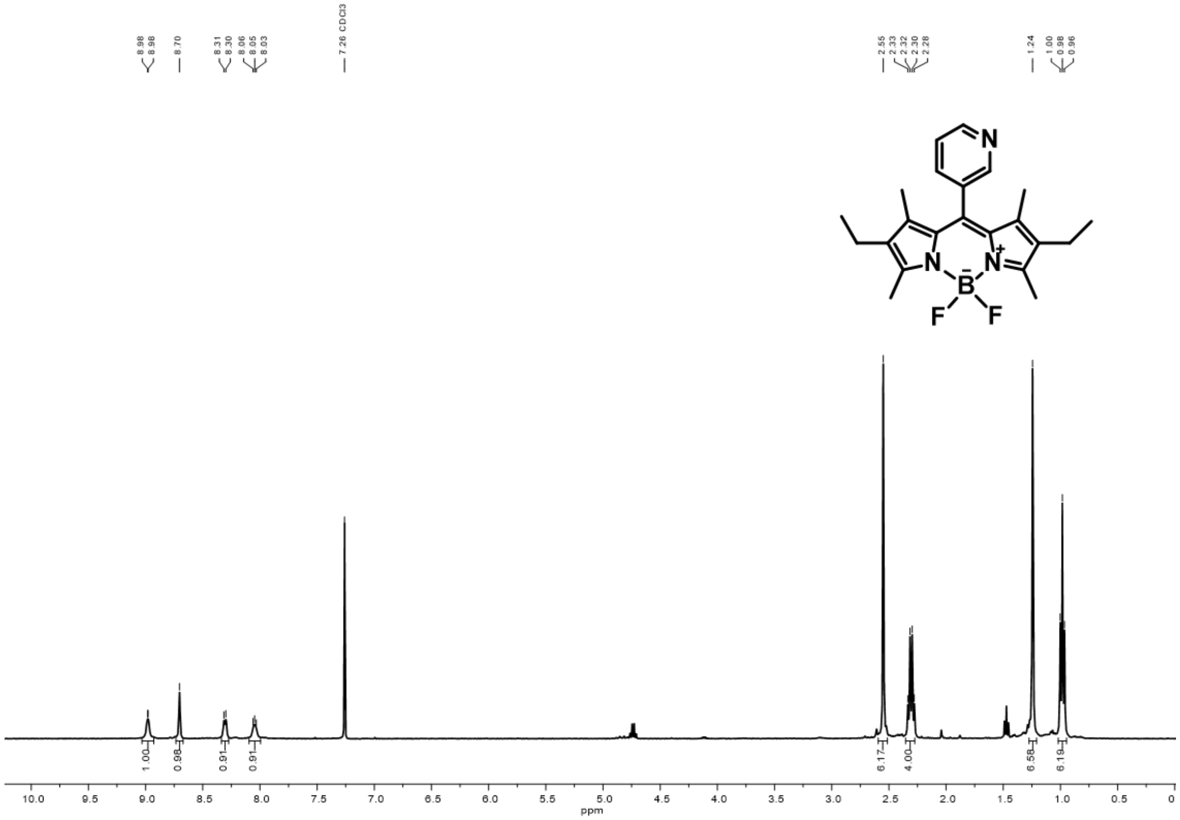


**Figure S7.** $^{1}$H NMR spectrum (400 MHz, $CDCl_3$, 298 K) of **1c**.

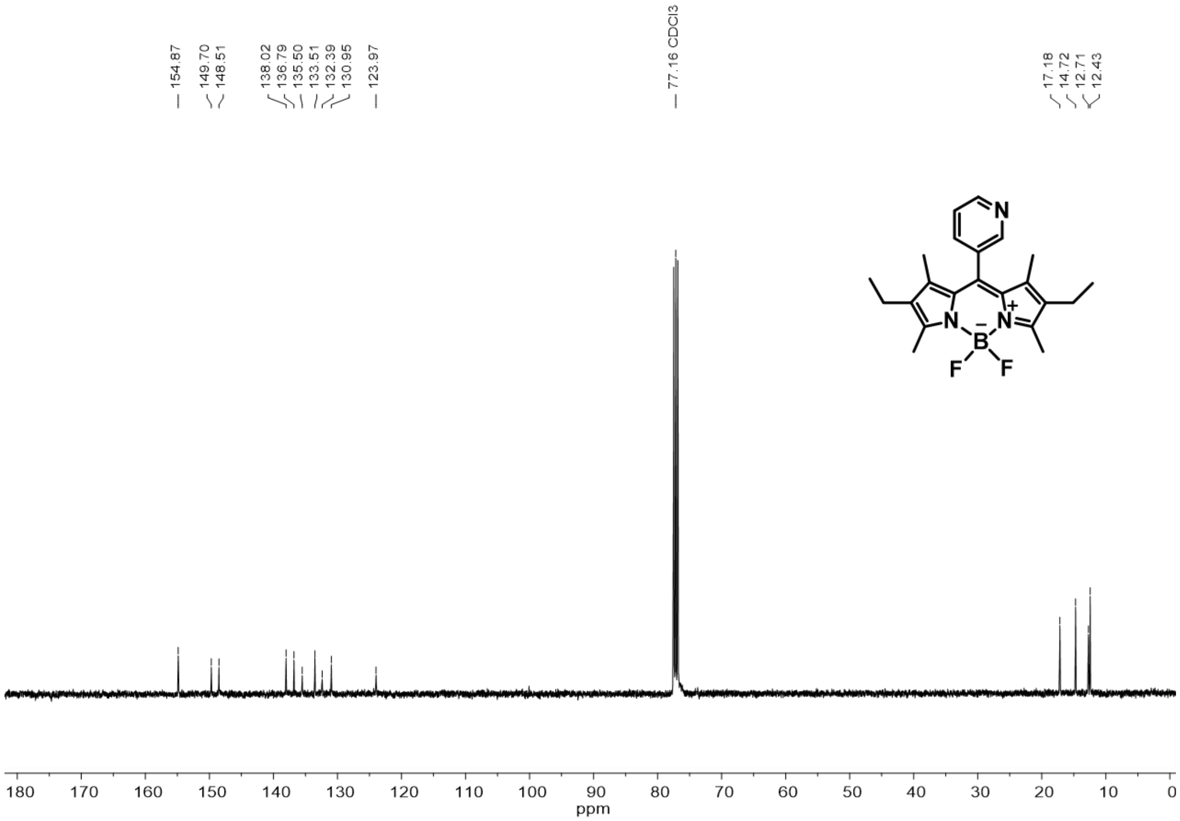


**Figure S8**. $^{13}C\{^{1}H\}$ NMR (101 MHz, $CDCl_3$, 298 K) of **1c**.

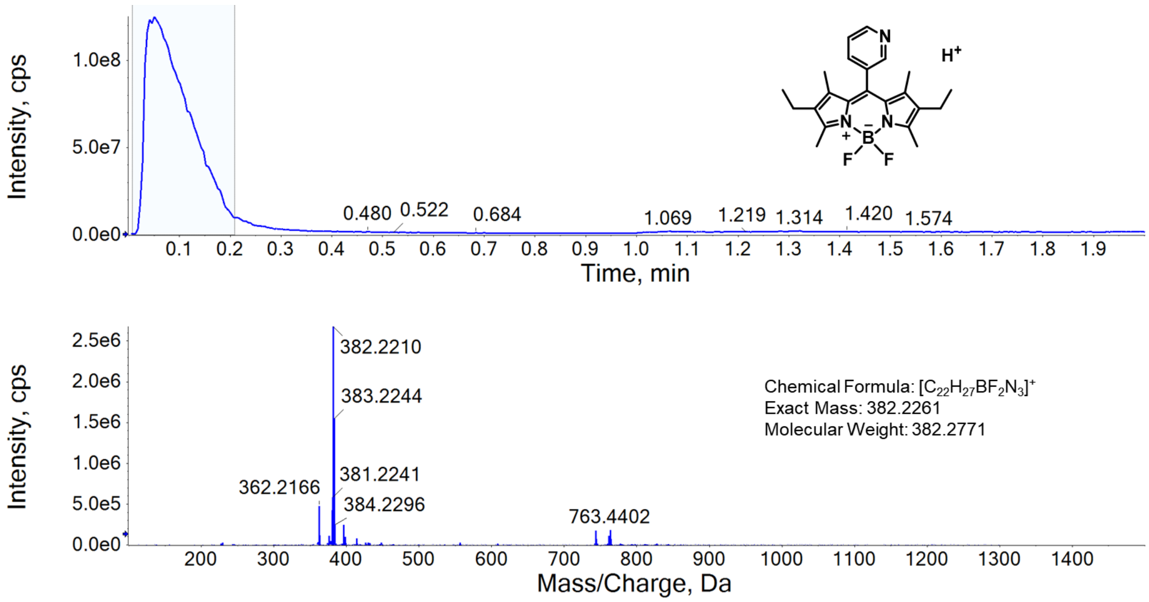


**Figure S9.** HRMS (+ve ESI) spectrum of **1c**.

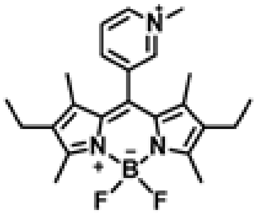


Preparation of **1d**:

The compound was prepared using the similar procedure as described in the preparing of compound **1b**, using **1c** (100 mg, 0.262 mmol) and methyl iodide (195 μL, 3.13 mmol) as reactants. Yield: 123 mg, 89.7%, red solid $^{1}$H NMR (400 MHz, $CD_3CN$, 298 K) δ 8.82 – 8.75 (m, 2H), 8.59 (d, J = 8.1 Hz, 1H), 8.22 – 8.14 (m, 1H), 4.36 (s, 3H), 2.50 (s, 6H), 2.36 (q, J = 7.6 Hz, 4H), 1.32 (s, 6H), 0.99 (t, J = 7.6 Hz, 6H). $^{13}$C NMR (101 MHz, $CD_3CN$, 298 K) δ 157.84, 148.12, 148.02, 146.04, 139.91, 137.24, 136.04, 131.93, 131.58, 130.33, 50.48, 18.09, 15.49, 14.04, 13.63. HRMS (+ve ESI): calculated for $C_{23}H_{29}BF_2N_3^+$ $[M]^+$ *m/z* 396.2417, found 396.2446.

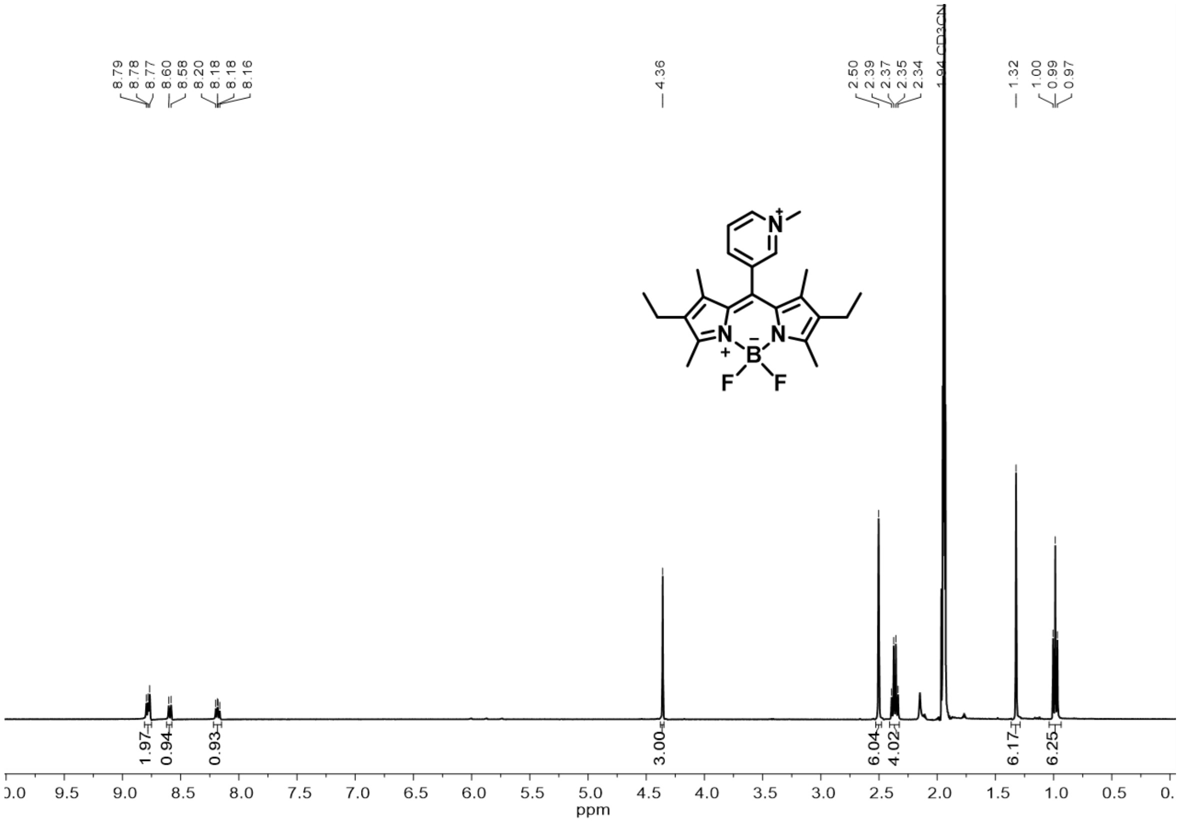


**Figure S10.** $^{1}$H NMR spectrum (400 MHz, $CD_3CN$, 298 K) of **1d**.

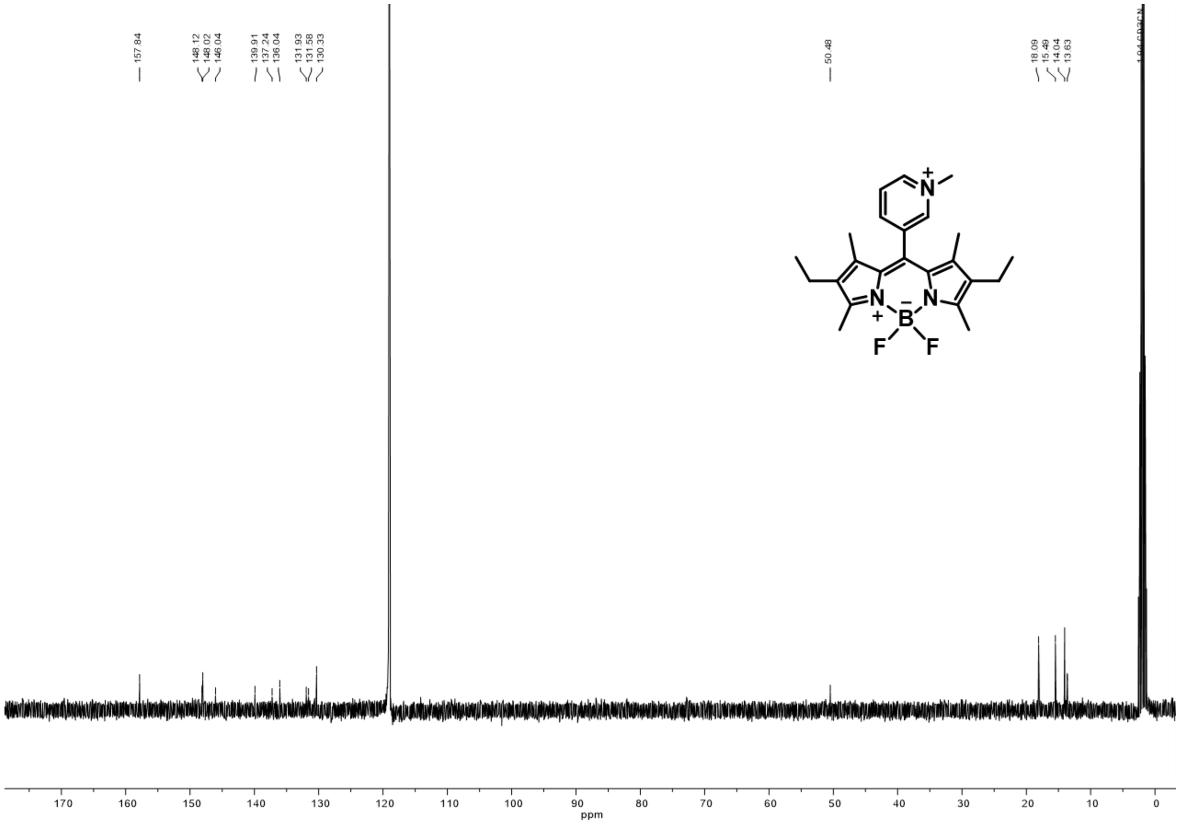

**Figure S11**. $^{13}C\{^{1}H\}$ NMR (101 MHz, $CD_3CN$, 298K) of **1d**.

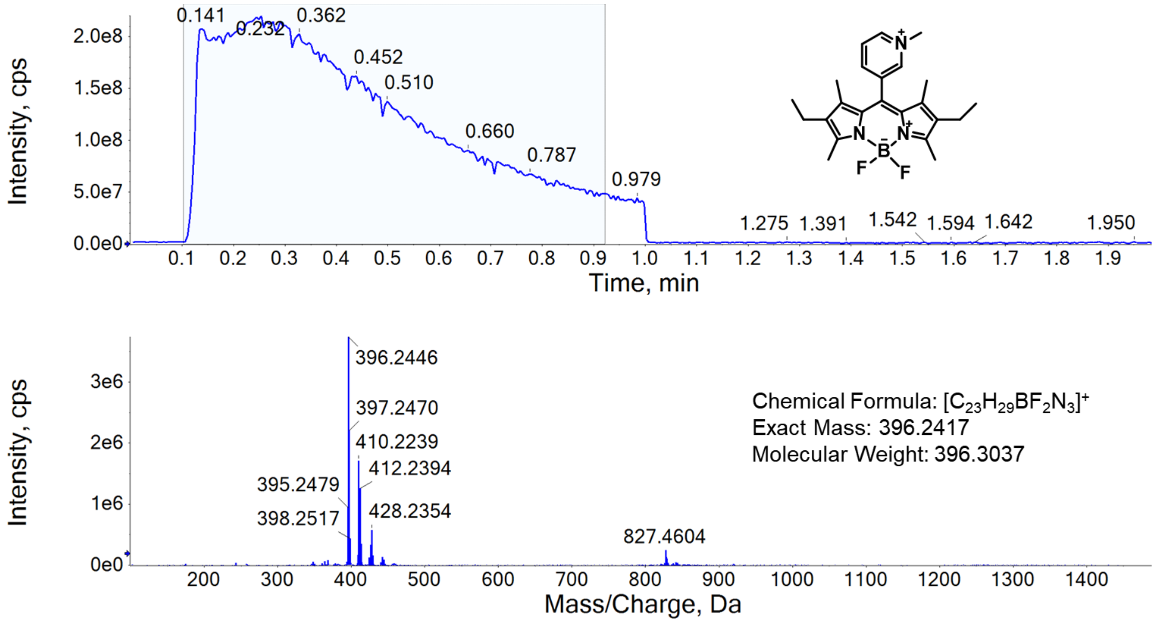


**Figure S12.** HRMS (+ve ESI) spectrum of **1d**.

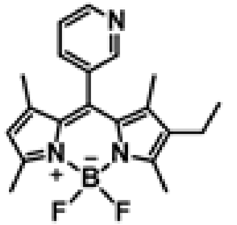

Preparation of **1e**: The compound was prepared using the similar procedure as described in the preparing of compound **1a** using 3-pyridine carboxaldehyde (0.42 g, 3.9 mmol), 3-ethyl-2,4-dimethylpyrrole (0.48 g, 3.90 mmol), and 2,4-dimethylpyrrole (0.37 g, 3.90 mmol). Yield: 114 mg, 8.2 %, orange solid. $^{1}H$ NMR (400 MHz, $CDCl_3$, 298 K) δ 8.93 (d, J = 4.8 Hz, 1H), 8.69 (s, 1H), 8.17 (d, J = 7.9 Hz, 1H), 7.92 (dd, J = 7.6, 5.6 Hz, 1H), 6.01 (s, 1H), 2.57 (s, 3H), 2.55 (s, 3H), 2.32 (q, J = 7.6 Hz, 2H), 1.33 (s, 3H), 1.28 (s, 3H), 0.99 (t, J = 7.6 Hz, 3H). $^{13}C$ NMR (101 MHz, $CDCl_3$, 298 K) δ 156.64, 154.83, 149.78, 148.31, 141.58, 138.90, 136.64, 136.18, 134.26, 131.94, 131.44, 131.00, 123.98, 121.21, 17.14, 15.04, 14.61, 12.80, 12.45. HRMS (+ve ESI): calculated for $C_{20}H_{23}BF_2N_3$ $[M+H]^+$ *m/z* 354.1948, found 354.1934.

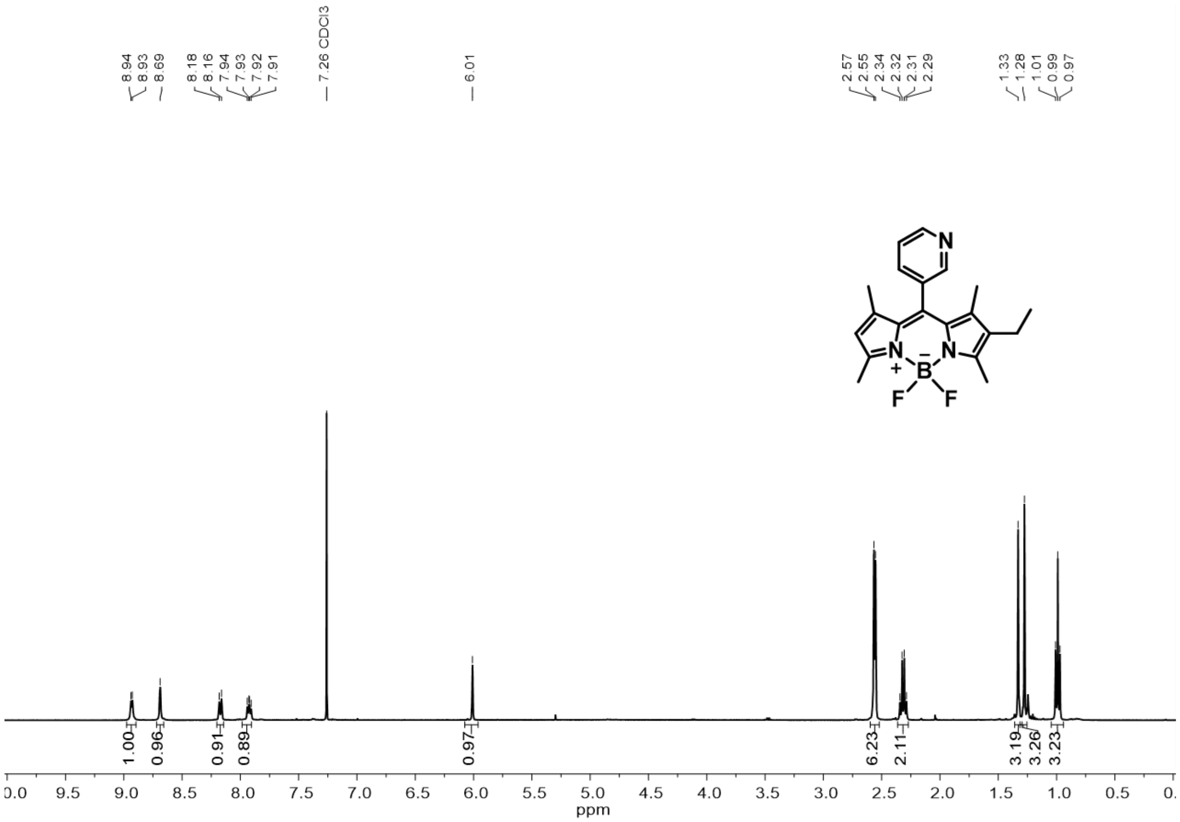


**Figure S13**. $^{1}H$ NMR spectrum (400 MHz, $CDCl_3$, 298 K) of **1e**.

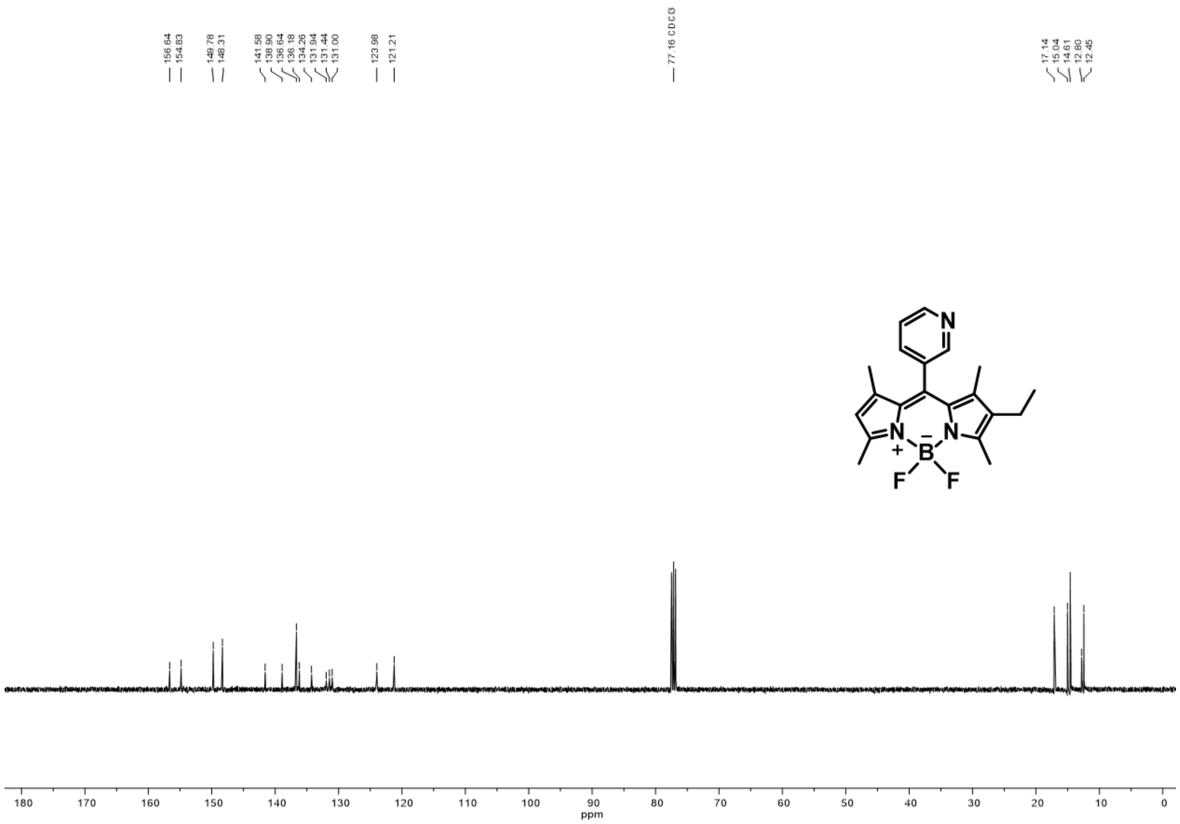


**Figure S14**. $^{13}C\{^{1}H\}$ NMR (101 MHz, $CDCl_3$, 298 K) of **1e**.

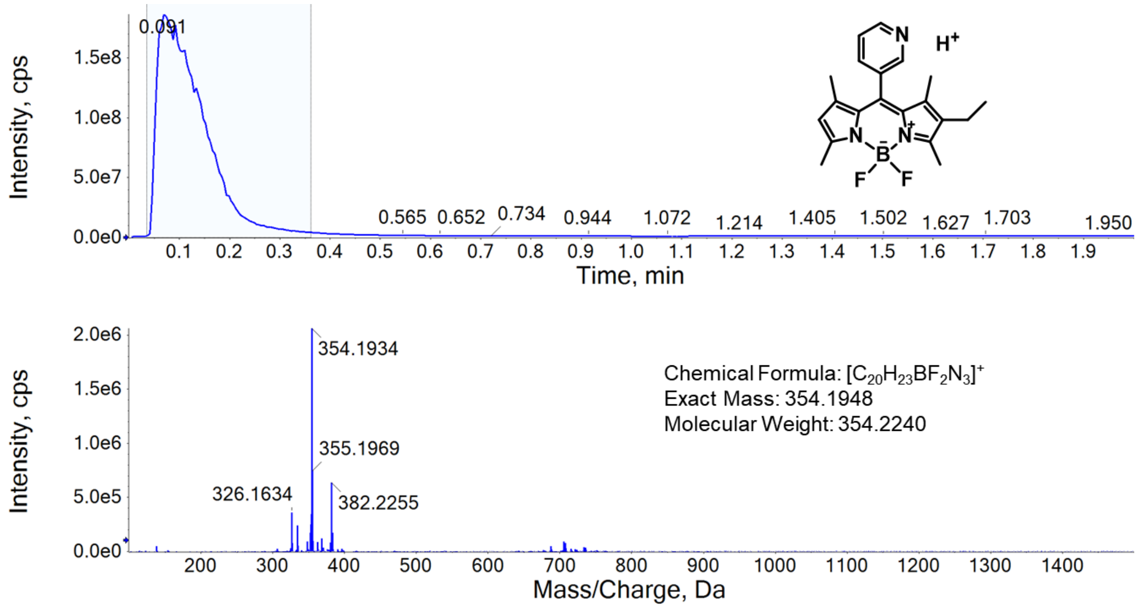


**Figure S15.** HRMS (+ve ESI) spectrum of **1e**.

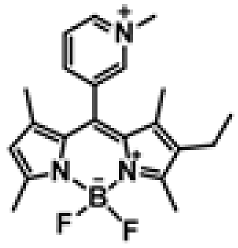

Preparation of **1f**:

The compound was prepared using the similar procedure as described in the preparing of compound **1b**, using **1f** (100 mg, 0.28 mmol) and methyl iodide (205 µL, 3.3 mmol). Yield: 18.5 mg, 13.2%, red solid. $^1$H NMR (400 MHz, $CD_3CN$, 298 K) δ 9.05 (d, J = 6.1 Hz, 1H), 9.00 (s, 1H), 8.61 – 8.57 (m, 1H), 8.23 – 8.17 (m, 1H), 5.45 (s, 1H), 4.45 (s, 3H), 2.60 – 2.46 (m, 6H), 2.37 (q, J = 7.5 Hz, 2H), 1.46 – 1.31 (m, 6H), 0.99 (t, J = 7.6 Hz, 3H). $^{13}$C NMR (101 MHz, $CD_3CN$, 298 K) δ 159.43, 157.04, 147.61, 147.43, 145.44, 142.92, 140.30, 136.38, 132.06, 131.64, 131.35, 129.84, 126.45, 122.84, 49.95, 30.70, 17.56, 15.92, 14.81, 13.55, 13.26. HRMS (+ve ESI): calculated for $C_{21}H_{25}BF_2N_3$ $[M]^+$ $m/z$ 368.2104, found 368.2097.

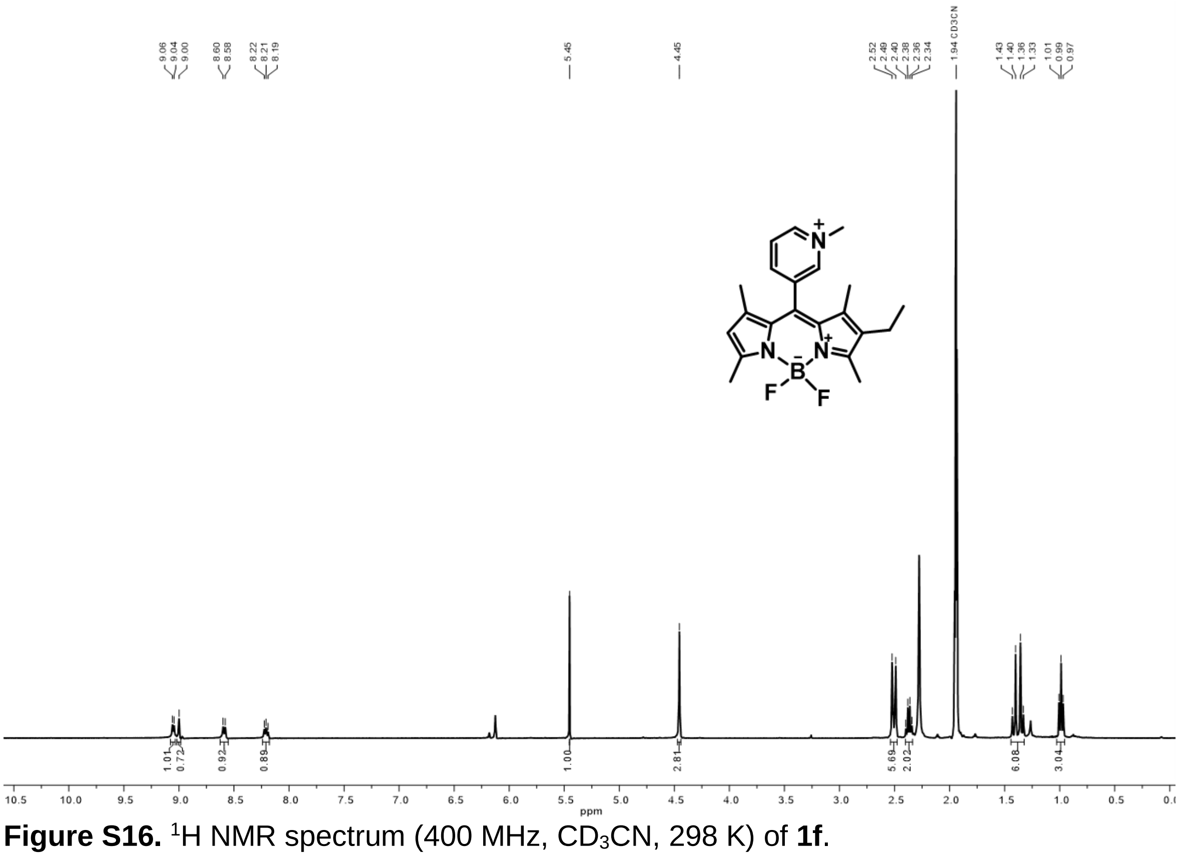


**Figure S16.** $^1$H NMR spectrum (400 MHz, $CD_3CN$, 298 K) of **1f**.

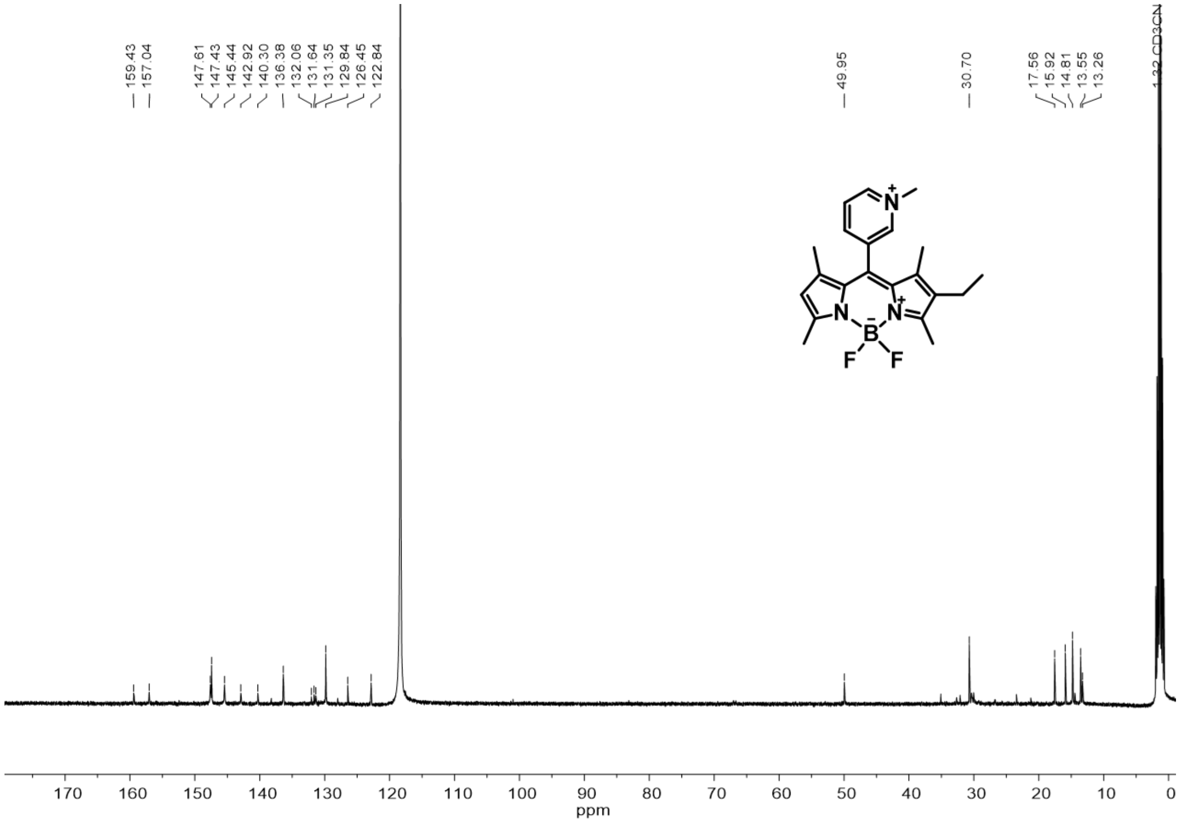


**Figure S17**. $^{13}C\{^{1}H\}$ NMR (101 MHz, $CD_3CN$, 298 K) of **1f**.

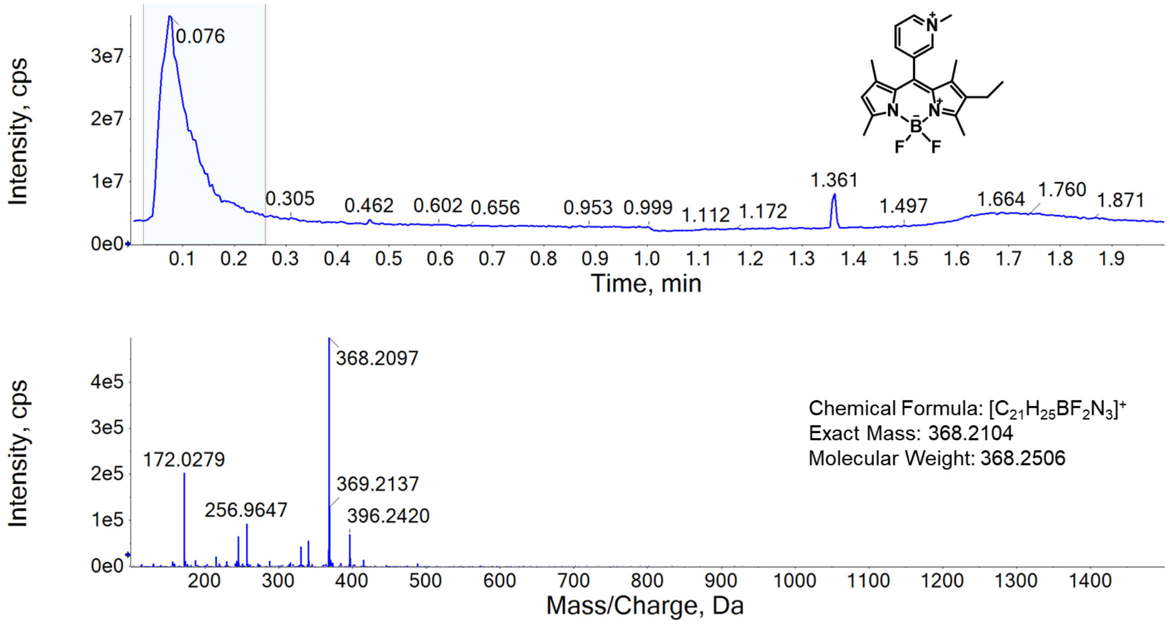


**Figure S18.** HRMS (+ve ESI) spectrum of **1f**.

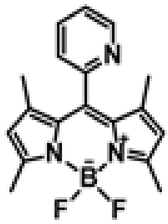

Preparation of **2a**:

The compound was prepared using the similar procedure as described in the preparing of compound **1a** using 2-pyridine carboxaldehyde (0.42 g, 3.9 mmol) instead of 3-pyridine carboxaldehyde. Yield = 480 mg, 38 %, orange solid. $^{1}H$ NMR (400 MHz, $CDCl_3$, 298 K) δ 8.82 (d, J = 3.5 Hz, 1H), 7.92 (t, J = 7.7 Hz, 1H), 7.57 – 7.44 (m, 2H), 5.99 (s, 2H), 2.55 (s, 6H), 1.32 (s, 6H). $^{13}C$ NMR (101 MHz, $CDCl_3$, 298 K) δ 156.43, 153.95, 150.24, 142.67, 138.55, 137.19, 131.48, 124.51, 124.03, 121.37, 14.79, 13.87. HRMS (+ve ESI): calculated for $C_{18}H_{19}BF_2N_3$ $[M+H]^+$ *m/z* 326.1635., found 326.1593.

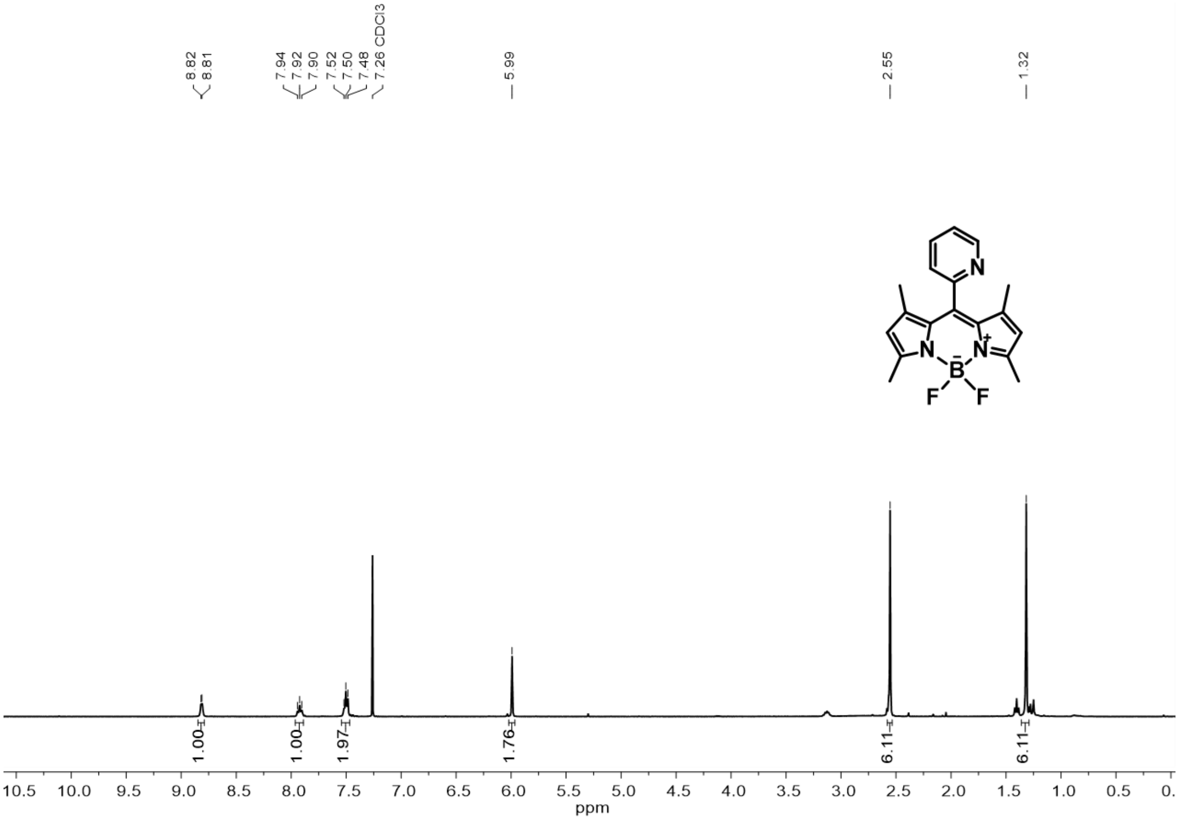


**Figure S19.** $^{1}H$ NMR spectrum (400 MHz, $CDCl_3$, 298 K) of **2a**.

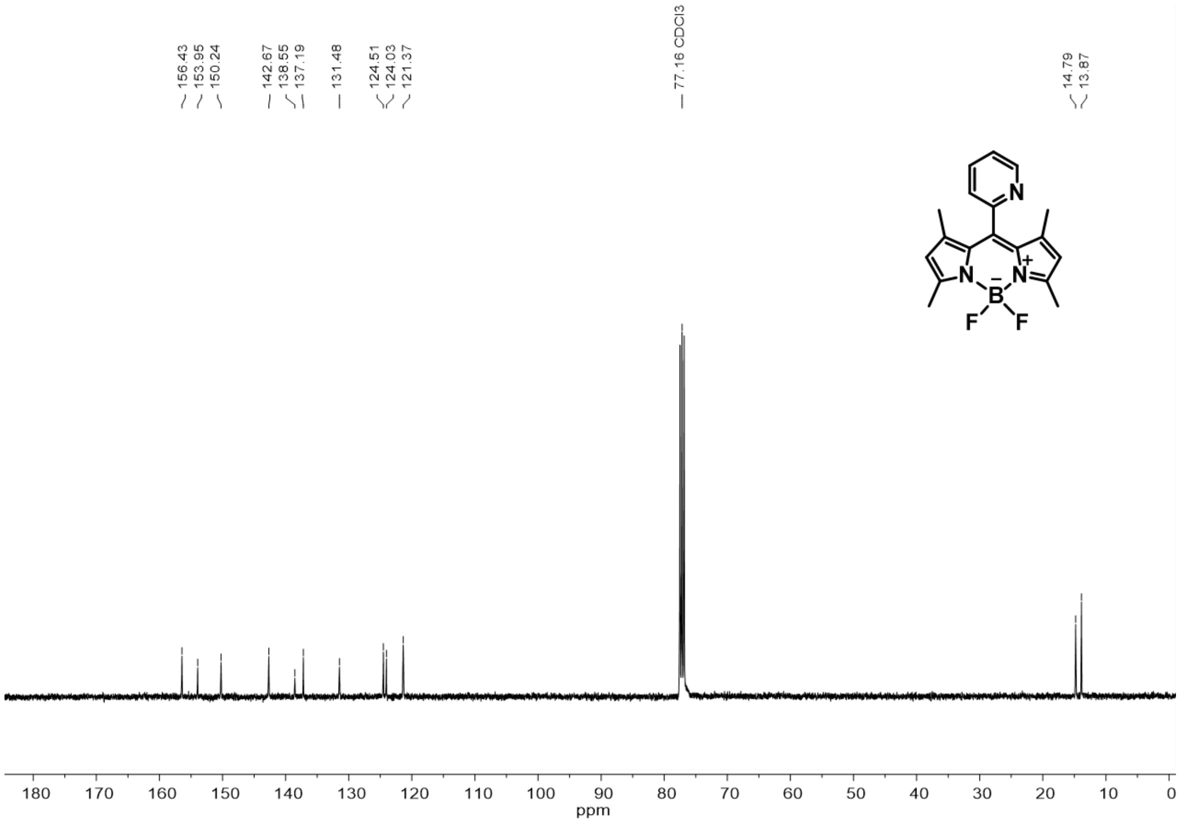


**Figure S20**. $^{13}C\{^{1}H\}$ NMR (101 MHz, $CDCl_3$, 298 K) of **2a**.

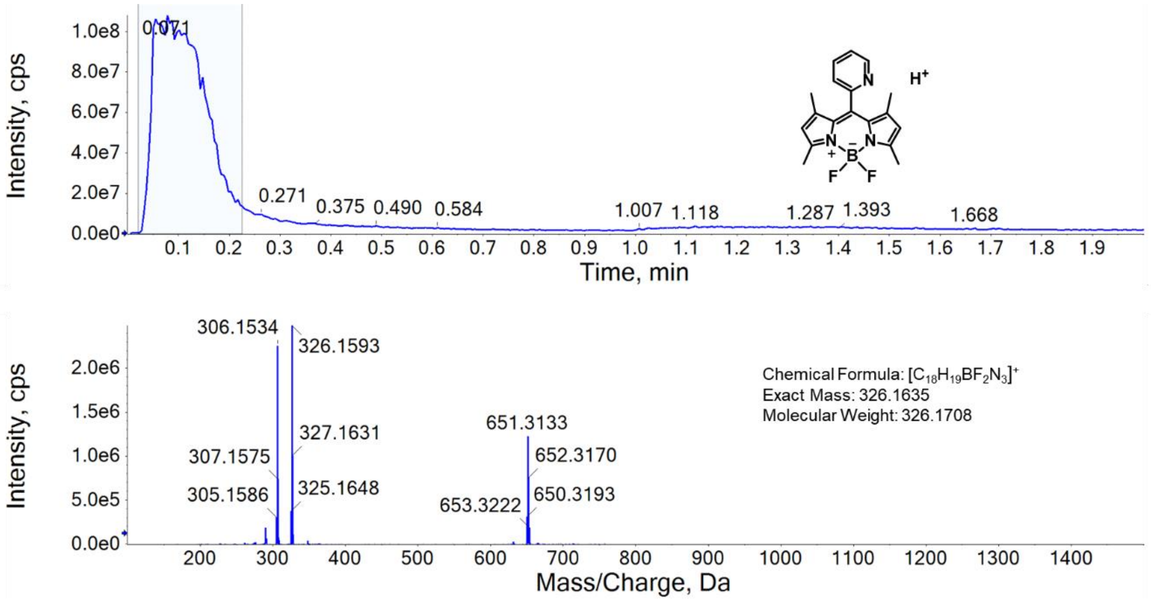


**Figure S21.** HRMS (+ve ESI) spectrum of **2a**.

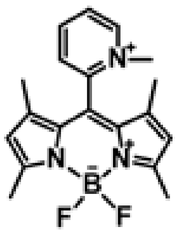

Preparation of **2b**:

The compound was prepared using the similar procedure as described in the preparing of compound **1a**, using **2a** (201 mg, 0.62 mmol) and methyl iodide (385 µL, 6.18 mmol). Yield = 107 mg, 37 %, brownish red solid. $^{1}$H NMR (400 MHz, $CD_3CN$, 298K) δ 9.16 (d, J = 5.6 Hz, 1H), 8.69 (t, J = 7.8 Hz, 1H), 8.27 (d, J = 7.7 Hz, 2H), 6.24 (s, 2H), 4.25 (s, 3H), 2.54 (s, 6H), 1.34 (s, 6H). $^{13}$C NMR (101 MHz, $CD_3CN$, 298 K) δ 161.57, 149.86, 149.22, 146.68, 144.02, 132.49, 131.11, 124.63, 124.57, 47.89, 15.77, 13.99. HRMS (+ve ESI): calculated for $C_{19}H_{21}BF_2N_3$ [M]$^+$ *m/z* 340.1791, found 340.1839.

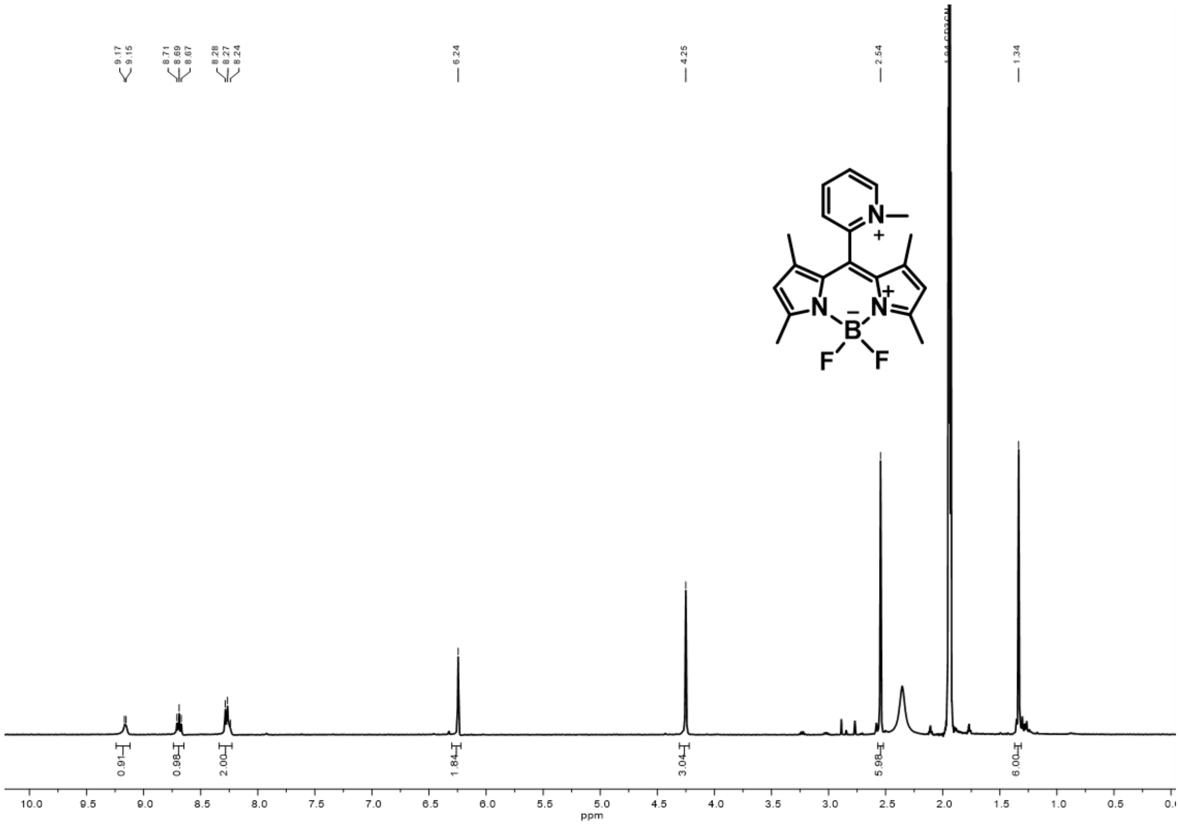


**Figure S22.** $^{1}$H NMR spectrum (400 MHz, $CD_3CN$, 298 K) of **2b**.

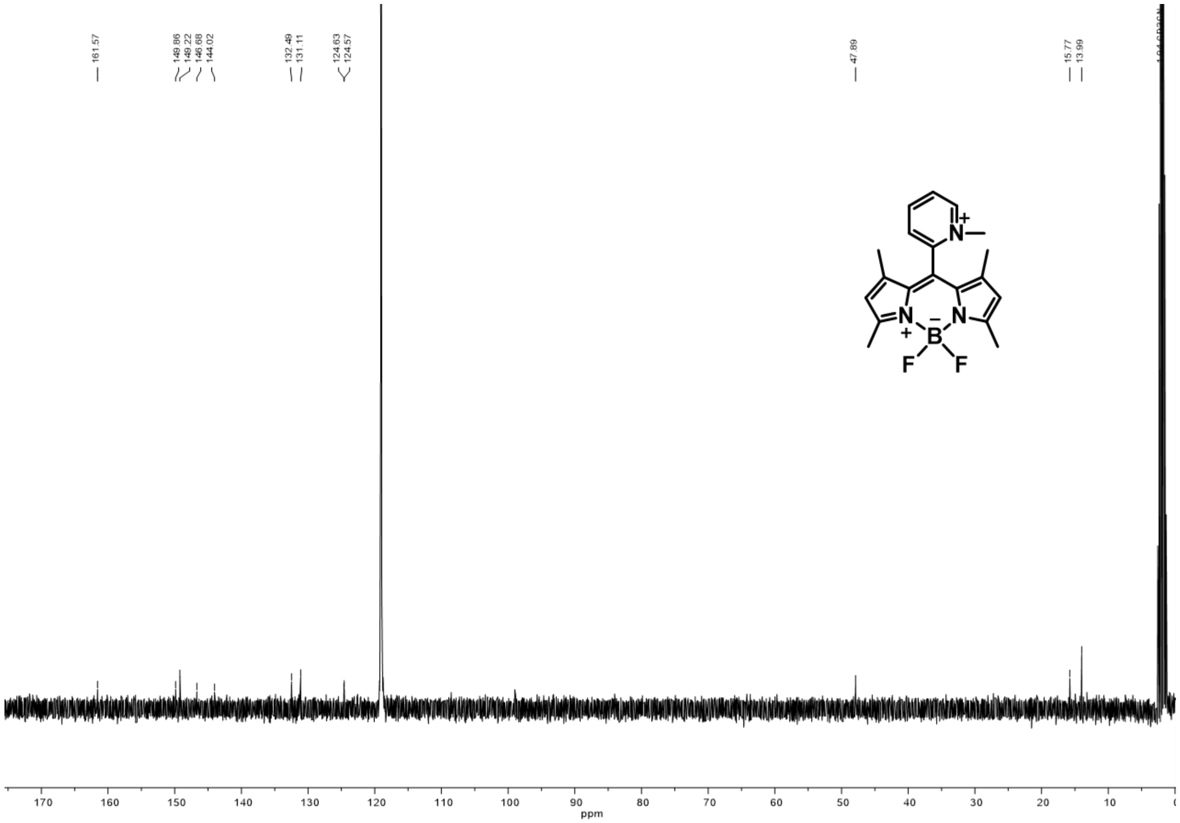


**Figure S23**. $^{13}C\{^{1}H\}$ NMR (101 MHz, $CD_3CN$, 298 K) of **2b**.

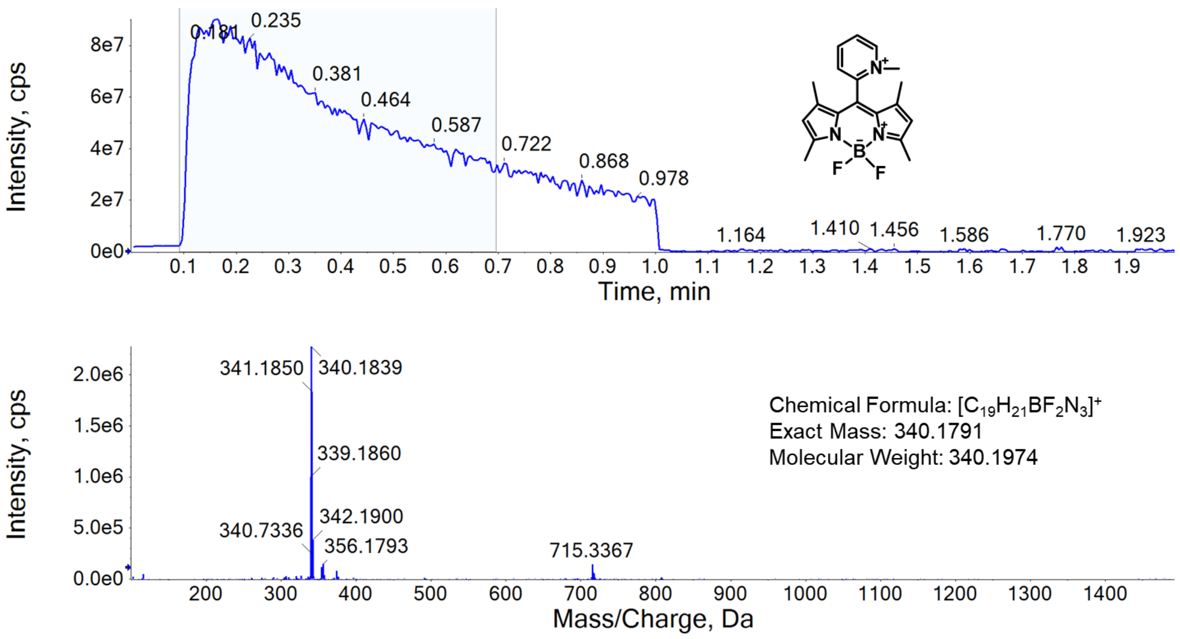


**Figure S24.** HRMS (+ve ESI) spectrum of **2b**.

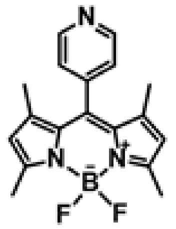

Preparation of **3a**:

The compound was prepared using the similar procedure as described in the preparing of compound **1a**, using 4-pyridine carboxaldehyde (0.42 g, 3.9 mmol) instead of 2-pyridine carboxaldehyde. Yield: 290 mg, 23 %, red solid. $^1$H NMR (400 MHz, $CDCl_3$, 298 K) δ 9.01 (d, J = 6.4 Hz, 2H), 7.97 (d, J = 6.4 Hz, 2H), 6.08 (s, 2H), 2.58 (s, 6H), 1.37 (s, 6H). HRMS (+ve ESI): calculated for $C_{18}H_{19}BF_2N_3$ $[M+H]^+$ *m/z* 326.1635., found 326.1615.

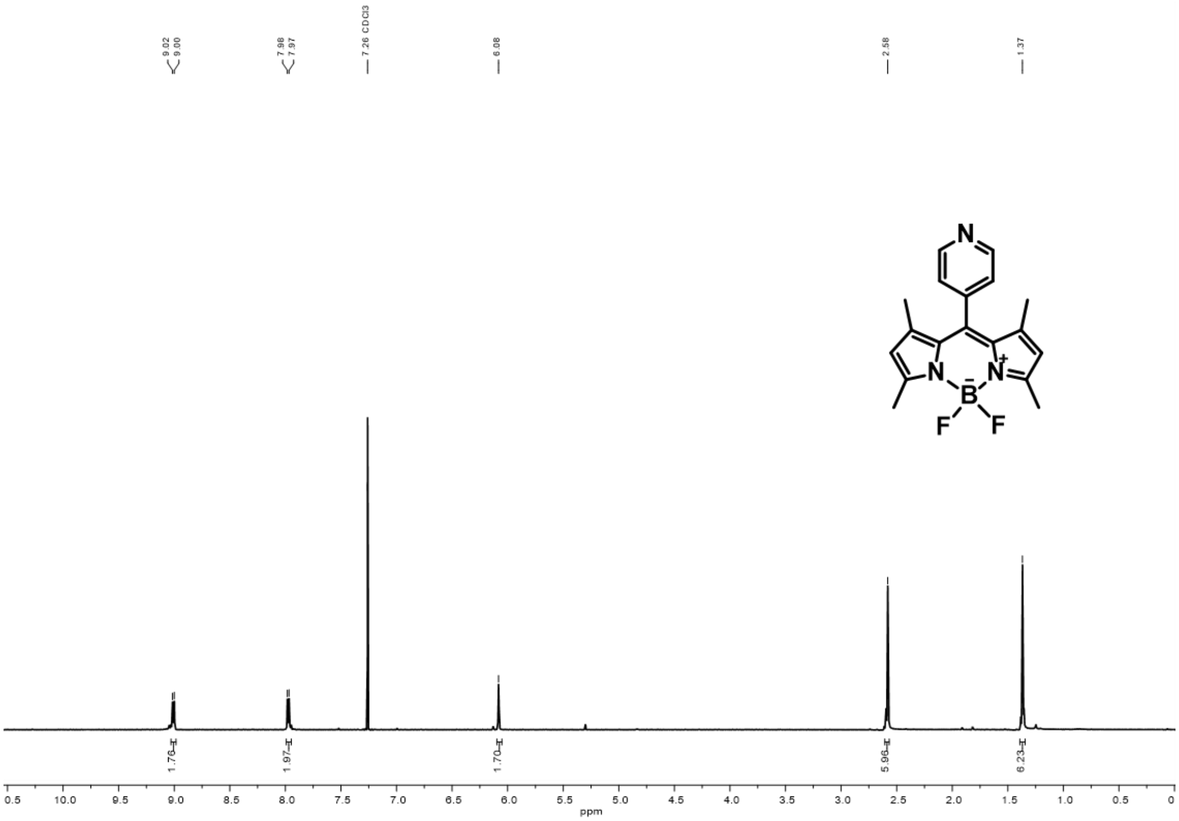

**Figure S25.** $^1$H NMR spectrum (400 MHz, $CDCl_3$, 298 K) of **3a**.

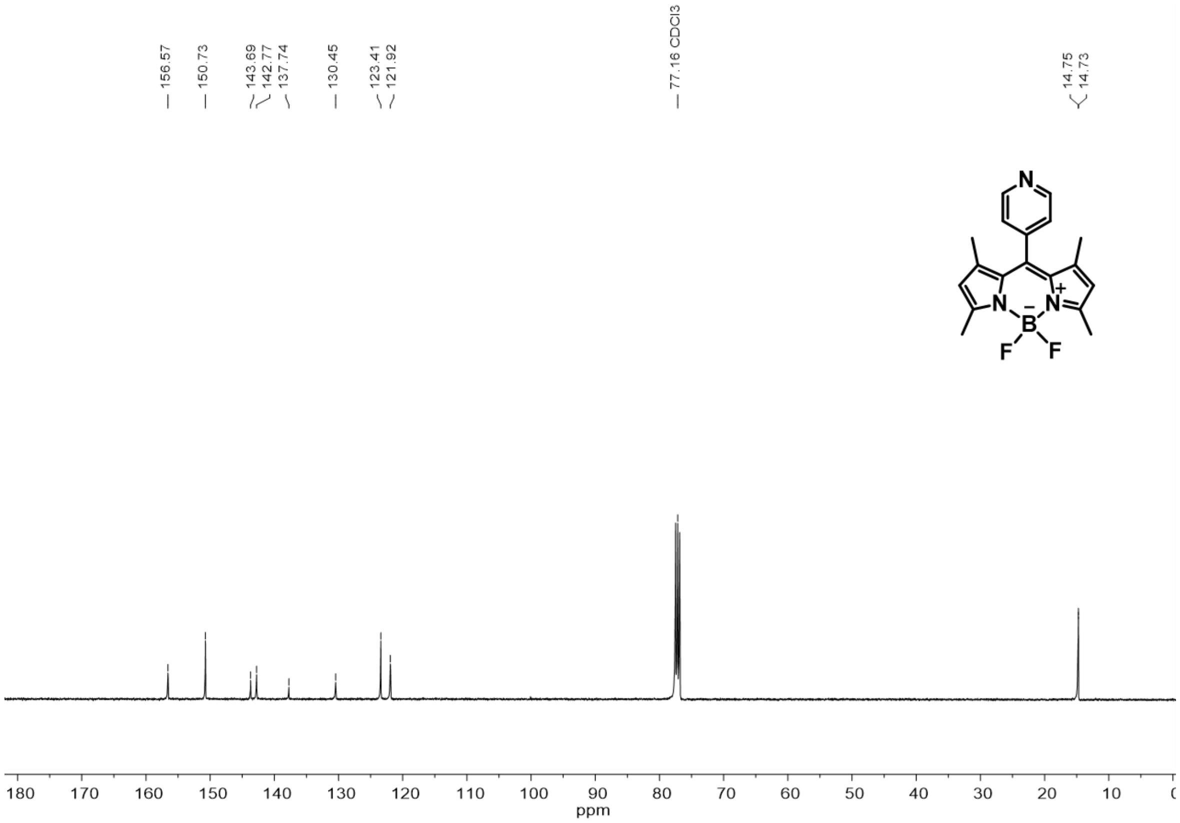


**Figure S26**. $^{13}C\{^{1}H\}$ NMR (101 MHz, $CDCl_3$, 298 K) of **3a**.

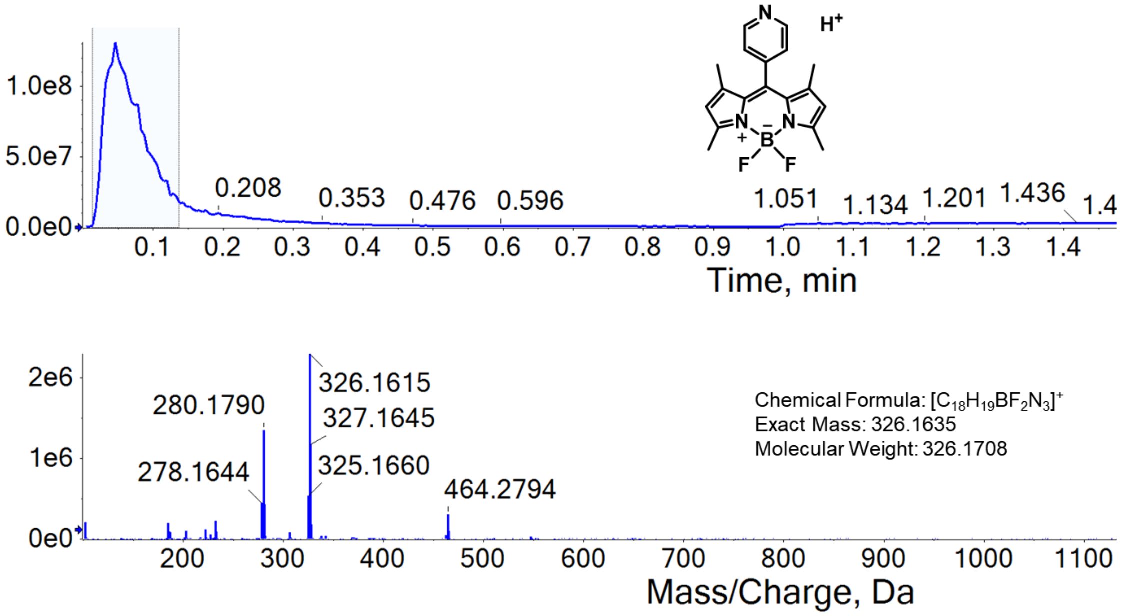


**Figure S27.** HRMS (+ve ESI) spectrum of **3a**.

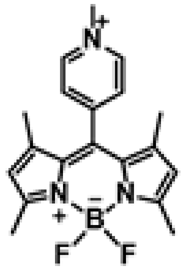

Preparation of **3b**: The compound was prepared using the similar procedure as described in the preparing of compound **1b**, using **3a** (100 mg, 0.31 mmol) and methyl iodide (195 μL, 3.13 mmol). Yield: 74 mg, 51 %, red solid. $^{1}$H NMR (400 MHz, $CD_3CN$, 298 K) δ 8.76 (d, J = 6.5 Hz, 2H), 8.14 (d, J = 6.5 Hz, 2H), 6.17 (s, 2H), 4.37 (s, 3H), 2.51 (s, 6H), 1.43 (s, 6H). $^{13}$C NMR (101 MHz, $CD_3CN$, 298 K) δ 157.74, 152.15, 151.40, 146.60, 143.14, 129.57, 128.59, 122.50, 43.88, 14.67, 13.94. HRMS (+ve ESI): calculated for $C_{19}H_{21}BF_2N_3$ $[M]^+$ *m/z* 340.1791, found 340.1832.

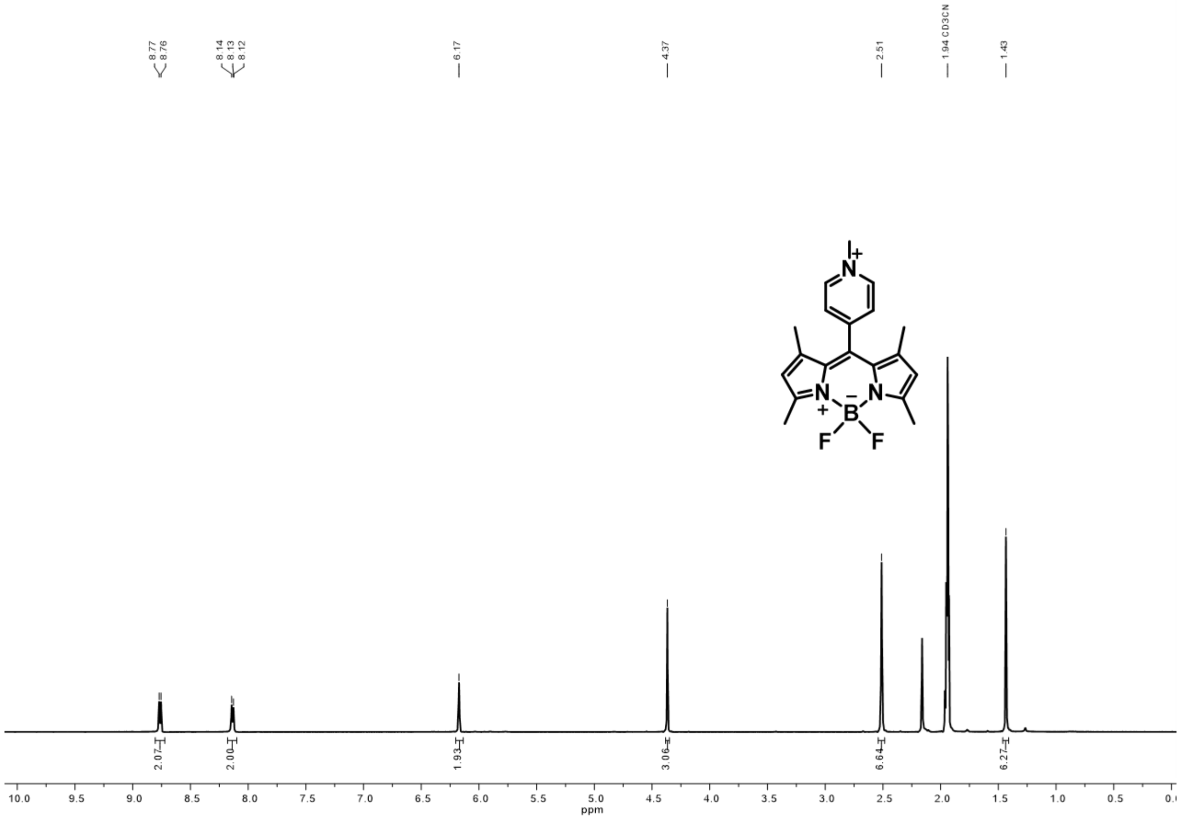


**Figure S28.** $^{1}$H NMR spectrum (400 MHz, $CD_3CN$, 298 K) of **3b**.

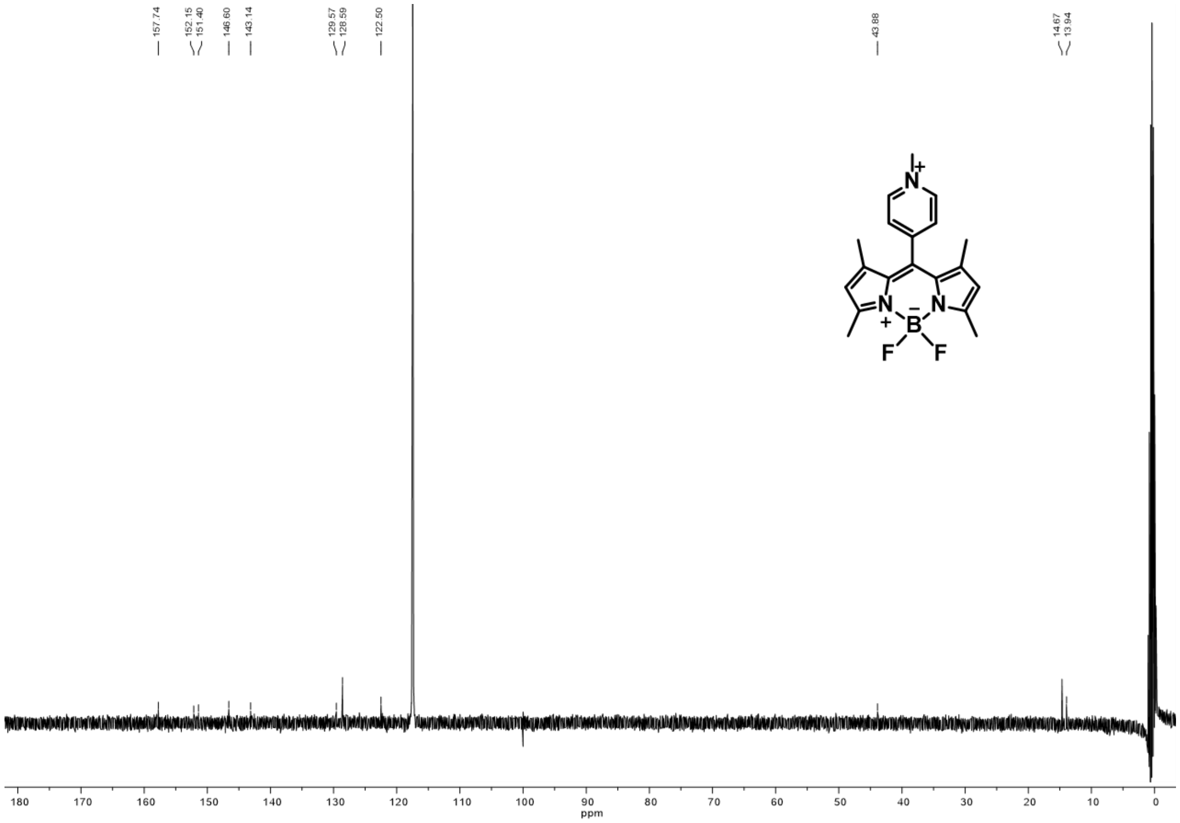


**Figure S29**. $^{13}C\{^{1}H\}$ NMR (101 MHz, $CD_3CN$, 298 K) of **3b.**

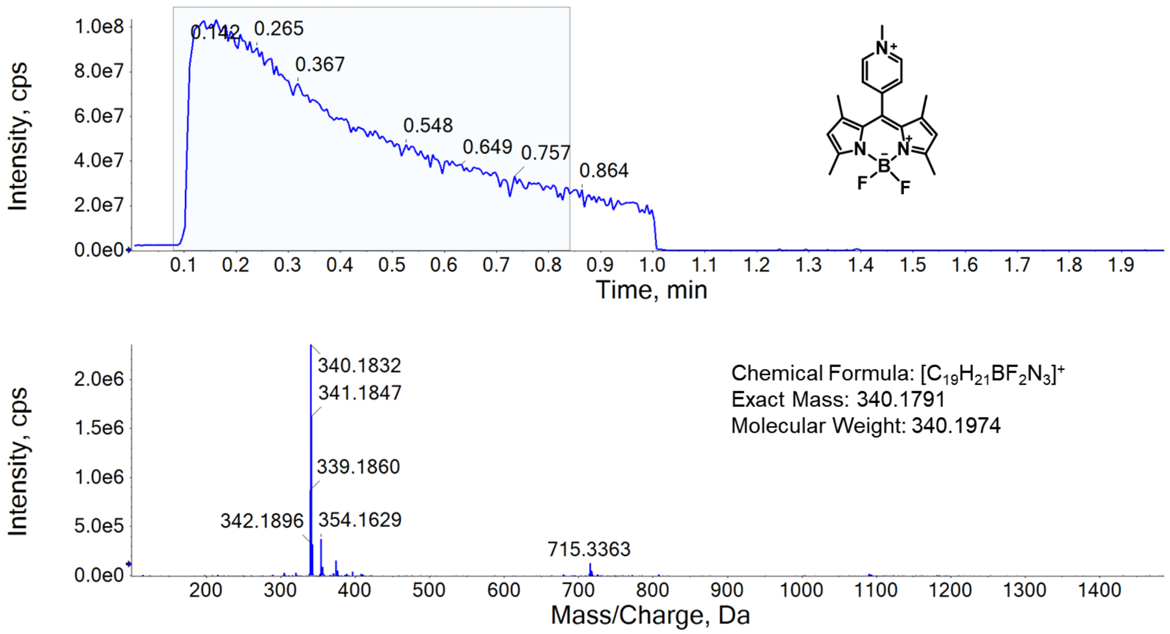


**Figure S30.** HRMS (+ve ESI) spectrum of **3b**.

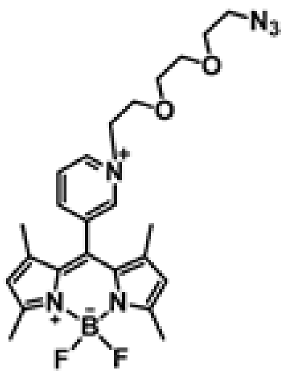

**Synthesis of 1b-azide:**

**1a** (95.3 mg, 0.293 mmol) was added to the solution of azide-PEG3-iodide (Lumiprobe, CAS No.: 1309457-01-9, 215 µL, 0.65 mmol) in 10 mL of MeCN. The mixture was heated at 80 ℃ overnight. The mixture was concentrated by rotary evaporator and the product was purified by neutral aluminium oxide chromatography using $CH_2Cl_2$/MeOH (*v/v* = 10:1) to give a red solid. Yield: 153 mg, 86 %. $^1$H NMR (400 MHz, $CD_3CN$, 298 K) δ 9.07 (d, *J* = 6.2 Hz, 1H), 8.97 (s, 1H), 8.66 (d, *J* = 8.1 Hz, 1H), 8.27 (dd, *J* = 7.9, 6.3 Hz, 1H), 6.19 (s, 2H), 4.94 – 4.76 (m, 2H), 4.07 – 3.89 (m, 2H), 3.64 – 3.55 (m, 2H), 3.52 – 3.40 (m, 4H), 3.30 – 3.20 (m, 2H), 2.51 (s, 6H), 1.43 (s, 6H). $^{13}$C NMR (101 MHz, $CD_3CN$, 298 K) δ 158.91, 147.68, 146.99, 145.21, 144.18, 135.69, 132.37, 131.96, 130.05, 123.51, 71.19, 70.84, 70.52, 69.30, 62.92, 51.25, 15.75, 14.87. HRMS (+ve ESI): calculated for $C_{24}H_{30}BF_2N_6O_2$ $[M]^+$ *m/z* 483.2486, found 483.2472.

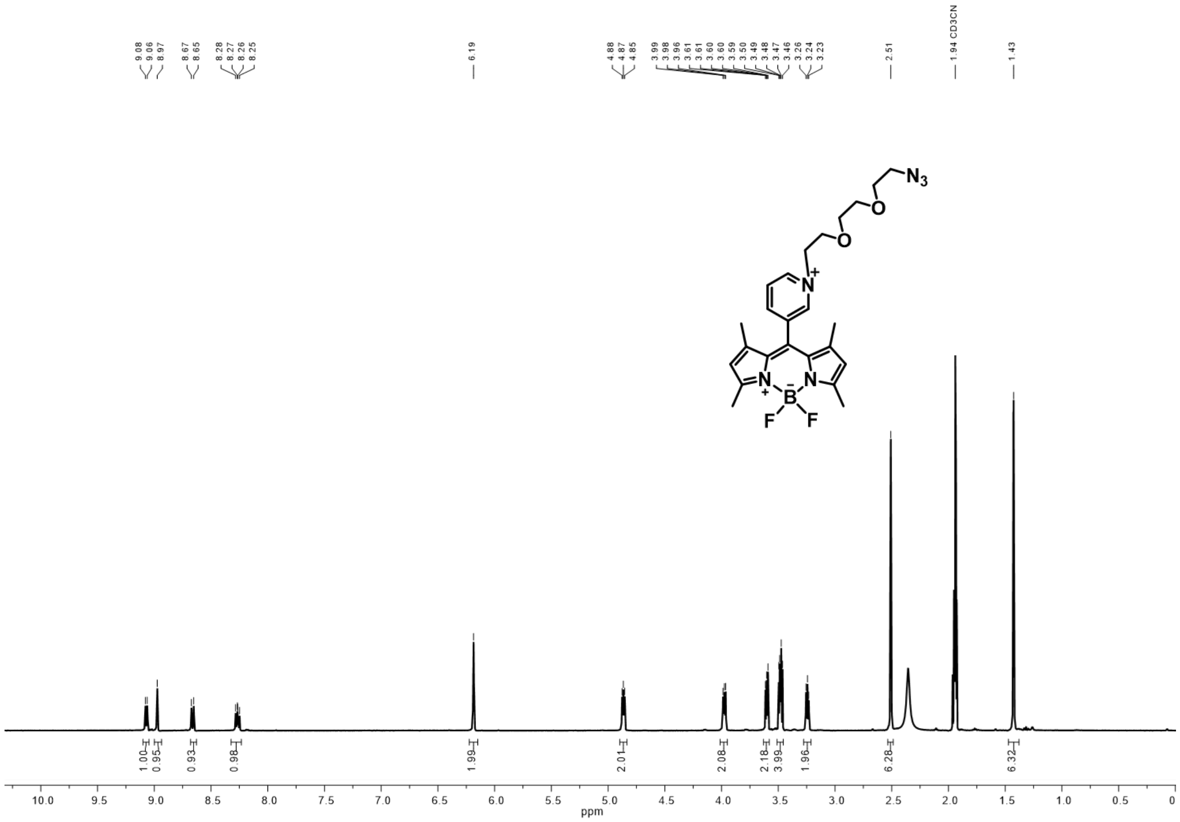


**Figure S31.** $^1$H NMR spectrum (400 MHz, $CD_3CN$, 298 K) of **1b-azide**.

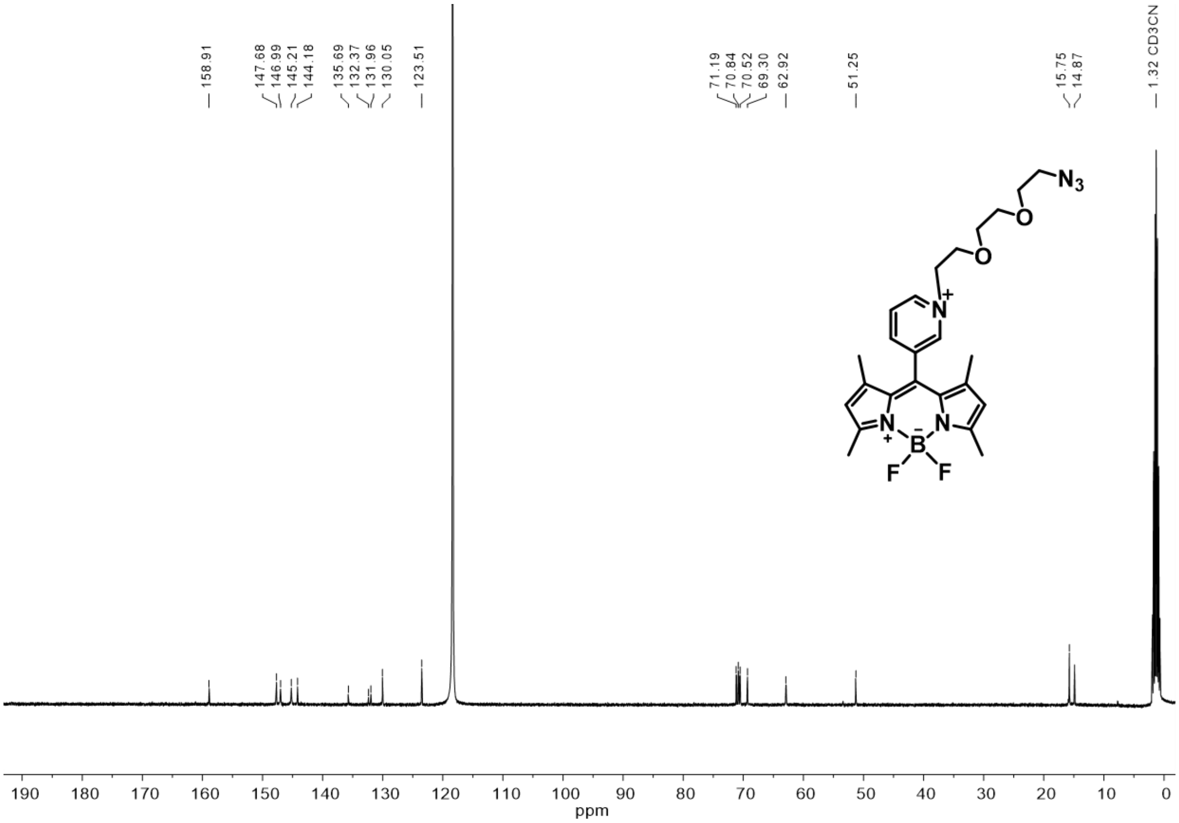


**Figure S32**. $^{13}C\{^{1}H\}$ NMR (101 MHz, $CD_3CN$, 298 K) of **1b-azide**.

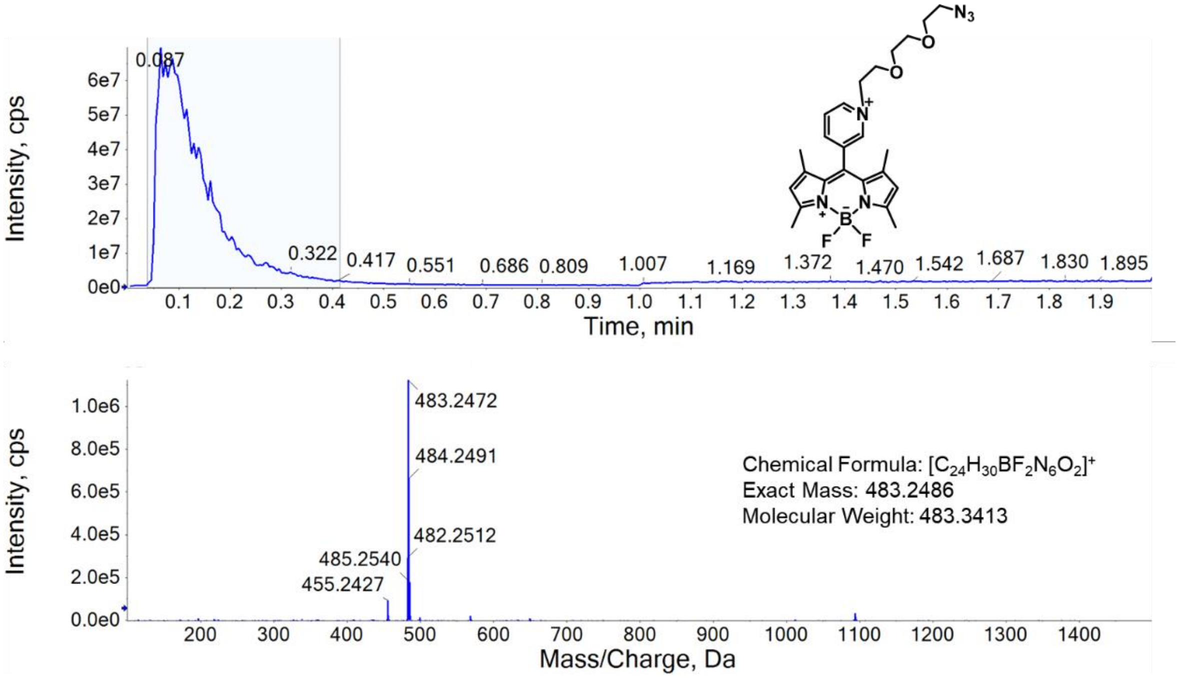


**Figure S33.** HRMS (+ve ESI) spectrum of **1b-azide**.

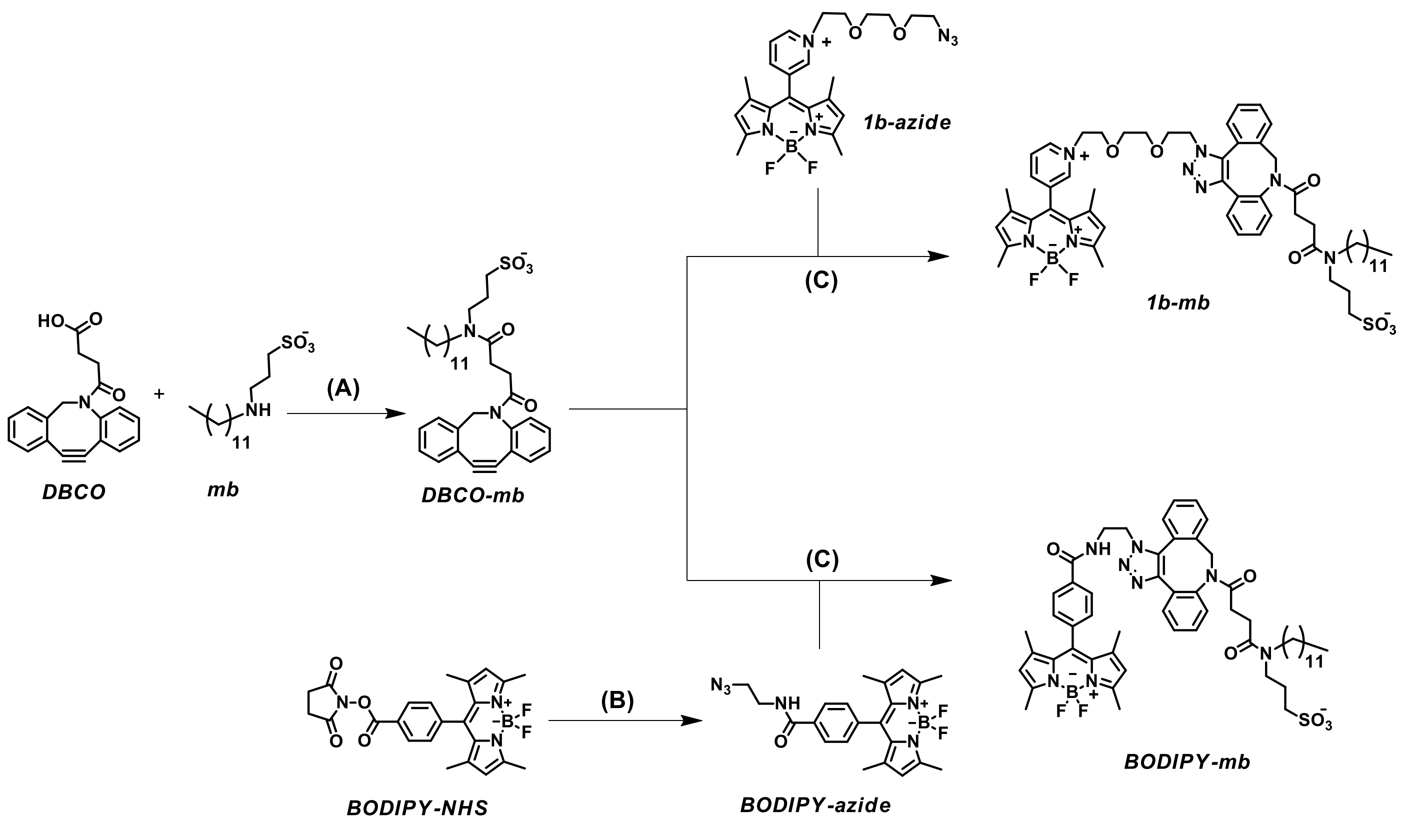


**Scheme S2.** Synthesis of membrane targeting ligand **DBCO-mb** and its conjugation to fluorophores to form **1b-mb** and **BODIPY-mb**. (**A**) HATU, DIPEA, DMF, room temperature, 3 h; (**B**) 2-Azidoethylamine hydrochloride, triethylamine, $CH_2Cl_2$, room temperature, overnight; (**C**) $CH_2Cl_2$, room temperature, 20 min.

**Synthesis of DBCO-mb:**

**DBCO**[3] and **mb**[4] were synthesized using the literature procedure. **DBCO** (100 mg, 0.3278 mmol. 1 equiv.) was dissolved in dry DMF (3 mL), then DIPEA (114 µL, 0.66 mmol, 2 equiv.) was added and stirred for 5 min. Next, HATU (131 mg, 0.344 mmol, 1.05 equiv.) was added and stirred for another 10 min. A solution of **mb** (106 mg, 1.05 mmol, 1.05 equiv.) and DIPEA (114 µL, 0.6556 mmol, 2 equiv.) in DMF (3 mL) was added and stirred for 3 h. Reaction progress was monitored by TLC. After completion of the reaction, DMF was removed by rotary evaporator, and the crude compound was purified by silica gel column chromatography using (9:1 DCM/MeOH) to obtain 40 mg of **DBCO-mb** as a pale-yellow solid. Yield: 40 mg, 67.4 µmol, 10%. HRMS (+ve ESI): calculated for $C_{34}H_{47}N_2O_5S$ $[M+H]^+$ *m/z* 595.3200, found: 595.3176.

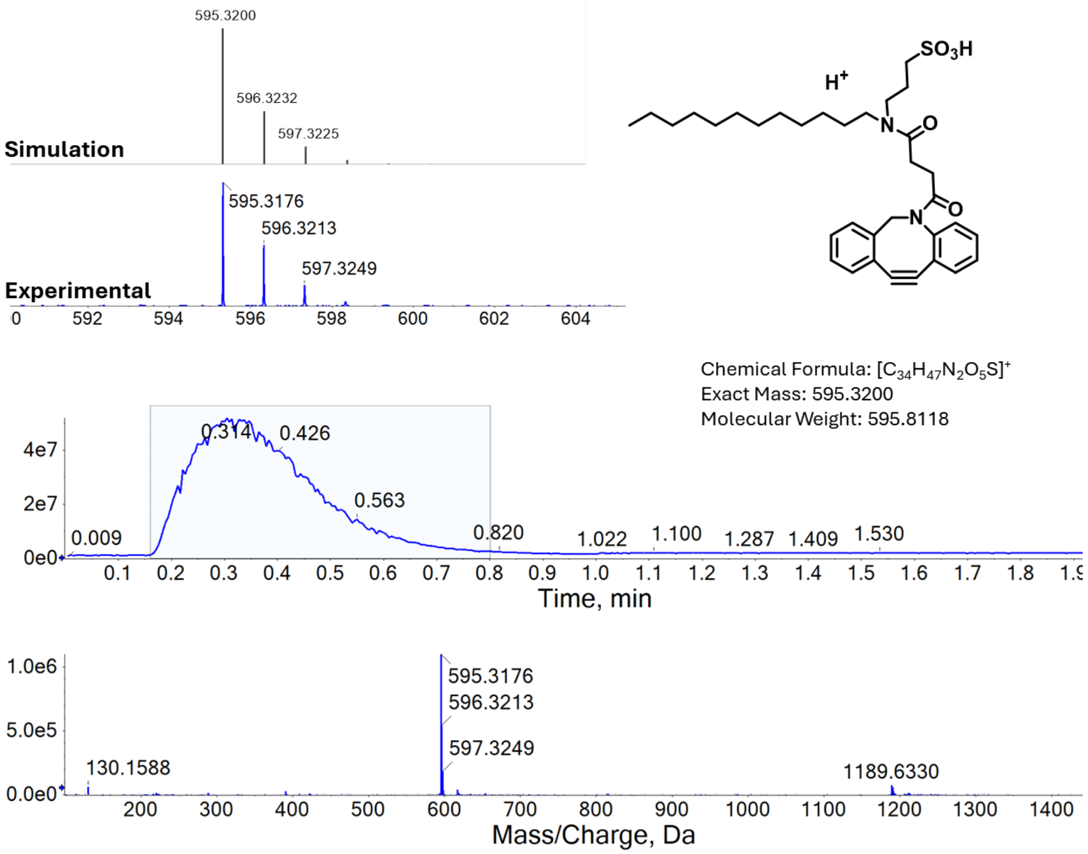


**Figure S34**. HRMS (+ve ESI) spectrum of **DBCO-mb**.

**Synthesis of BODIPY-azide:**

**BODIPY-NHS**[5] (93.2 mg, 0.2 mmol) was dissolved in 5 mL of $CH_2Cl_2$. 2-Azidoethylamine hydrochloride (155 mg, 1.26 mmol, 6 equiv.) was dissolved in 5 mL of $CH_2Cl_2$ with triethylamine (200 μL, 1.44 mmol, 6 equiv.). The two solutions were added together and stirred overnight at room temperature. The organic layer was washed with $H_2O$ three times. The organic layer was concentrated by rotary evaporator and the product was purified by silica gel column chromatography using ethyl acetate/hexanes (*v/v* = 1:1) to give **BODIPY-azide** as a red solid. Yield: 44 mg, 50%. $^1H$ NMR (400 MHz, $CDCl_3$, 298 K) δ 7.93 (d, *J* = 8.2 Hz, 2H), 7.40 (d, *J* = 8.3 Hz, 2H), 5.99 (s, 2H), 3.73 – 3.65 (m, 2H), 3.64 – 3.59 (m, 2H), 2.56 (s, 6H), 1.35 (s, 6H). $^{13}C$ NMR (101 MHz, $CDCl_3$, 298 K) δ 166.91, 156.15, 143.03, 138.82, 134.64, 129.92, 128.73, 128.48, 127.97, 121.64, 51.02, 39.66, 14.76. HRMS (+ve ESI): calculated for $C_{22}H_{24}BF_2N_6O$ $[M+H]^+$ *m/z* 437.2067, found 437.2056.

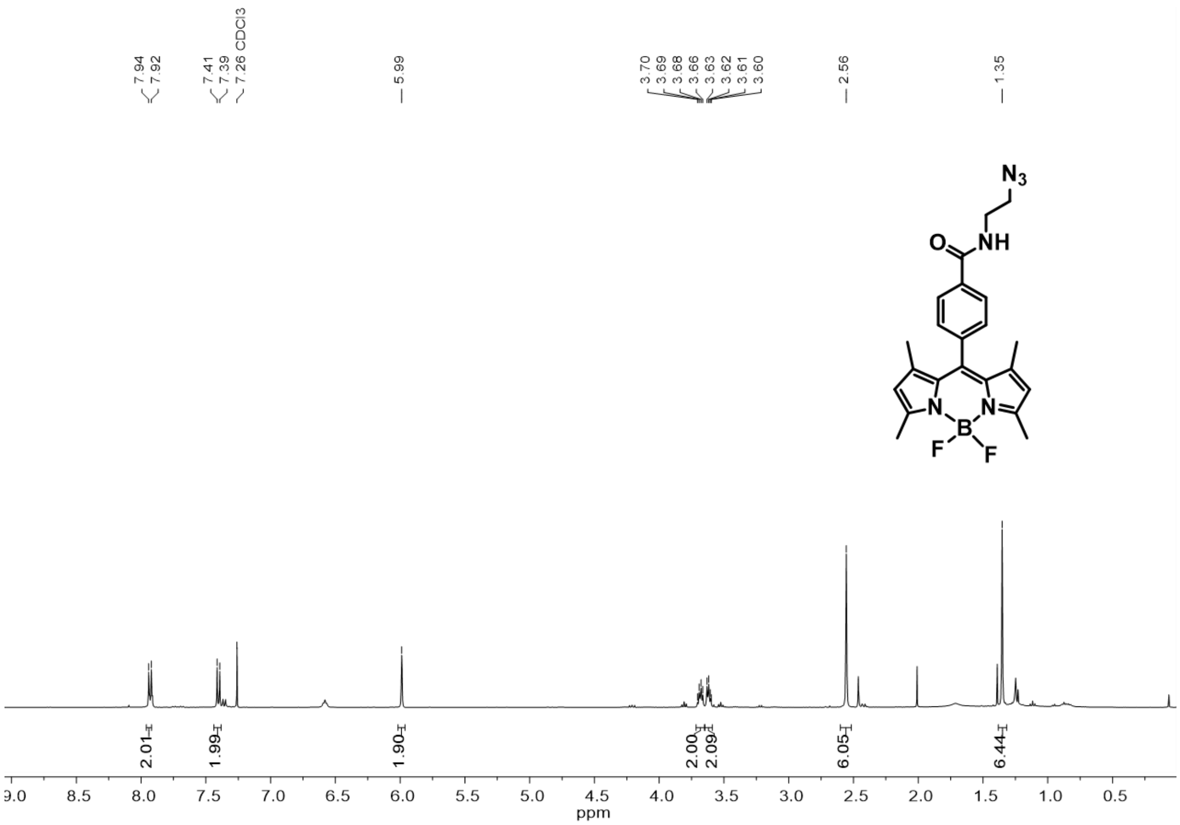


**Figure S35.** $^1H$ NMR spectrum (400 MHz, $CDCl_3$, 298 K) of **BODIPY-azide**.

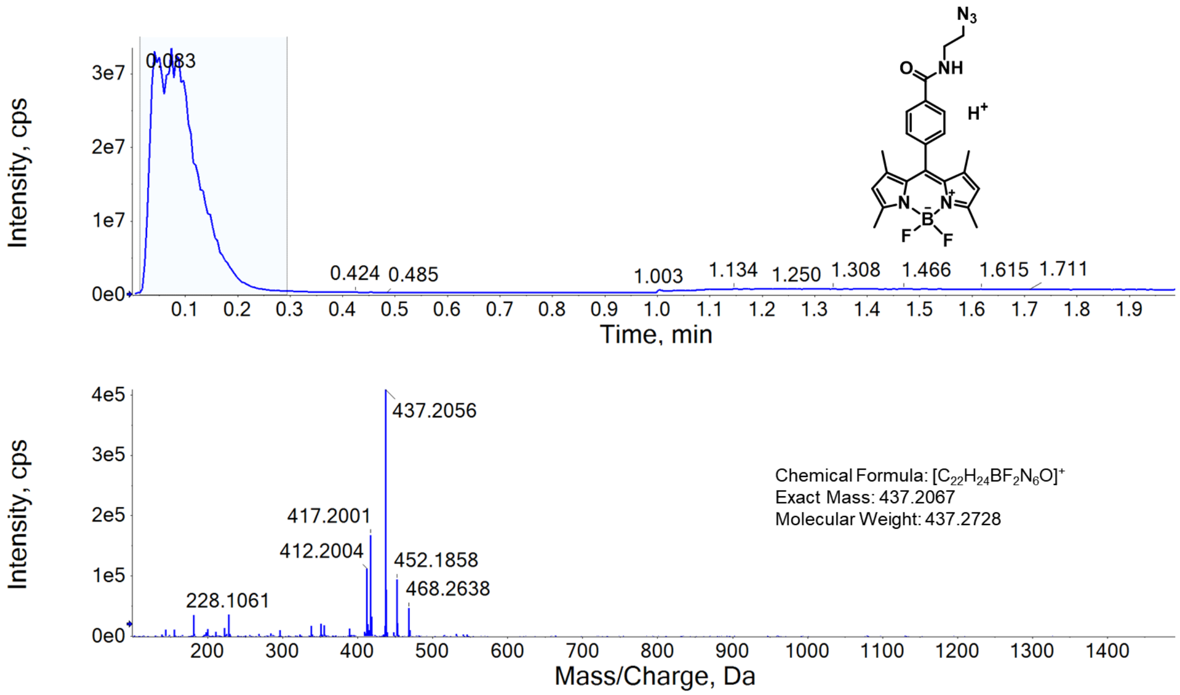


**Figure S36**. HRMS (+ve ESI) spectrum of **BODIPY-azide**.

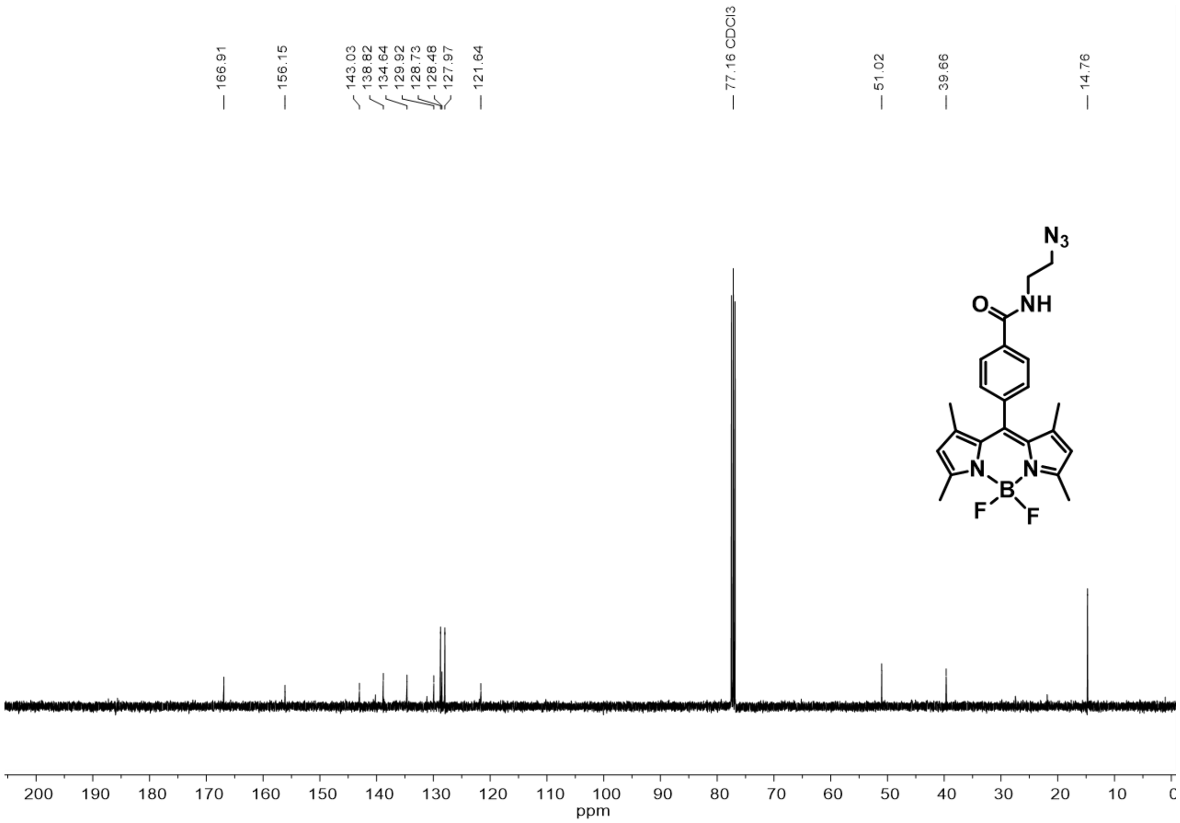


**Figure S37.** $^{13}C\{^{1}H\}$ NMR (101 MHz, $CDCl_3$, 298 K) of **BODIPY-azide**.

**Synthesis of BODIPY-mb:**

**DBCO-mb** (5 mg, 1 equiv.) was dissolved in 200 µL of $CH_2Cl_2$, **BODIPY-azide** (1 equiv.) was added and stirred for 15 min at rt, reaction progress was monitored by TLC, and after completion of the reaction, the crude product was purified by preparative TLC to obtain 3 mg of **BODIPY-mb** as an orange solid. Yield: 3 mg, 8.42 µmol, 35%. HRMS (+ve ESI): calculated for $C_{56}H_{70}BF_2N_8O_6S$ $[M+H]^+$ *m/z* 1031.5195, found: 1031.5230; calculated for $C_{56}H_{69}BFN_8O_6S^+$ $[M–F]^+$ *m/z* 1011.5138, found: 1011.5131.

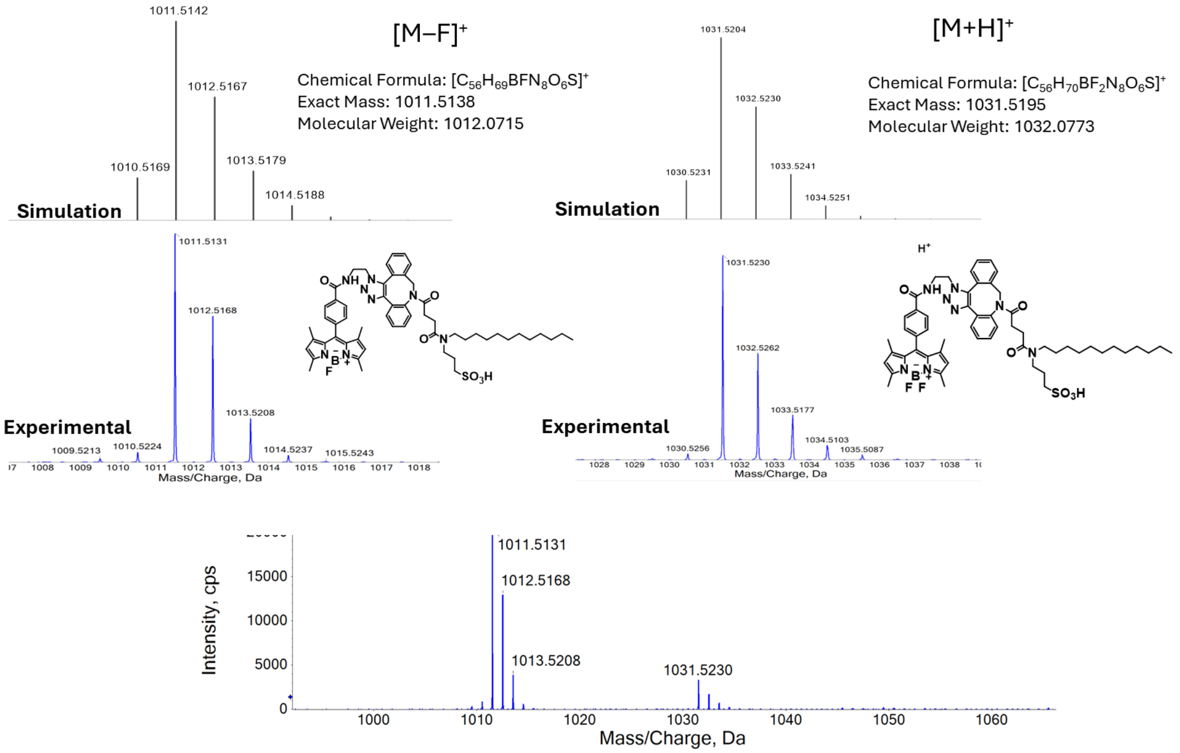


**Figure S38**. HRMS (+ve ESI) spectrum of **BODIPY-mb**.

**Synthesis of 1b-mb:**

**DBCO-mb** (3 mg, 1 equiv.) was dissolved in 200 µL of $CH_2Cl_2$, **1b-azide** (1 equiv.) was added and stirred for 15 min at rt, reaction progress was monitored by TLC, and after completion of the reaction, the crude product was purified by preparative TLC to obtain 3 mg of **1b-mb** as an orange solid. Yield: 3 mg, 2.79 µmol, 52%. HRMS (+ve ESI) calculated for $C_{58}H_{76}BF_2N_8O_7S^+$ $[M+H]^+$ *m/z* 1077.5613, found: 1077.5585.

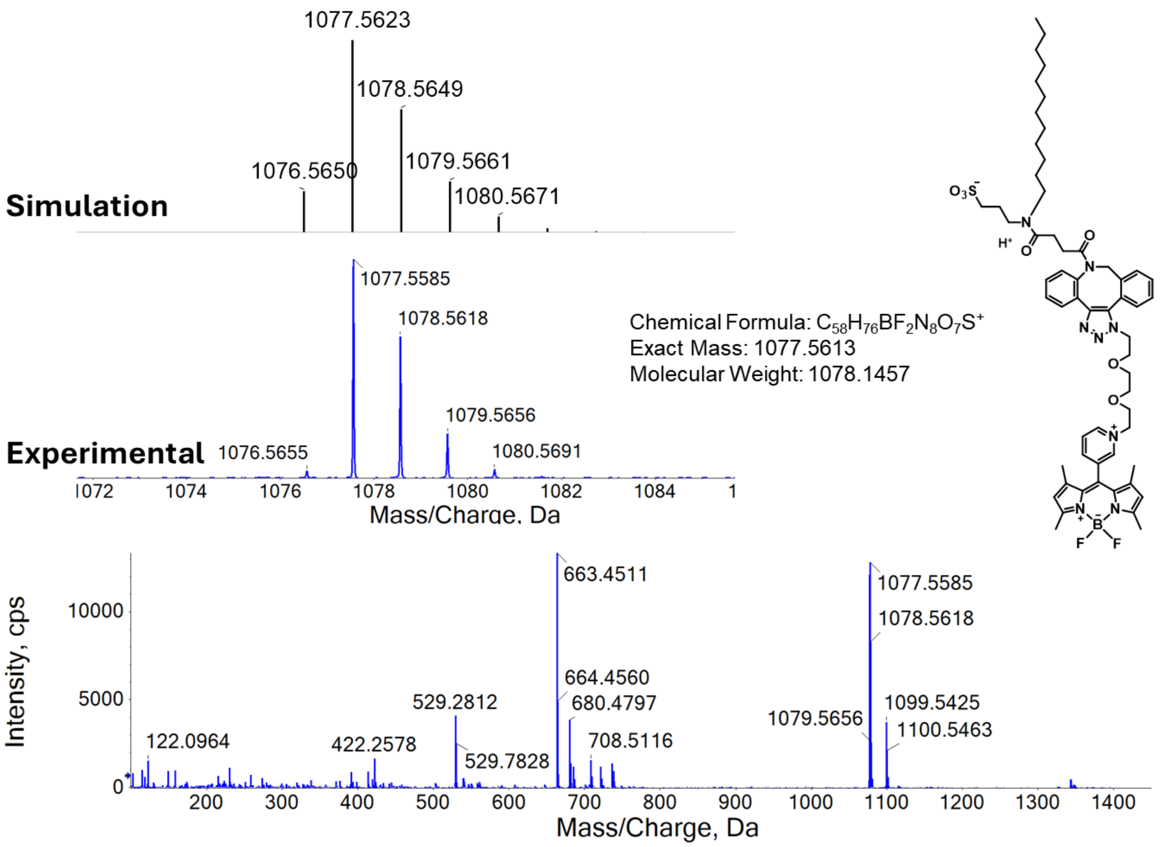


**Figure S39**. HRMS (+ve ESI) spectrum of **1b-mb**.

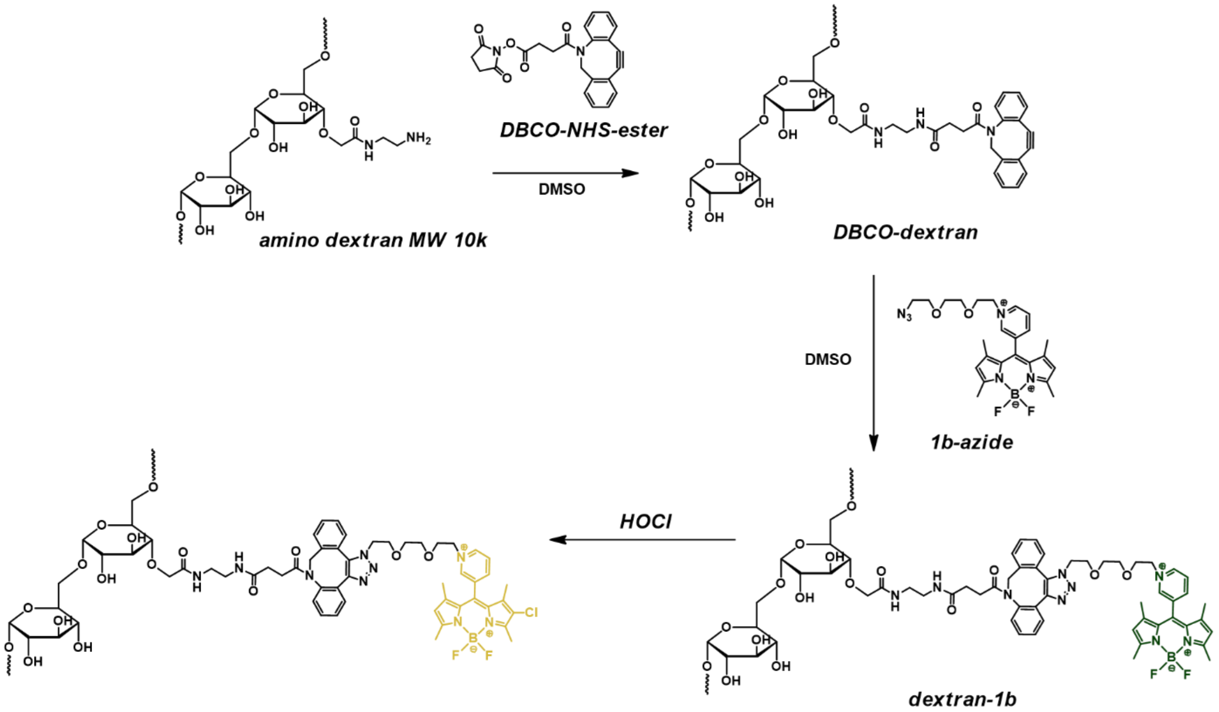


**Scheme S3.** Synthetic **dextran-1b**.

**Synthesis of dextran-1b:**

In a 10 mL glass vial, 55 mg of amino dextran (FinaBio, AD10X10, 10kDa) was dissolved in 5 mL of anhydrous DMSO. Then DBCO-NHS-ester (5 mg, ~0.2 mol. equiv. to $–NH_2$ groups) was added and the mixture was allowed to stir at room temperature for three days. The mixture was diluted with water and dialyzed to remove the DMSO (Thermo Scientific, SnakeSkinTM Dialysis Tubing with MWCO of 3500). The resulting solution was lyophilized to give **DBCO-dextran** as a white solid. Yield: 28.2 mg, 51 % based on mass recovery (Step 1).

Next, 14 mg of the **DBCO-dextran** was dissolved in 5 mL of anhydrous DMSO, then **1b-azide** (2 mg, ~0.2 mol. equiv. to $–NH_2$ groups) was added and the mixture was stirred at room temperature for three days. The mixture was diluted with water and dialyzed to remove the DMSO (Thermo Scientific, SnakeSkinTM Dialysis Tubing with MWCO of 3500). The resulting solution was lyophilized to give **dextran-1b** as a purple solid. Yield: 6.9 mg, 49 % based on mass recovery (Step 2).
